# Co-transcriptional folding orchestrates sequential multi-effector sensing by a glycine tandem riboswitch

**DOI:** 10.1101/2025.05.28.656632

**Authors:** Rosa A. Romero, Adrien Chauvier, Serena S. Teh, Vincent A. Reed, Sicheng Zhang, Courtney E. Szyjka, Shi-Jie Chen, Eric J. Strobel, Nils G. Walter

## Abstract

Riboswitches are non-coding RNA motifs that regulate gene expression in response to ligand binding. The glycine tandem riboswitch (GTR) is notable because it comprises two distinct glycine aptamers that interact extensively. These inter-aptamer contacts drive conformational changes in the downstream expression platform to control gene expression. Despite extensive studies, the role of glycine and RNA folding pathways in co-transcriptional regulation remains unclear. Here, we integrate single-molecule kinetic analysis, co-transcriptional RNA structure probing, and computational modeling to reveal that the GTR processes multiple molecular inputs sequentially, guided by polymerase pausing. Our findings elucidate its stepwise 5’-to-3’ folding pathway and demonstrate how sequential glycine binding to each aptamer, K^+^ binding to a kink-turn, non-native RNA folding intermediates, inter-aptamer docking that drives binding site pre-organization, and modulation by the transcription factor NusA collectively orchestrate co-transcriptional gene regulation. These results support a model in which glycine binding cooperativity arises through non-equilibrium mechanisms rather than a classical concerted model.

## INTRODUCTION

In all domains of life, gene regulation by non-coding RNAs is coupled to transcription^1^. As RNA polymerase (RNAP) synthesizes RNA, the nascent transcript begins to fold into functional structures in real-time^1^. This process, known as co-transcriptional folding, is crucial for gene expression control mechanisms^2,3^. This dynamic, temporally coordinated folding landscape is shaped by intrinsic and auxiliary factors that collectively allow cells to adapt to fluctuating environmental cues. Notably, site-specific RNAP pausing and co-transcriptional RNA-protein interactions have been shown to reciprocally influence RNA folding and function^2,4^. Transcription pauses are pervasive across all domains of life and serve as critical checkpoints for eukaryotic pre-mRNA splicing, prokaryotic transcription-translation coupling, and the control of gene expression^5–7^.

Riboswitches are cis-regulatory structural motifs that are typically found in the 5’ untranslated region of bacterial mRNAs^8,9^. In a typical riboswitch-mediated gene regulation mechanism, a specific ligand binds to a pocket in an aptamer domain and induces local and global structural rearrangements in the downstream expression platform that control gene expression, typically at the level of transcription or translation. Although the aptamer domains are highly conserved^10^, emerging evidence suggests that ligand recognition and the functional outcome of riboswitch folding depend on the transient accessibility of conformational isomers during transcription, defining a time window for the ligand to affect gene regulation^11–18^. In transcriptional riboswitches, ligand binding must kinetically trap a regulatory conformation before the RNAP reaches the termination point^19,20^.

Among riboswitch families, a subset harbors two tandem aptamers that can recognize either identical or distinct ligands^21–23^. The glycine tandem riboswitch (GTR) is a well-characterized example in which both aptamers sense glycine^24^. Other tandem aptamer riboswitches can function as Boolean logic gates, which have been used in bioengineering, by integrating dual inputs to control gene expression output^22,25^. While GTRs can act as either ON or OFF switches depending on downstream gene context^26^, the majority (∼81%) function as ON switches in glycine catabolism pathways^27^. Notably, in this class aptamer 1 typically has higher GC content and binding affinity than the downstream aptamer 2^26^, suggesting an ordered mechanism for cooperative glycine sensing. However, the discovery of an upstream kink-turn motif^28,29^ that disrupts glycine-binding cooperativity has challenged prevailing models^27,28^.

Recent structural and mutational studies have revealed tertiary inter-aptamer interactions―specifically the α and β A-minor motifs and a Hoogsteen base pair termed γ (Figure 1A)―that help organize the riboswitch in both ligand-free and bound states^30,31^. However, the mechanistic contributions of each aptamer and the kink-turn motif to glycine binding and gene regulation remain unresolved. The GTR from *Bacillus subtilis* (*Bsu*-GTR), in particular, appears to deploy a stepwise binding mechanism, where glycine and K⁺ first stabilize aptamer 1 and the kink-turn, priming the structure for interactions with aptamer 2 later in the folding pathway^27,32^. Binding of the kink-turn motif by the protein YbxF was shown to enhance glycine binding^33^, while the transcription factor NusA is known to prolong transcription pauses that support transcriptional regulation^13,20^―underscoring the role of auxiliary cellular factors.

**Figure 1.**
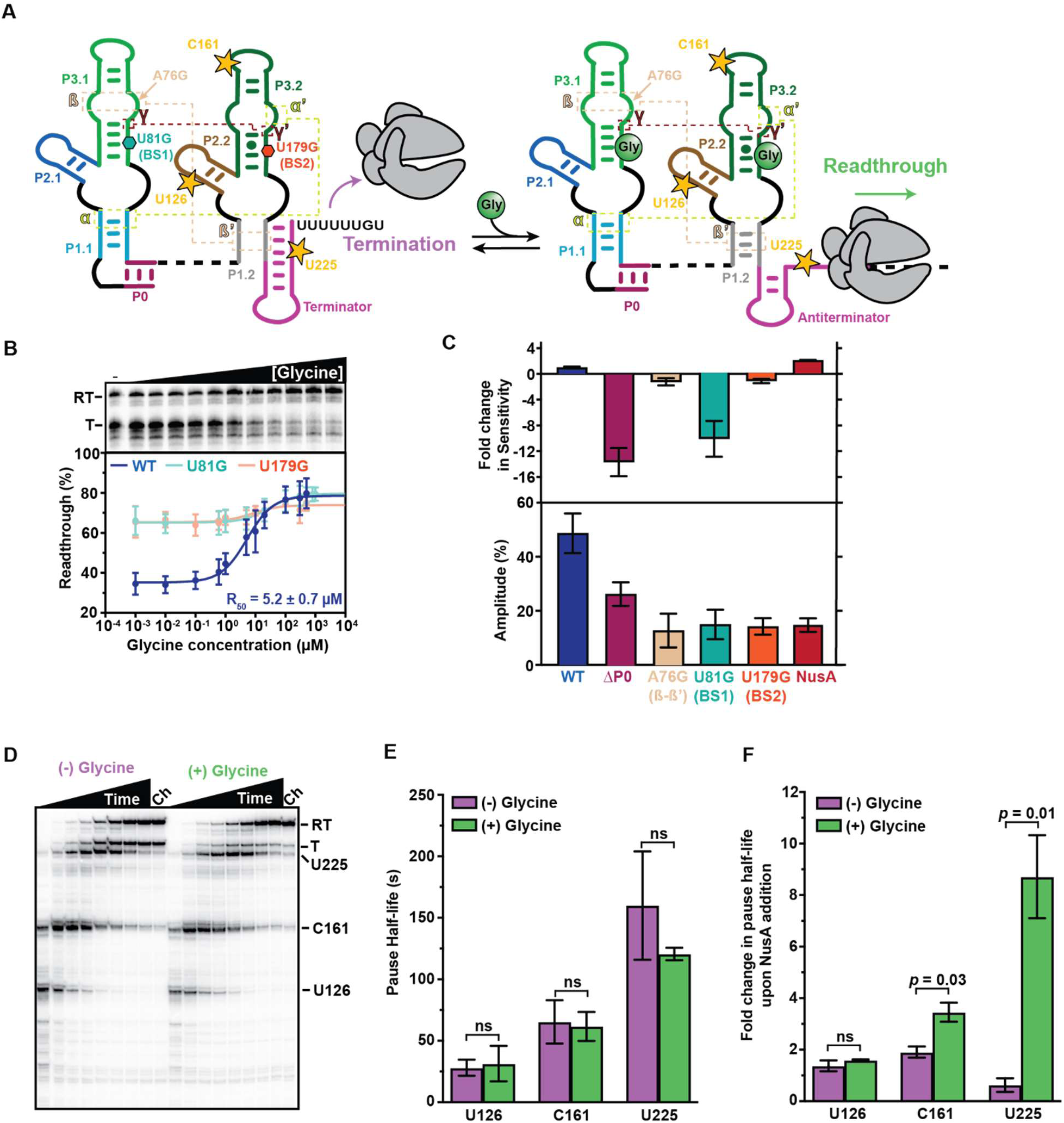
Co-transcriptional folding and pausing determine gene regulation. (**A**) The *Bsu*-GTR regulates gene expression using a glycine binding-mediated transcription antitermination mechanism. Long-range tertiary interactions (α, β, and γ) and the positions of RNAP pause sites (yellow stars) are indicated. (**B**) Glycine dose-response curves for the WT *Bsu*-GTR, U81G variant, and U179G variant measured by single-round *in vitro* transcription. A representative gel for the WT riboswitch is shown. (**C**) Top: Fold change in sensitivity relative to the WT construct for each variant. Bottom: Amplitude of the glycine-mediated transcription antitermination response. (**D**) Time-resolved single-round *in vitro* transcription of the *B. subtilis* glycine riboswitch in the presence and absence of glycine. (**E**) Quantification of pause half-lives. (F) Fold change of pause half-lives in the presence of 100 nM NusA. All error bars are the SD of the mean from independent replicates (n=3). RT, Readthrough; T, Terminated.

Here, we integrate single-molecule fluorescence microscopy, co-transcriptional RNA structure probing, and computational modeling to investigate the folding and regulatory mechanisms of the *Bsu-*GTR. We find that transcription is punctuated by strategically positioned pauses that coordinate the binding of K^+^ and two glycine molecules with transcription. Binding of the first glycine and K⁺ pre-organizes aptamer 1 for the α, β and linchpin γ interactions, facilitating aptamer 2 docking. The final pause, which is embedded in the transcription terminator, extends the window for formation of the inter-aptamer contacts and for glycine binding to aptamer 2, and facilitates a secondary path for glycine binding in which inter-aptamer contacts pre-organize both binding sites to promote glycine binding ahead of terminator hairpin folding. The pauses also help recruit NusA, promoting structural transitions that increase regulatory efficiency. Our findings support a non-equilibrium model of cooperativity―distinct from the classical Monod-Wyman-Changeux paradigm―in which kinetic partitioning of folding intermediates defines ligand sensing and regulatory function. We propose that such kinetically driven cooperativity is a general principle guiding the integration of multiple cellular factors that can interact during co-transcriptional processes.

## RESULTS

### Co-transcriptional folding and pausing determine gene regulation

The *Bsu*-GTR regulates expression of the *gcvT* operon, which encodes proteins of the glycine cleavage system, using a glycine-mediated transcription antitermination mechanism^24^ (Figure 1A). To assess glycine riboswitch function under standard *in vitro* conditions, we performed single-round transcription of the riboswitch using *E. coli* RNAP across a range of glycine concentrations (Figure 1B). The glycine concentration required to achieve half-maximal readthrough (R_50_) by the wild-type (WT) *Bsu*-GTR was 5.2 ± 0.7 µM, which suggests nearly tenfold tighter binding than previously reported^27^. This difference may be due to the inclusion of an extended leader sequence in the previous study^27^, which is likely to influence RNA folding. The difference in readthrough amplitude we observed between the no-ligand and glycine-saturated conditions was significant at 43.4 ± 1.2% (Figure 1B and Table S2).

Variants that disrupt either binding site 1 (U81G) or binding site 2 (U179G) showed little to no glycine-mediated transcription antitermination (Figure 1B), indicating that ligand engagement at both aptamers is required for effective gene regulation. Notably, disruption of binding site 1 substantially impaired glycine sensitivity, shifting the R_50_ from 5.2 ± 0.7 to 28.6 ± 3.8 µM. In contrast, mutation of binding site 2 primarily reduced the amplitude of the readthrough response from 43.4 ± 1.2 to 8.4 ± 1.0% (Figure 1C and Table S2). These observations align with prior findings that suggest distinct functional roles for the two aptamer domains, with aptamer 1 in the *Bsu*-GTR possessing a thermodynamically more stable binding site and accordingly exhibiting higher glycine affinity^26,27,34^.

To assess how riboswitch conformational features influence transcription antitermination, we introduced sequence perturbations that either eliminate P0 (ΔP0) or weaken the β A-minor motif (A76G) (Figures 1A and S1A). Removing P0 caused a substantial, 12-fold reduction in glycine sensitivity, indicating that P0 folding is essential for efficient co-transcriptional ligand binding (Figures 1C, S1B and Table S2). To further test the role of P0 stabilization in glycine binding, we performed *in vitro* transcription assays in the presence of *Bsu*-YbxF, a protein known to bind the kink-turn and promote its bent conformation^33,35^. Consistent with its expected stabilizing role, *Bsu*-YbxF enhanced the glycine sensitivity of the WT *Bsu*-GTR by 4-fold and broadly increased transcription readthrough (Figure S1C and Table S2). In contrast, *Bsu*-YbxF had no significant effect on the sensitivity or readthrough amplitude of the ΔP0 variant (Figure S1D and Table S2), further reinforcing the notion that P0 is integral to establishing the global tandem aptamer architecture. Conversely, weakening the β A-minor motif using an A76G mutation only had a modest effect on glycine sensitivity but significantly impaired the glycine-mediated antitermination efficiency (Figures 1C, S1E and Table S2), highlighting that the β-β’ tertiary interaction is critical for *Bsu*-GTR function.

We next mapped transcriptional pause sites using time-resolved *in vitro* transcription (Figure 1D). All identified pauses sites were located within the aptamer 2 region, and glycine addition had no significant impact on their lifetimes (Figures 1A, 1E and Table S3). Notably, each of the three pauses conformed to the established consensus pause sequence^36,37^ (Figure S2A). To examine potential regulatory involvement of transcriptional pausing, we tested the effect of NusA, a transcription elongation factor known to interact with RNAP and stabilize pause events^38,39^. In the presence of glycine, NusA significantly prolonged the lifetime of the C161 and U225 pauses (Figures 1F, S1G and Table S3), suggesting that it contributes to the regulatory function of the *Bsu-*GTR. Supporting this interpretation, NusA also enhanced the glycine sensitivity of the WT *Bsu*-GTR by ∼2-fold and promoted transcription readthrough overall (Figures 1C, S1F and Table S2).

Together, our results invoke a model in which transcriptional pauses act as checkpoints that coordinate co-transcriptional aptamer folding with ligand sensing to drive effective gene regulation. The formation and stabilization of the kink-turn motif, along with NusA-mediated pause enhancement, likely contribute to the precise temporal orchestration of riboswitch function, ensuring that structural transitions are properly aligned with transcriptional progression to enable a robust gene regulatory response.

### Glycine enhances NusA interaction and *Bsu*-GTR folding at PEC225

To further dissect the dynamics of the NusA interaction with paused elongation complexes (PEC) at sites C161 and U225, we conducted single-molecule colocalization binding assays using fluorescently labeled PECs and NusA^20^. RNAP was stalled at these sites (PEC-161 and PEC-225) using biotin-streptavidin roadblocks, and NusA binding and dissociation kinetics were determined at the single-molecule level (Figure 2A and S2B). As observed previously^13,20^, NusA transiently binds to PECs, allowing extraction of bound (τ_bound_) and unbound (τ_unbound_) lifetimes at both PEC161 (Figures 2B and S3A, B) and PEC-225 (Figure 2A, D and S3C). Data were fit with single- or double-exponential functions to derive dissociation (*k*_NusA-off_) and association (*k*_NusA-on_) rate constants, respectively (Figures 2C, E and S3D-G).

**Figure 2.**
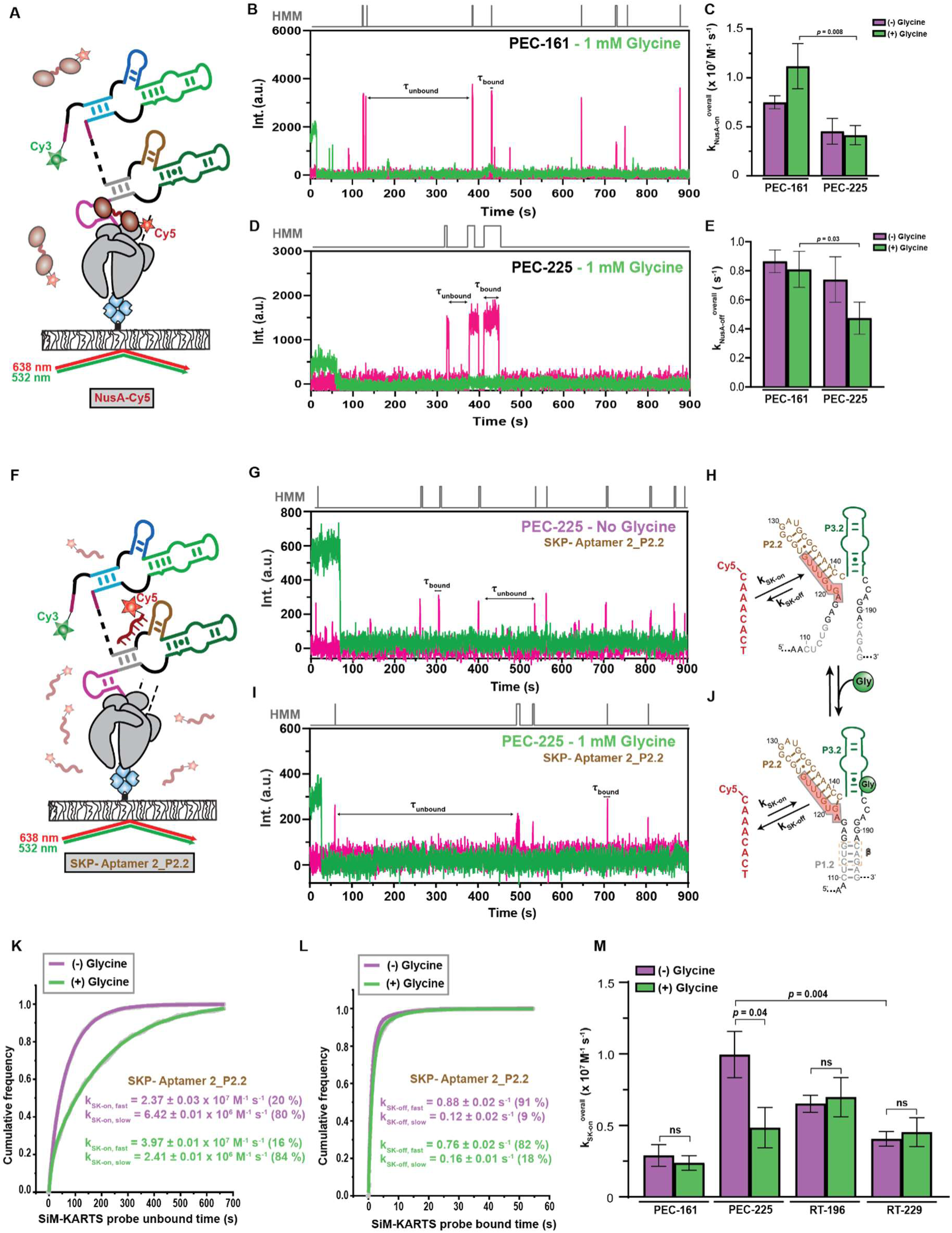
Glycine enhances NusA interaction and *Bsu*-GTR folding at PEC225. (**A**) Single-molecule setup for a NusA colocalization study conducted using NusA-Cy5 and PEC-Cy3. The PEC is immobilized by a biotin-streptavidin roadblock. Binding of NusA-Cy5 to the PEC is monitored through direct excitation of Cy5. (**B**, **D**) Representative single-molecule trajectories showing NusA-Cy5 binding (pink) to PEC-161 (B) and PEC-225 (D) in the presence of 1 mM glycine. (**C**, **E**) Binding rate constants *k*_on_ (C) and *k*_off_ (E) of NusA-Cy5 in the context of PEC-225 and PEC-161 in the absence and presence of 1 mM glycine. (**F**) SiM-KARTS experimental setup. (**G-J**) Binding of the SiM-KARTS probe targeting the β-β’ interaction in aptamer 2 is monitored through direct excitation of the Cy5 fluorescent dye in the absence and presence of glycine. Representative single-molecule trajectories showing the SiM-KARTS probe binding (pink) to P2.2 of PEC-225 recorded in the absence (G) or presence (I) of 1 mM glycine. Hidden Markov modeling (HMM) is shown above of each trace. (**K**, **L**) Plots showing the cumulative unbound (K) and bound (L) dwell times of the SiM-KARTS probe in the absence and presence of 1 mM glycine in the context of PEC-225. The association (*k*_on_) and dissociation (*k*_off_) rate constants of the SiM-KARTS probe are indicated. The reported errors are the error of the fit. The total number of molecules analyzed for each condition is: (−) Glycine = 312; (+) Glycine = 348. (M) Overall binding rate constants (*k*_on_) of the SiM-KARTS probe in the context of PEC-161, PEC-225, RT-196 and RT-229 constructs determined in the absence (purple) and presence (green) of 1 mM glycine. Error bars are the SD of the mean of independent replicates. The statistical significance of differences was determined using the two-tailed Student’s t-test. a.u., arbitrary unit.

While glycine binding had no significant effect on either *k_NusA-on_* or *k_NusA-off_* at PEC-161 or PEC-225 (Figure 2C,E), side-by-side comparison in the presence of glycine revealed that NusA was recruited more rapidly to PEC-161 than PEC-225 (Figure 2A-E). Specifically, at PEC-161, NusA exhibited two distinct association rate constants: 23.4 ± 0.14 × 10^6^ M^−1^ s^−1^ (*k_NusA-on,fast_*; fraction 45%) and 2.29 ± 0.01 × 10^6^ M^−1^ s^−1^ (*k_NusA-on,slow_*; fraction 55%), resulting in a weighted average rate constant (*k_NusA-on_*) of 11.2 ± 2.31× 10^6^ M^−1^ s^−1^ (Figures 2C, S3D and Table S4). In contrast, at PEC-225, the average rate constant was ∼3-fold slower at *k_NusA-on_* = 4.16± 0.98 × 10^6^ M^−1^ s^−1^ (*k_NusA-_*_on,fast_ = 7.26 ± 0.04 × 10^6^ M^−1^ s^−1^; *k_NusA-_*_on,slow_ = 1.38 ± 0.01 × 10^6^ M^−1^ s^−1^, both at 50%; Figures 2C, S3F and Table S4). Interestingly, while binding was faster at PEC-141, NusA dissociation in the presence of glycine was also significantly faster at PEC-161 than PEC-225 (*k_NusA-off_* = 0.81 ± 0.12 s^−1^ versus 0.47 ± 0.11 s^−1^; Figures 2E, S3E, S3G and Table S4), suggesting that NusA binds earlier but stabilizes more effectively at later transcriptional stages, possibly in coordination with terminator hairpin emergence.

To probe the riboswitch’s intramolecular dynamics, we next used single-molecule kinetic analysis of RNA transient structure (SiM-KARTS^13,20,40^) on co-transcriptionally folded, nascent transcripts. We designed a fluorescent probe targeting P2.2 in the second aptamer, enabling real-time tracking of the β A-minor motif inter-aptamer contact (Figure S1A). Monitoring binding at U225 (Figure 2F-J)—where all aptamer-aptamer interactions can form—revealed two binding rate constants in the absence of glycine: *k_SK-_*_on,fast_ = 2.37± 0.03 × 10^7^ M^−1^ s^−1^ (20%) and *k_SK-_*_on,slow_ = 6.42 ± 0.01 × 10^6^ M^−1^ s^−1^ (80%; Figure 2K and Table S5). Two dissociation rate constants were also identified: *k_SK-_*_off,fast_ = 0.88± 0.02 s^−1^ (91%) and *k_SK-_*_off,slow_ = 0.12 ± 0.02 s^−1^ (9%; Figure 2L and Table S5), suggesting the presence of at least two RNA conformers. In the presence of glycine, the slow association rate decreased significantly (*k_SK-_*_on,slow_ = 2.41 ± 0.01 × 10^6^ M^−1^ s^−1^compared to 6.42 ± 0.01 × 10^6^ M^−1^ s^−1^ in the absence of ligand), indicating that glycine binding promotes formation of the β A-minor motif (Figure 2K-M and Table S5). Probing other elements such as P1.1 and the terminator showed no glycine-dependent changes (Figures S4 and Table S5), underscoring the functional specificity and flexibility of the β A-minor motif for co-transcriptional regulation of the *Bsu-*GTR. In PEC-161, where aptamer 2 is not yet transcribed, glycine had no effect on probe kinetics (Figure S5 and Table S5), confirming that inter-aptamer contacts form later in transcription.

To assess the role of RNAP in these dynamics, we probed released transcripts (RTs) lacking RNAP (Figure S2C). In full-length RT-229, glycine had no effect on SiM-KARTS probe kinetics (Figures 2M, S6F-J and Table S5), and probe binding was significantly slower (4.08 ± 0.01 × 10^6^ M^−1^ s^−1^) than in the corresponding PEC, suggesting a shift to a more thermodynamically stable state. Since RNAP protects ∼14 nt at the 3’ end of the nascent RNA^41^, we hypothesized that release of the RNA allows the terminator hairpin to form fully, locking the structure into a nonresponsive conformation ^20,42,43^. To test this, we examined RT-196, lacking the terminator but retaining both aptamers. While glycine still had no effect on probe kinetics (Figures 2M, S6A-E and Table S5), probe binding improved 1.5-fold compared to RT-229, indicating that the terminator competes with aptamer folding and ligand binding, consistent with prior studies^11,15–18,20^.

Together, these results demonstrate that glycine binding and NusA-stabilized pausing establish a transcriptional checkpoint at U225 that promotes β A-minor motif formation. This checkpoint likely extends the window for productive aptamer interaction, facilitating transcription readthrough upon ligand binding.

### Inter-aptamer contacts pre-organize the glycine binding sites

Previous studies of RNA have shown that co-transcriptional folding pathways often include transient, non-native structures that gradually resolve as the transcript is extended^16,18,44,45^. Our finding that *Bsu*-GTR folding is punctuated by strategically positioned transcriptional pause sites suggests that these pauses help coordinate key structural transitions, including the formation of native folds and glycine binding. To systematically identify intermediate folding states along the *Bsu*-GTR folding pathway, we applied <u>v</u>ariable length <u>T</u>ranscription <u>E</u>longation <u>C</u>omplex RNA structure <u>prob</u>ing (TECprobe-VL^18^) using dimethyl sulfate (DMS) in the absence and presence of 1 mM glycine (Figures 3, S7, and S8). TECprobe-VL, like other co-transcriptional RNA structure probing approaches, leverages high-throughput chemical probing to assess the structure of nascent RNA folding intermediates. Nascent RNA is chemically probed in the context of transcription elongation complexes (TECs) that have been halted at every position of a DNA template, allowing us to map natively folded RNA structures across transcript lengths and detect folding transitions as transcript length-dependent changes in reactivity patterns.

**Figure 3.**
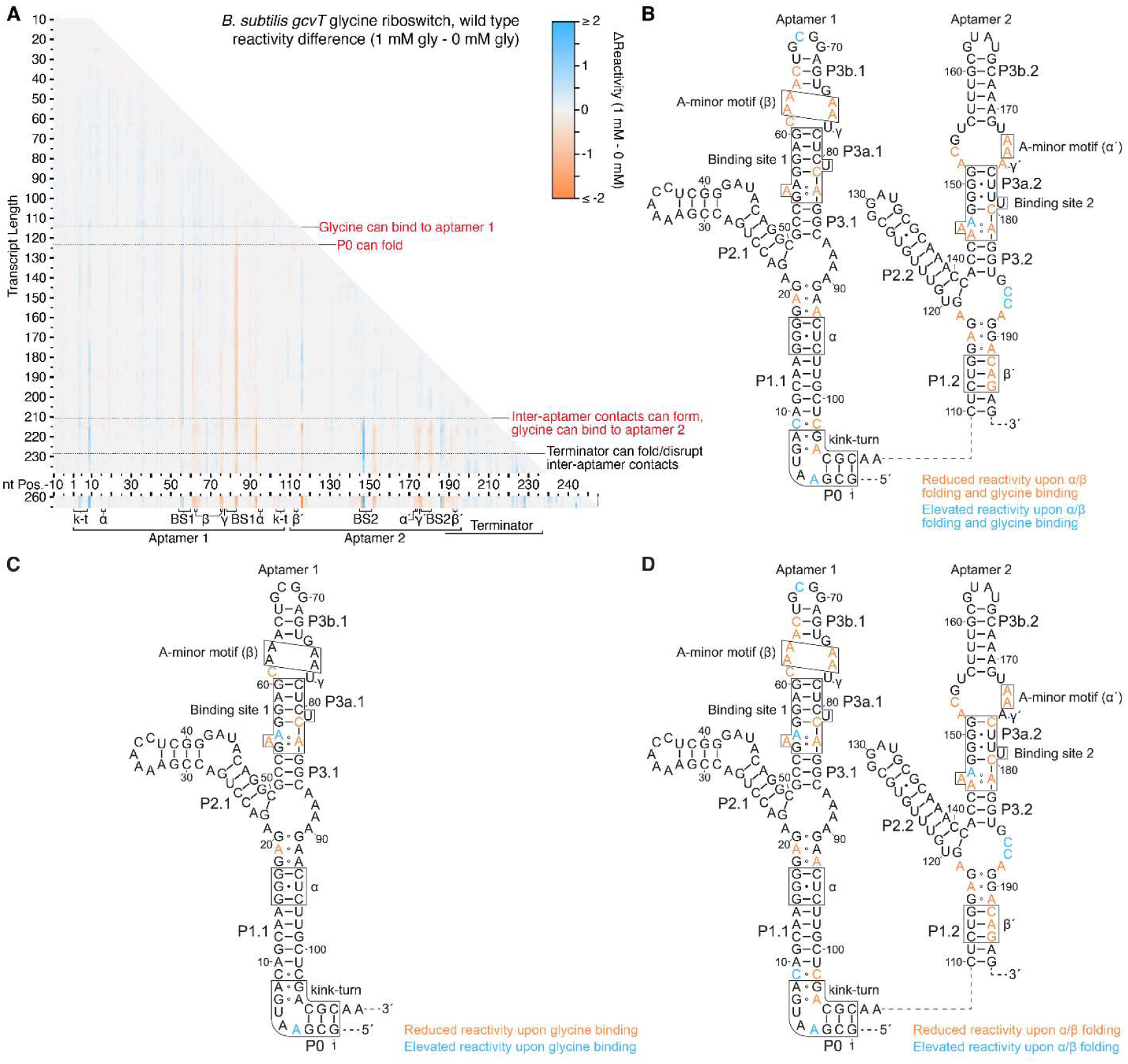
Glycine- and inter-aptamer contact-dependent changes in wild type *Bsu*-GTR aptamer DMS reactivity. (**A**) TECprobe-VL DMS reactivity difference matrix. Transcripts 239-259, which were not enriched, are excluded. (**B-D**) *B. subtilis gcvT* glycine aptamer secondary structures colored to indicate nucleotides that exhibit reduced or elevated DMS reactivity upon α and β A-minor motif folding in the presence of glycine (B), glycine binding to aptamer 1 (C), and upon α and β A-minor motif folding in the absence of glycine (D). TECprobe-VL reactivity data are from the experiment shown in Supplementary Figure 7 in which two independent replicates were processed separately and merged. Gly, glycine; k-t, kink-turn; BS1, binding site 1; BS2, binding site 2.

The DMS reactivity profile of the full-length *Bsu*-GTR aptamers is consistent with the established secondary structure (Figure S7). Comparing reactivity profiles with and without 1 mM glycine revealed both glycine-dependent and glycine-independent reactivity changes, corresponding to direct ligand binding and the formation of inter-aptamer contacts, respectively (Figure 3). Glycine binding to aptamer 1 was first detected around position +115 as decreased reactivity at A55, C82, and A83, accompanied by increased reactivity at A56 (Figures 3C and S8A). Reduced reactivity at A55, C82, and A83 is consistent with the glycine-dependent formation of a network of hydrogen bonds within binding site 1, as observed in the crystal structure of the glycine-bound *Fusobacterium nucleatum* (*Fnu*) GTR (PDB: 3P49), which likely reduces the accessibility of these nucleotides to DMS modification^30^. Conversely, increased reactivity at A56 suggests a glycine-bound conformation in which the Watson-Crick face of A56 remains unpaired and solvent-exposed, again consistent with the *Fnu* GTR structure^30^. In the absence of glycine, we observed reduced reactivity at sites within α′, β, and β′ and adjacent to α near a transcript length of +211, as expected for the formation of the α and β A-minor motifs (Figures 3D and S8D, E). However, formation of the γ interaction was not detected without glycine (Figure S8F). In coordination, the glycine-induced reactivity changes identified in aptamer 1 (at A55, A56, C82, and A83) both occurred in aptamer 1 and were mirrored at the corresponding nucleotides in aptamer 2 (A146, A147, C180, and A181; Figures 3D and S8A, B). This suggests that pre-organization of both glycine binding sites occurs via the α and β A-minor tertiary structure contacts independently of glycine binding. Upon glycine addition, these changes were amplified, and reduced reactivity at A175 indicated that the γ Hoogsteen base pair is glycine-dependent, acting as a linchpin of inter-aptamer docking (Figures 3B and S8). Together, these observations suggest that inter-aptamer contacts can form prior to glycine binding, pre-organizing both aptamer binding sites, and that glycine subsequently stabilizes these interactions, closing the γ linchpin.

To further confirm these glycine-dependent structural changes, we analyzed the folding intermediates of three riboswitch variants in which one or both glycine-binding sites were disrupted (Figures S9-S14). As expected, disruption of binding sites 1 and 2 by the U81G/U179G double substitution variant eliminated all glycine-dependent reactivity changes but did not disrupt inter-aptamer contacts (Figure S10). This variant retained glycine-independent pre-organization of binding site 2, but not binding site 1, likely due to U81G-induced structural disruption, as C82 and A83 were constitutively unreactive after emerging from RNAP (Figure S10A, B). Disruption of binding site 1 alone (U81G) abolished glycine-dependent reactivity changes in aptamer 1 (Figure S12A), weakened the glycine-mediated stabilization of α and β (Figure S12D, E), and reduced―but did not eliminate―ligand-induced reactivity changes in aptamer 2 (Figure S12B). Conversely, disruption of binding site 2 alone (U179G) had no effect on glycine binding by aptamer 1 (Figure S14A), similarly weakened the glycine-mediated stabilization of α and β (Figure S14D, E), and abolished glycine-induced reactivity changes in aptamer 2 (Figure S14B). The glycine-dependent formation of the γ interaction was observed in U81G, but not in U179G (Figures S12F and S14F), indicating that γ folding either requires glycine binding to aptamer 2 or is abolished by the U179G substitution. Glycine-independent pre-organization of binding site 2 was detected in both the U81G and U179G variants (Figures S12B and S14B). Glycine-independent pre-organization of binding site 1 was lost in the U81G variant due to the associated structural changes described above, but was detected in the U179G variant (Figures S12A and S14A). These results further support a model in which the α and β A-minor motif contacts promote pre-organization of both glycine binding sites independently of glycine binding and independently of each other.

To further test whether inter-aptamer contacts are required for binding site pre-organization, we characterized the folding of variants that directly disrupt the β A-minor motif (A76G and A75U/A76G; Figures S15-S18). In the absence of glycine, both variants abolished nearly all reactivity signatures of α and β contact formation, with the exception of decreased reactivity at A116 in P1.2 (Figures S16D, E and S18D, E). Consistent with the double substitution variant A75U/A76G being more disruptive, glycine-induced folding of α and β A-minor contacts was observed in A76G but not in A75U/A76G (Figures S16D, E and S18D, E). Although glycine-independent formation of the α and β contacts was not evident in A76G, glycine addition triggered their formation―suggesting either transient formation stabilized by ligand binding or ligand-induced formation. A76G also suppressed glycine-independent pre-organization at the glycine binding sites except for partially decreased reactivity at A146 and C180 (Figure S16A, B), which may result from the P3.2/P3a.2 structure folding (Supplementary Note 1). However, in the presence of glycine, most signatures of glycine-dependent binding site organization that occur upon α and β folding were recovered (Figure S16A, B). In contrast, A75U/A76G abolished the α and β contact-dependent amplification of glycine binding signatures (Figure S18A, B). These findings demonstrate that the α and β A-minor motifs are central to pre-organizing the glycine binding sites in both aptamers, and that this architecture is stabilized―but not initiated―by glycine binding.

### P0 facilitates inter-aptamer contacts

The formation of P0 establishes a Mg^2+^- and K^+^-dependent kink-turn motif bridging aptamers 1 and 2 of the *Bsu*-GTR (Figure 4A, B)^29^. P0 folding is independent of glycine binding and is first detected near transcript length +126 as decreased DMS reactivity at nucleotides C2, C105, and C107, consistent with the formation of the G1:C107, C2:G106, and G3:C105 base pairs (Figure S19). Once folded, P0 remains generally stable; however, C2 and C107 exhibit transiently increased reactivity upon folding of a downstream non-native hairpin, IH7, within aptamer 2 (Figure S25D). To investigate the role of P0 in co-transcriptional folding, we performed TECprobe-VL on a ΔP0 variant in which nucleotides 1-9 were deleted (Figures 4A, B and S20A, C). The full-length *Bsu*-GTR ΔP0 aptamer reactivity profile aligns with the expected secondary structure, but shows increased reactivity of A and C nucleotides in the inter-aptamer linker relative to wild-type, reflecting the loss of stabilizing P0 base pairs (Figure S20B, D, compare with Figure S7B, D). Upon IH7 formation (Figure S25D), several linker nucleotides show decreased reactivity, suggesting that in the absence of P0 non-native base pairs form between aptamer 2 and the inter-aptamer linker (Figure S19).

**Figure 4.**
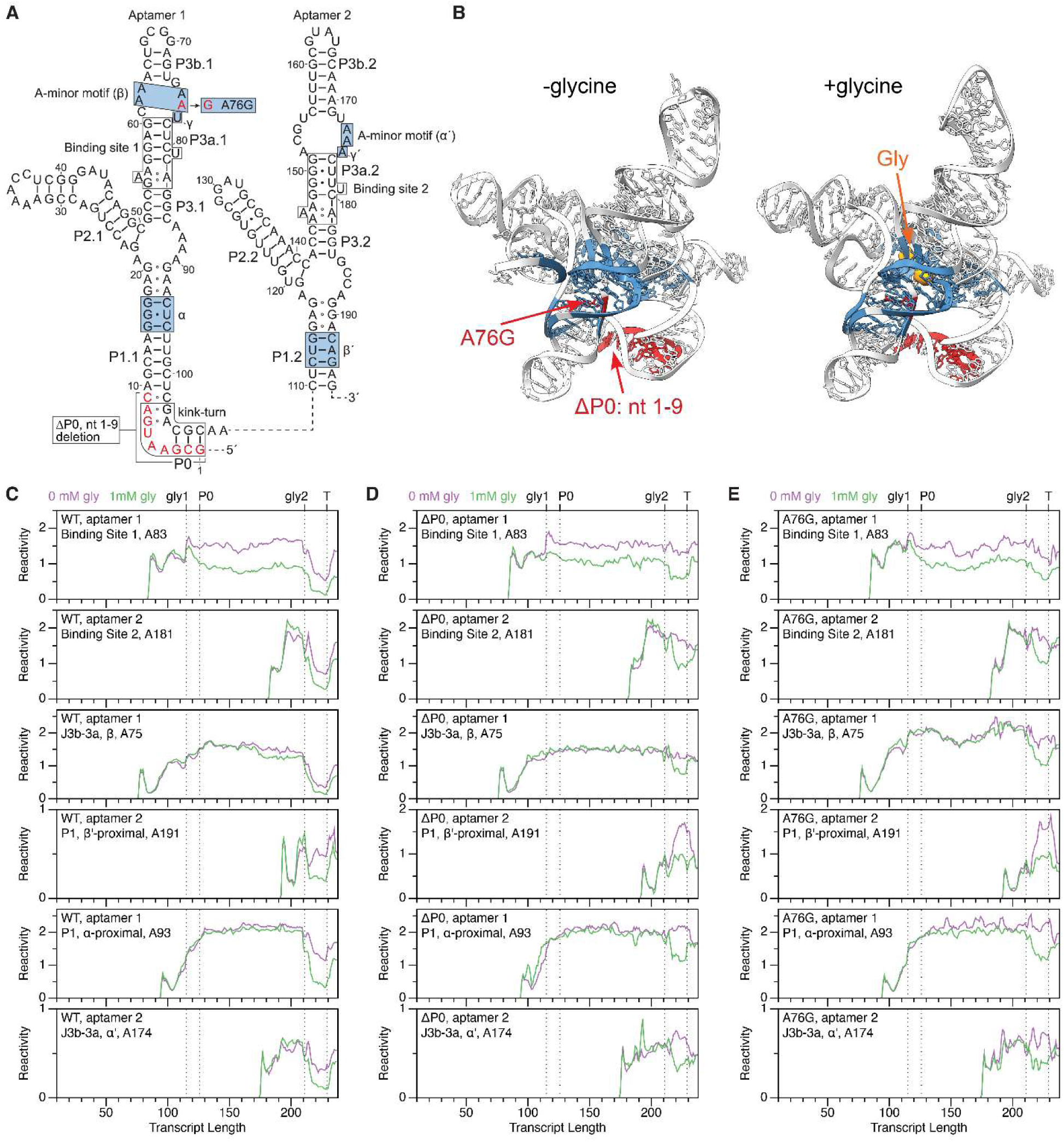
TECprobe-VL analysis of *Bsu-*GTR A76G and ΔP0 variants. (**A**) *Bsu*-GTR secondary structure showing the A76G and ΔP0 perturbations. (**B**) 3D structural models of the antiterminator (right) and terminator (left) annotated to indicate the location of the ΔP0 and A76G perturbations. P0 is located at the site of the kink-turn structural motif, whereas A76 is located in the core region of the GTR, in close proximity to the inter-aptamer tertiary interactions and glycine binding pockets (blue). (**C-E**) TECprobe-VL DMS reactivity trajectories for select nucleotides from the wild type (C), A76G (D), and ΔP0 (E) variants. TECprobe-VL data are from the experiments shown in Supplementary Figures 7, 15, and 20. Gly, glycine.

Strikingly, the ΔP0 variant lacks the glycine-independent reactivity changes associated with inter-aptamer contact formation in the wild-type riboswitch, yet retains the glycine-dependent reactivity changes (Figures 4C, D, S8, and S21). This pattern mirrors that of the A76G variant, which weakens the β A-minor contact, suggesting that in both cases, inter-aptamer contacts are either transiently formed and stabilized by glycine, or form only after ligand binding. Supporting this interpretation, the ΔP0 and A76G variants exhibit similar trends in their folding trajectories and glycine responsiveness (Figures 4D, E, S16, and S21). Collectively, these data indicate that P0 plays a key role in promoting the glycine-independent formation of inter-aptamer contacts during co-transcriptional folding.

### P1 folding drives P3 and P3a folding

The co-transcriptional folding pathway of the *Bsu*-GTR is populated by several non-native intermediate hairpins that are resolved as native structures form (Supplementary Note 1, Figures S22-S26). While the specific intermediates differ between the two aptamers, both exhibit non-native structures that delay the folding of P3/P3a until P1 has formed.

During aptamer 1 folding, decreased reactivity of C78, C80, and C82―nucleotides involved in P3a.1 base pairing—is observed from transcript length +100 to +106 (Figure S23E). However, the reactivity of the P3/P3a.1 nucleotides A59 and C86, as well as A55 and A56 of binding site 1, remains relatively unchanged across these transcripts (Figure S23E, F), suggesting that the reduced reactivity of C78, C80, and C82 does not reflect P3/P3a.1 formation. Instead, a coordinated decrease in the reactivity of A71 suggests the transient formation of a non-native hairpin involving base pairs G70:C80, A71:U79, and G72:C78 (Figures S22E and S23E). Folding of P1.1 is detected as decreased reactivity at A10, C12, A13, and A14 from transcript length +111 to +118, consistent with base pairing in this region of P1.1 (Figure S23F). This transition is coordinated with reduced reactivity at A59, C78, C80, C82, and C86 and increased reactivity at A55, A56, and A83, indicating that P3.1/P3a.1 folding coincides with P1.1 formation (Figures S22F and S23E, F). These structural transitions align with the first glycine-dependent reactivity differences in aptamer 1, which appear at +115 after the aptamer has fully emerged from the RNAP.

The reactivity signatures of P1.2 folding are more subtle than those of P1.1 folding due to the involvement of its upstream nucleotides in transient, non-native structures until P1.2 folds once its downstream nucleotides emerge from the RNAP (Figures S25E, F). Nonetheless, a decrease in reactivity at C110, C112, and C192―nucleotides that pair with G196, G194, and G114 in P1.2―is detected from +211 to +216 (Figure S26A). This coincides with the earliest glycine-dependent reactivity differences in aptamer 2 (Figure S8). Two observations indicate that P3.2/P3a.2 folding is in fact driven by P1.2 formation: (1) the reactivity of A146, A147, and A181 in binding site 2 remains approximately constant until transcript length +211 (Figure S26E), and (2) the reactivity of A142, C143, C144, C176, and C180―participants in P3.2/P3a.2 base pairing―decreases from +214 to +220 (Figure S26C, E). Concurrently, decreased reactivity of A193 and A195—nucleotides pairing with U113 and U111 in P1.2—is also observed, suggesting β A-minor contact formation (Figure S26F). Disruption of this interaction in the A76G, A75U/A76G, and ΔP0 variants results in elevated reactivity in the P1.2 region and at non-canonical base pairs (Figures S15A, B, S17, and S20A, B, compare with Figure S7), further supporting its role in this structural transition. Since P1.2 folding promotes P3.2/P3a.2 formation, these results suggest that inter-aptamer interactions enhance aptamer 2’s affinity for glycine.

### K^+^ and glycine binding pre-organizes inter-aptamer docking during transcription

Our analysis of the ΔP0 variant indicates that P0 facilitates inter-aptamer interactions once both aptamers have emerged from the RNAP. To further investigate how glycine binding influences global docking dynamics at this stage, we examined the behavior of the *Bsu*-GTR when RNAP is paused at U225 (PEC-225), using single-molecule FRET (smFRET) with fluorophores positioned at P0 (Figure 5A). To this end, we employed a stepwise transcription strategy to fluorescently label the 5’ end of the nascent RNA with Cy3 and incorporate an azido-modified NTP at position 10, which was subsequently labeled with DBCO-Cy5 via click-chemistry^13,46^. RNAP was stalled at position 225 by immobilizing the DNA template on streptavidin-coated magnetic beads via a downstream desthiobiotin-TEG modification, allowing for subsequent bead release using a competing biotinylated DNA oligonucleotide (Figure 5A).

**Figure 5.**
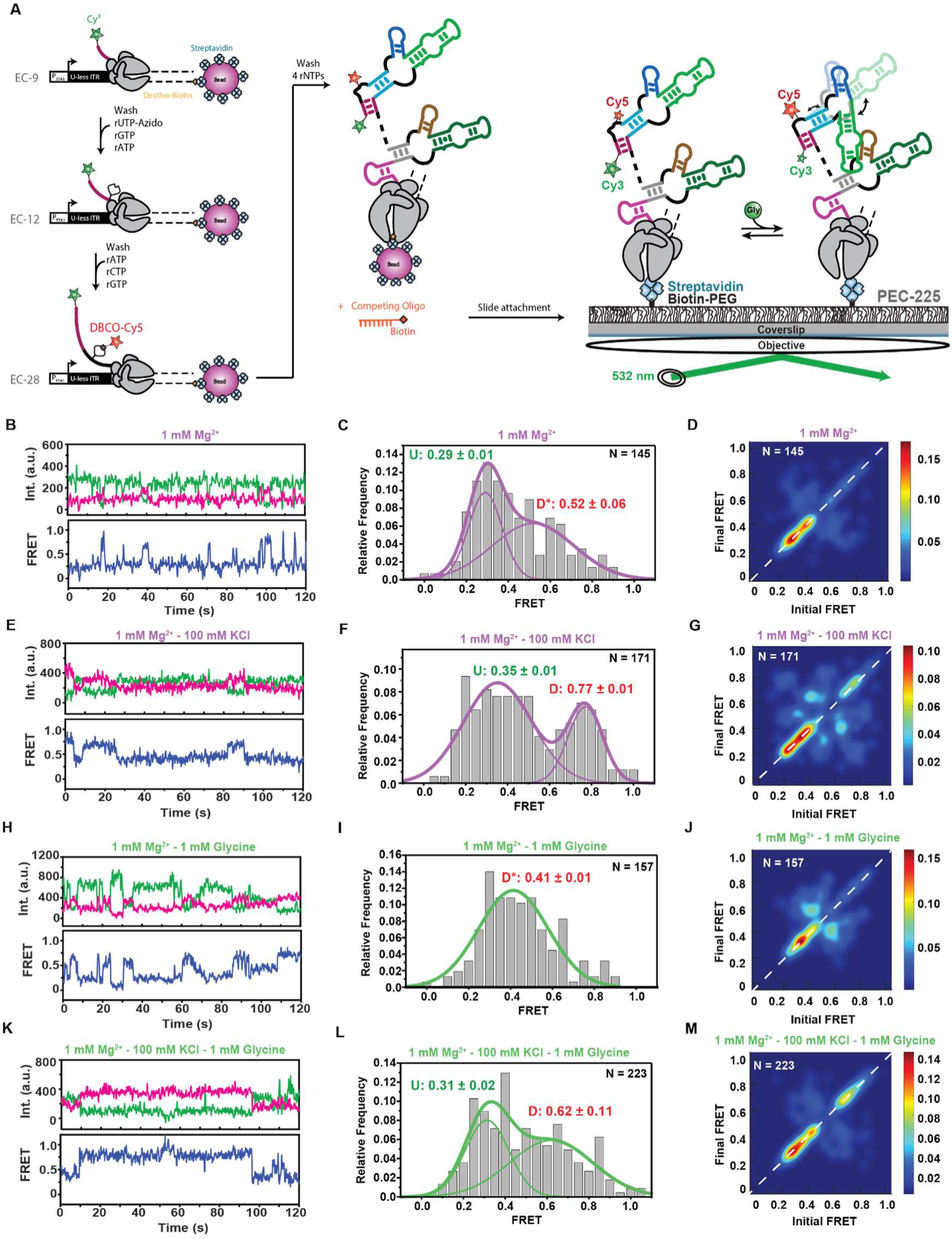
K^+^ and glycine binding pre-organizes inter-aptamer docking during transcription. (A) smFRET experimental setup. The locations of donor (Cy3 – green) and acceptor (Cy5 – red) fluorophores are indicated. (**B-M**) The following plots are shown for PEC-225 in the presence of 1 mM Mg^2+^ (B, C, D), 1 mM Mg^2+^ and 100 mM KCl (E, F, G), 1 mM Mg^2+^ and 1 mM glycine (H, I, J), and 1 mM Mg^2+^, 100 mM KCl and 1 mM glycine (K, L, M): representative dynamic donor-acceptor (top) and smFRET (bottom) traces (B, E, H, K), smFRET histograms (C, F, I, L), and transition occupancy density plots (TODPs) to capture both dynamic and static molecular behaviors (D, G, J, M). The riboswitch conformational state corresponding to each FRET value is indicated as undocked (U) or docked (D). The SD values of each FRET value and total number of traces (N) are shown. TODPs represent dynamic traces as “off-diagonal” and the static traces as “on-diagonal” contour, where the color scale shows the prevalence of each population.

At a near-physiological concentration of Mg^2+^ (1 mM), the riboswitch in PEC-225 sampled two predominant conformations (Figure 5B, C): a low-FRET (∼0.3) state and a mid-FRET (∼0.5) state, which we assigned to undocked and pre-docked architectures, respectively, based on the following observations. The mid-FRET state was dominant (61%) and exhibited a relatively broad distribution, suggesting the presence of multiple, closely related, partially docked conformations (Figure 5C). Transition Occupancy Density Plot (TODP) analysis revealed that most low-FRET molecules were static, consistent with a stably undocked state in the absence of ligand (Figure 5D).

Given that K⁺ ions are known to stabilize kink-turn motifs^47,48^, we next examined docking dynamics upon addition of 100 mM KCl (Figure 5E-G). The predominant low-FRET (∼0.35) population remained, indicating that the undocked state persists under these conditions (Figure 5F). Notably, the mid-FRET population shifted to a high FRET value (∼0.77), suggesting that K^+^ promotes formation of an even more compact, now fully docked state conformation.

We then assessed the effect of glycine binding. In the absence of KCl, addition of 1 mM glycine shifted the riboswitch into a single unified mid-FRET state (Figure 5H, I), whereas in the presence of both glycine and K⁺ ions, low and high-FRET populations could again be discerned―with the high-FRET population increasing from ∼30% to ∼60% relative to the presence of only K^+^ (Figure 5K, L)―indicating that glycine binding shifts the equilibrium toward the fully docked state. In contrast, in the absence of K⁺ TODP analysis showed that docking events remained transient to a significant extent (Figure 5J), supporting a model in which monovalent K⁺ ions are critical for establishing the glycine-dependent, full inter-aptamer contacts under physiological conditions.

Further supporting this model, in the presence of K⁺ most molecules displayed either static low- or high-FRET values, with glycine further promoting the static high-FRET state (Figure 5G, M). These findings suggest that K⁺ promotes a pre-organized riboswitch architecture in which aptamer 1 is dynamically positioned toward aptamer 2, promoting inter-aptamer interactions that are further stabilized upon glycine binding. Kinetic analysis additionally revealed slow transitions between the conformational states in a sub-population of molecules, consistent with glycine binding stabilizing a compact, stably docking state, as observed for other riboswitches in similar co-transcriptional contexts^13,49^.

Together, our findings demonstrate that under physiological ion conditions, the *Bsu*-GTR is pre-organized via the P0 element at the U225 transcriptional checkpoint. This pre-organization enables the riboswitch to rapidly respond to environmental cues, ultimately promoting transcription activation. These results highlight the critical roles of both K⁺ ions and glycine in *Bsu*-GTR structural compaction and reveal an additional regulatory layer in which intermediate RNA conformations at the PEC follow distinct pathways depending on ion availability and ligand presence.

### Computational modeling reveals how local and global conformational changes couple for efficient riboswitching

To visualize the global conformational effects of glycine binding, we employed computational modeling to generate 3D structures of the *Bsu*-GTR structure under two regulatory conditions: the glycine-bound state, in which P1.2 is stabilized and functions as an antiterminator, and the apo state, which favors P1.2 disruption and promotes formation of the terminator. Coarse-grained molecular dynamics (CGMD) simulations were performed using the IsRNA model^50–52^. The resulting structural models (Figure 6A for the antiterminator and Figure 6B for the terminator) are consistent with our experimental observations. Glycine binding stabilizes the inter-aptamer interaction, the kink-turn motif, and the overall compactness of the *Bsu*-GTR. Notably, despite sequence differences, the *Bsu*-GTR structure closely resembles previously solved GTR structures from *Fnu*^53^ and *Vibrio cholerae* (*Vch*)^31^ (PDB IDs: 6WLM and 3OWI, respectively; Figure 6C). These similarities suggest that GTR tertiary structures and regulatory mechanisms may be conserved across diverse bacterial species.

**Figure 6.**
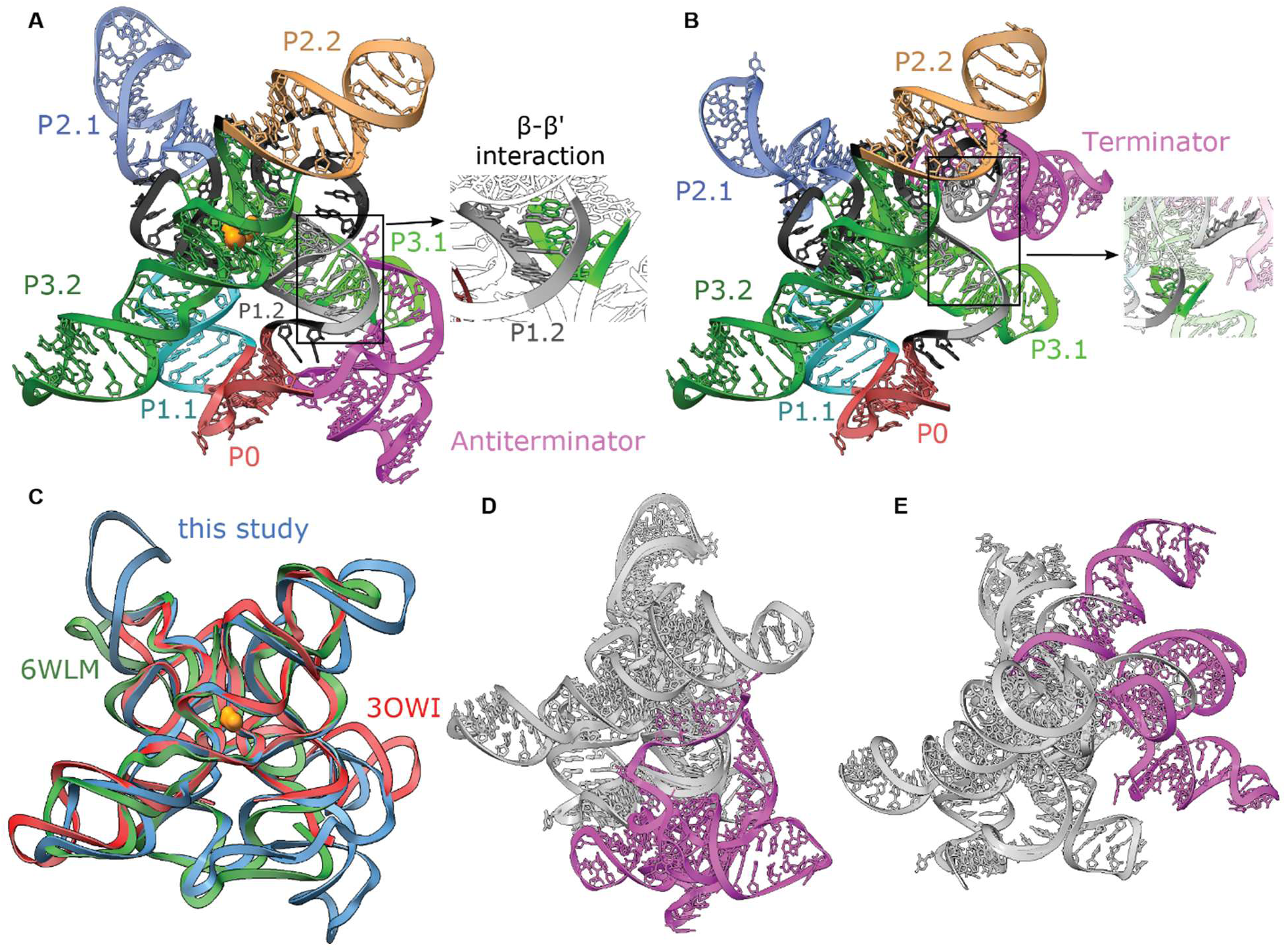
Computational 3D simulation of *Bsu*-GTR terminator and anti-terminator. (**A-B**) The modeled antiterminator and terminator. The structures are colored corresponding to the nucleotide regions shown in Figure 1A and highlight the β-β’ interactions within the antiterminator (A) and the disruption of P1.2 in the terminator (B). (**C**) A comparison of the GTR structure models with *Fnu* and *Vch* tandem glycine aptamer structures (PDB IDs: 6WLM (green) and 3OWI (red)). The two known GTR structures are aligned with the modeled structure in A and show strong structural similarities. The side chains have been hidden for enhanced visual comparison. (**D-E**) Alignment of the three top-ranked models obtained from the simulations for the antiterminator and terminator, respectively. The antiterminator and terminator hairpins are depicted in purple. The alignments without P4 indicate that, in comparison to the antiterminator, the terminator is more dynamic and flexible mainly in the P4 region.

In contrast to our work, however, prior studies mostly have focused on the isolated GTR RNA, excluding the expression platform^24,29,30,33^. Our modeling results indicate that the stability of P1.2 influences not only inter-aptamer compactness but also the conformational dynamics of the antiterminator and terminator elements. Compared to the antiterminator, the terminator structure appears significantly more dynamic and flexible (Figure 6D, E). This increased flexibility may serve as an intrinsic regulatory feature, entropically favoring the terminated state by preventing reformation of P1.2.

This model is further supported by SiM-KARTS data, which show no kinetic differences between the glycine-bound and apo states when probing either P1.2 or the terminator in the context of the full-length riboswitch (Figure S4F–J). These findings suggest that once the full-length GTR has formed, its regulatory architecture is effectively “locked in” and functionally committed to its chosen expression state.

## DISCUSSION

Riboswitches are pervasive across bacterial species and regulate gene expression through sequential structural transitions that occur during transcription. Our study highlights the importance of co-transcriptional checkpoints―specifically, three newly identified RNAP pause sites—as integral nodes of regulation in the *Bsu*-GTR (Figures 1 and S2). These findings contribute to a growing appreciation of strategic transcription pausing as a central feature of both transcriptional and translational riboswitch mechanisms^13,54–58^, where such pauses create kinetic windows that allow structured RNA intermediates to emerge and engage ligands or auxiliary factors. By integrating single-molecule fluorescence microscopy with co-transcriptional RNA structure probing, we resolve transient RNA folding events and characterize intermediate states that are typically inaccessible. This approach supports a model in which the *Bsu*-GTR functions via kinetically driven cooperativity, in contrast to the classic Monod-Wyman-Changeux model, which is frequently applied beyond its original scope of allosteric protein complexes. According to our model, the riboswitch integrates sequential structural and environmental signals from 5′-to-3′ end, fine-tuning gene regulation in response to fluctuating conditions. Specifically, the architecture of the *Bsu*-GTR supports sensing of three effectors sequentially―a glycine molecule as a measure of the intake, utilization, and catabolism of amino acids, followed by K^+^ ions, reflecting the broader physiological state of the bacterial cell^59^, and finally another glycine (Figure 7).

**Figure 7.**
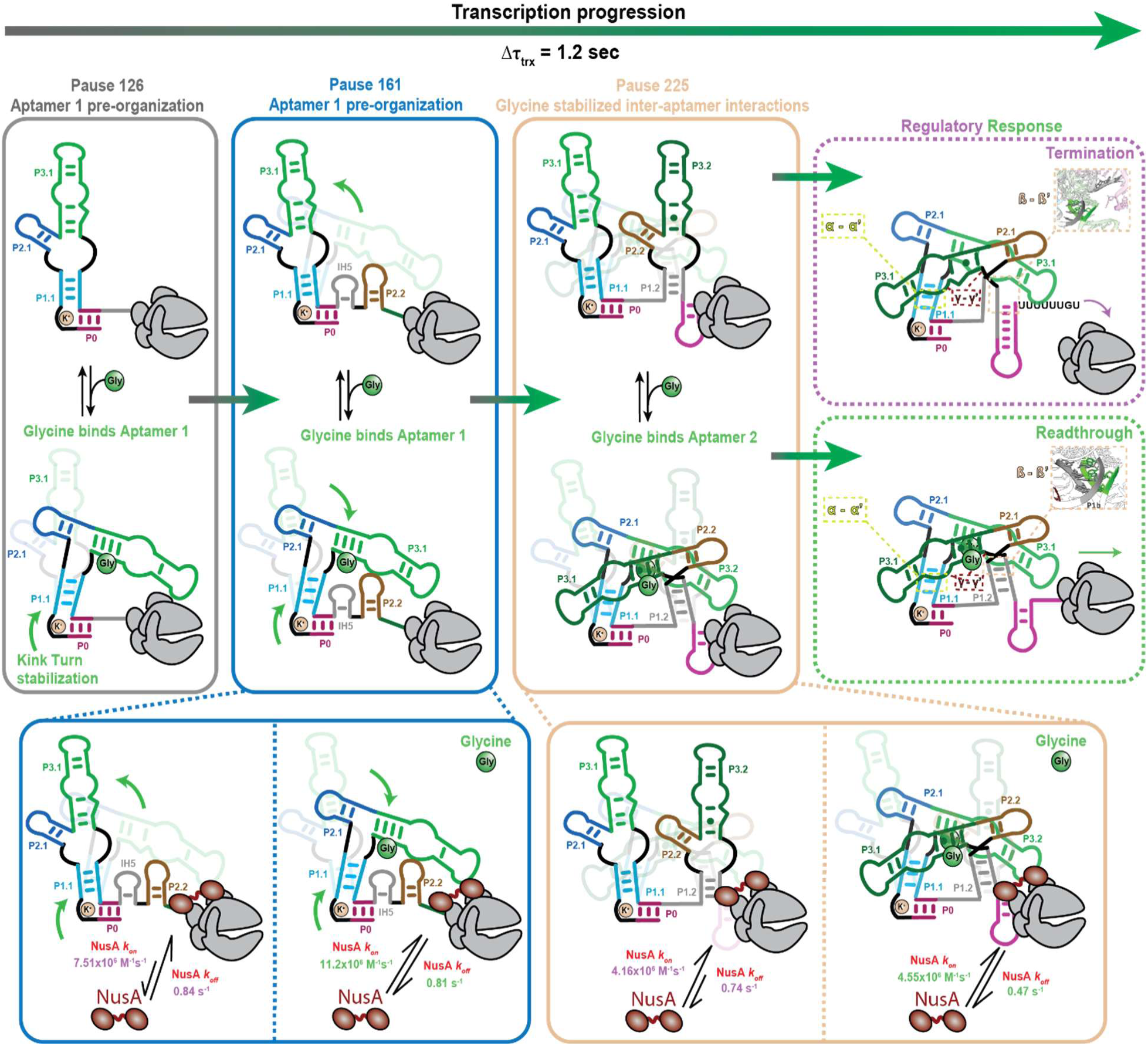
*Bsu-*GTR co-transcriptional folding model. This model depicts the co-transcriptional folding pathway of the *Bsu*-GTR, emphasizing the influence of multiple auxiliary factors at each of three key pause sites that function as transcriptional checkpoints, and illustrates their impact on gene regulation as revealed in the current work.

Central to this model is an RNAP pause at U126, located just downstream of the first aptamer domain (Figure 7). This pause facilitates thermodynamically favorable folding of the initial aptamer structure, promoting formation of aptamer 1 and the potential for K^+^ recognition by the kink-turn motif independently of glycine binding, serving as an master osmotic stress signal (Figures 3, 5, S24, and S28)^59^. Glycine binding then induces local rearrangements in aptamer 1 that stabilize the ligand-binding pocket and adjacent regions (Figures 3 and S28). Previous studies have shown that K^+^ ions stabilize long-range inter-aptamer contacts in both *Bsu* and *Vch* GTR family members^33,48^, a notion supported by our smFRET data and computational modeling (Figures 3, 5 and S27). The sharp kink of the kink-turn motif orients aptamer 1 toward the RNAP exit channel which, together with the 45° angle between P1 and P2^30,31,53^, positions it to interrogate the emerging nascent transcript (Figure 7). Such a juxtaposition will be essential for establishing and stabilizing the three long-range inter-aptamer interactions (Figure 6). This structural logic, together with the functionally positioned U126 pause, reinforces the importance of temporal coordination in riboswitch activation: By enabling the formation of a ligand-competent conformation at the right moment, the system ensures robust gene regulatory outcomes.

As transcription progresses, RNAP encounters a second regulatory checkpoint at position C161 (Figure 7). Our co-transcriptional RNA structure probing uncovered the formation of a previously uncharacterized intermediate hairpin, IH5, located just upstream of P2.2 (Figures S25 and S26). Notably, single-molecule experiments show that IH5 dynamics are unaffected by glycine binding (Figure 2), suggesting that both IH5 and P2.2 represent stable, glycine-independent secondary structures. These may function as further structural locks, maintaining the proximity of the first aptamer to the RNAP exit channel and promoting the establishment of downstream inter-aptamer interactions.

This finding underscores the regulatory potential of non-native, auxiliary secondary structures, which are often overlooked in RNA biology. Such structural intermediates provide temporal separation of folding events, allowing multiple auxiliary factors to act in sequence. Indeed, functional transient secondary structures have been implicated in a variety of riboswitches. For instance, in a translational TPP riboswitch from *E. coli*, a transcription pause promotes Rho-dependent termination by stabilizing a specific conformation^54^. Conversely, in the fluoride riboswitch from *B. cereus*, pausing facilitates NusA recruitment in the ligand-free state, further promoting transcription termination^56^. These examples highlight the critical importance of investigating riboswitch function from a co-transcriptional perspective, where temporal regulation and RNA structure are tightly intertwined.

A third and final regulatory checkpoint occurs at pause site U225, which plays a decisive role in determining whether transcription will proceed or terminate (Figures 1 and 7). This is the earliest transcriptional point at which both ligand-binding sites and inter-aptamer interactions can assemble into their functional configuration. In the absence of glycine, our structural analyses indicate that P1.2 fails to dock stably, leading to an unstable β-β′ interaction, which is essential for proper inter-aptamer compaction (Figures 2 and S24). In contrast, upon glycine binding both P1.2 and the long-range inter-aptamer contacts become stably docked, resulting in a fully compacted GTR conformation (Figures 2, 4 and S8). Importantly, computational folding simulations reveal that this compacted state also requires the proper orientation of the kink-turn; in the absence of P0, inter-aptamer contacts become destabilized, likely due to misalignment of the aptamer domains (Figure S27). Thus, formation of the functional regulatory conformation requires both K^+^ and glycine. Here, K^+^ acts as a dial, tuning the *Bsu*-GTR’s sensitivity to glycine by facilitating the formation of inter-aptamer contacts, which in turn increases aptamer 2’s glycine binding affinity (Figure 4, 5 and S27). The *Bsu*-GTR’s logic effectively functions as a Boolean AND gate when considering how the two glycine binding events stabilize inter-aptamer contacts and promote readthrough. This compacted conformation likely precludes strand displacement by the terminator hairpin, thereby shifting the transcriptional outcome toward readthrough in the presence of ligand (Figure 7).

In addition to this primary mode of glycine-mediated transcription antitermination in which transcription pauses coordinate sequential aptamer folding and ligand binding, the U225 pause also provides a window for late glycine binding by both aptamers 1 and 2. The formation of inter-aptamer contacts pre-organizes both binding sites (Figures 3 and S8). Therefore, if glycine has not bound to aptamer 1 by the time RNAP has reached +225 and inter-aptamer contacts have formed, the increased affinity of the pre-organized binding sites may facilitate rapid glycine binding. In this way, the U225 pause facilitates redundant pathways for glycine-mediated transcription antitermination. Redundant folding mechanisms have been observed in at least two other riboswitch systems: First, the *B. cereus* fluoride riboswitch antiterminates transcription both by delaying terminator hairpin nucleation and, if the terminator nucleates, by blocking terminator base pair propagation^16^. Second, the *C. bacterium oral taxon* 876 str. F0540 ppGpp riboswitch aptamer exhibits a branched folding pathway in which distinct non-native structures converge upon the folding of native aptamer structures^18^. Notably, each of these modes of redundancy is mechanistically distinct and affects a different aspect of riboswitch folding. This suggests that folding pathway redundancy is a versatile mechanism for supporting efficient riboswitch function.

Checkpoint mechanisms such as this are increasingly recognized as a general feature of transcriptional riboswitches, enabling the precise timing of RNA folding events near the RNAP exit channel^13,17,18,60^. In the *Bsu*-GTR, readthrough depends heavily on glycine-induced stabilization of inter-aptamer interactions, particularly the single Hoogsteen base pair γ–γ′ contact, which functions as a final linchpin locking the active conformation (Figures 3, 5 and S8). Disruption of the β A-minor motif compromises these long-range structural interactions and impairs the riboswitch’s regulatory function (Figures 1, 5, S15, and S16).

Together, these findings reinforce the idea that strategically positioned transcription pauses are essential for granting sufficient time for the formation and stabilization of key RNA structures. Comparable strategies are observed in other riboswitch classes, including the lysine- and cobalamin-sensing riboswitches, where long-range kissing-loop interactions are crucial for stabilizing gene-regulatory conformations^57,61^. These parallels support a broader principle: that co-transcriptional folding, modulated by well-timed RNAP pauses, is a conserved and versatile strategy in RNA-based gene regulation.

Previous studies have shown that transcription factors like NusA and NusG promote pauses that affect riboswitch regulation^13,39,39,56,62,63^. NusA significantly prolongs pausing at C161 and U225, especially in the presence of glycine (Figure 1). Single-molecule colocalization experiments further reveal that NusA is preferentially recruited to PEC-161, yet remains bound longer at PEC-225 when glycine is present (Figure 2). These findings suggest that NusA modulates pause duration in a context-dependent manner, shaped by the nascent RNA and RNAP conformations, consistent with observations from a Mn^2+^-sensing riboswitch^13^.

Although NusA is traditionally associated with transcription termination, our data show that its presence—in concert with glycine—enhances readthrough (Figure 1). This suggests that NusA is recruited by nascent RNA structures, such as hairpins, regardless of whether they ultimately promote termination or antitermination. In this way, NusA appears to be differentially guided to distinct pause sites, fine-tuning their duration and ensuring that regulatory inputs are integrated sequentially and with temporal precision.

The initial study describing tandem aptamer glycine riboswitches observed cooperative binding of two glycine molecules by the *Vch* glycine aptamers^24^. Subsequent studies identified the conserved P0 helix, which establishes a kink-turn motif, and determined that the presence of P0 abolishes cooperative glycine binding^28,29,32–34^. Our analysis of *B*. *subtilis gcvT* glycine riboswitch folding suggests a mechanism by which deletion of P0 causes cooperative glycine binding: In the presence of P0, inter-aptamer contacts are more likely to form and pre-organize the glycine binding sites independent of glycine binding by stabilizing P1.1 and P1.2. In the absence of P0, the formation/stabilization of inter-aptamer contacts requires glycine binding to aptamer 1. Consequently, when P0 is absent, pre-organization of binding site 2 can only occur after glycine has bound to aptamer 1, emphasizing the role of P0 in our proposed kinetic cooperativity model, where P0 pre-organizes aptamer 1, thereby priming the inter-aptamer contacts. This in turn stabilizes the second glycine binding site, demonstrating how each pause can contribute sequentially to the overall gene response.

Our investigation of the *Bsu*-GTR, centered on co-transcriptional regulation, uncovered a multi-effector sensing mechanism governed by a series of structured RNA intermediates and coordinated protein factors. By combining multiple complementary methodologies, we demonstrate that nascent RNA folding intermediates serve as scaffolds for the stepwise recruitment and action of ligands and auxiliary factors during transcription elongation. This layered regulatory architecture enables a robust regulatory response to environmental inputs, underscoring the complexity of riboswitch-mediated control. These findings not only expand our understanding of the *Bsu*-GTR system but also provide a broader framework for exploring co-transcriptional regulation in other non-coding RNA systems.

## Resource Availability

## Lead Contact

Requests for further information and resources should be directed to and will be fulfilled by the lead contact, Nils G. Walter.

## Materials Availability

This study did not generate new unique reagents and materials will be available upon request.

## Data and Code Availability

## Data availability

The raw sequencing data generated in this study have been deposited in the Sequence Read Archive (https://www.ncbi.nlm.nih.gov/sra) with the BioProject accession code <u>PRJNA938111</u>. Individual BioSample accession codes are available in Table S9. The processed reactivity data have been deposited in the RNA Mapping Database (https://rmdb.stanford.edu/)^64^. Individual accession codes for each data set are available in Table S10. The ShapeMapper2 output files for these data have been deposited in Zenodo (<u>DOI: 10.5281/zenodo.15264637</u>). The computational 3D simulation data generated in this study is available at GitHub (https://github.com/Vfold-RNA/Computational-3D-simulation-data-for-Bus-GTR). Source data for single molecule and gel-based assays are available through University of Michigan DeepBlue deposit TBD.

## Code availability

TECtools can be accessed at https://github.com/e-strobel-lab/TECtools/releases/tag/v1.2.0. Data visualization scripts can be accessed at https://github.com/e-strobel-lab/TECprobe_visualization/releases/tag/v1.0.0. The Vfold3D-MD simulation code can be accessed at https://rna.physics.missouri.edu/vfold_software_download/vfold3D_download.html. The code used for the single-molecule experiments in this study are available through DeepBlue deposit TBD.

## ACKNOWLEDGMENTS

We thank Chad Torgerson for helpful discussions. This work was supported by the National Institute of General Medical Sciences of the National Institutes of Health under Award Numbers R35GM131922 (to N.G.W), R35GM147137 (to E.J.S), and R35GM134919 (to S-J.C), by the National Institute of Allergy and Infectious Diseases of the National Institutes of Health under Award Number U54AI170660 (to S-J.C.) by the National Science Foundation under Award Numbers MCB 2140320 (to N.G.W.) and CHE 2154924 (to S-J.C.), by the Michigan Economic Development Corporation under Award Number RC112630 (to N.G.W.), and by start-up funding from the University at Buffalo (to E.J.S). RR was supported, in part, by T32GM149391 (Michigan Predoctoral Training in Genetics). The content is solely the responsibility of the authors and does not necessarily represent the official views of the National Institutes of Health.

## AUTHOR CONTRIBUTIONS

Conceptualization, R.A.R., A.C., E.J.S., and N.G.W.; Methodology, R.A.R., A.C., S.Z., and E.J.S; Investigation, R.A.R., A.C., S.S.T., and S.Z.; Writing - Original Draft, R.A.R., A.C., S.Z., and E.J.S; Writing - Review & Editing, R.A.R., A.C., S.Z., S-J.C., E.J.S., N.G.W.; Supervision, S-J.C., E.J.S., and N.G.W; Funding Acquisition, S-J.C., E.J.S., and N.G.W.

## SUPPLEMENTAL INFORMATION

Document S1. Supplementary Notes 1-2, Figures S1-S28 and Table S1-S10

## METHODS

### DNA templates for single-molecule experiments

A 338-nucleotide DNA template including the Glycine-Tandem riboswitch from *B. subtilis* under the control of the T7A1 promoter was cloned into pUC-GW plasmid from Genewiz. Transcription templates for *in vitro* transcription were generated by PCR with Phusion DNA polymerase (ThermoFisher) using the “T7A1-PCR” forward oligonucleotide and the according reverse oligonucleotides. For variant DNA templates, the mutation was inserted in two PCR steps using overlapping oligonucleotides containing the corresponding mutation. Oligonucleotides used for single-molecule experiments are listed in Table S1.

### *In vitro* radioactive transcription assays

Halted complexes (EC-9) were prepared in transcription buffer TB20 (20 mM Tris-HCl, pH 8.0, 20 mM NaCl, 1 mM MgCl_2_, 14 mM 2-mercaptoethanol, 0.1 mM EDTA) containing 25 µM ATP/CTP mix, 50 nM α^32^P-GTP (3000 Ci/mmol), 10 µM ApU dinucleotide primer (Trilink), and 50 nM DNA template. *E. coli* RNAP holoenzyme (New England Biolabs) was added to 100 nM, and the mixture was incubated for 10 min at 37 °C. The sample was passed through G50 column (GE-Healthcare) to remove any free nucleotides according to the manufacturer. To complete the transcription reaction all four rNTPs were added to a variable concentration (described below) concomitantly with heparin (450 µg/mL) to prevent the re-initiation of transcription. Time-resolved single-round *in vitro* transcription experiments were performed using 25 µM rNTPs without or with 1 mM glycine. The mixture was incubated at 37 °C, and reaction aliquots were quenched at the specified times by mixing with an equal volume of loading buffer (95% formamide, 1 mM EDTA, 0.1% SDS, 0.2% bromophenol blue, 0.2% xylene cyanol). Sequencing ladders were prepared by combining the halted complex with a chase solution containing 250 µM of each rNTP, in addition to one 3’-OMe rNTP (at 25 µM for 3’-OMe GTP and 15 µM for 3’-OMe ATP, UTP and CTP). Samples were denatured by heating at 90°C in 90% formamide, 0.1% SDS, 0.1 mM EDTA before loading 5 µL of each onto a denaturing 8 M urea, 6% polyacrylamide sequencing gel at 65mW. The gel was dried and exposed to a phosphor screen (typically overnight), which was then scanned on a Typhoon Phosphor Imager (GE Healthcare).

### Transcription data analysis

To determine the R_50_ for regulation of antitermination by Glycine, the relative intensity of the full-length product was divided by the total amount of RNA transcript (full-length + terminated product) for each Glycine concentration. Percent readthrough was plotted against the Glycine concentration using equation: Y=Y_0_ + ((M1×X)/(T_50_+X)), where Y_0_ is the value of Y at X_min_ (no ligand condition) and M1 = Y_max_ - Y_min_.

Transcription pause half-life was determined by calculating the fraction of each pause species out of total RNA for each time point, which was analyzed in GraphPad Prism with pseudo-first-order kinetics to extract the half-life^65^. For each determination, we have subtracted the background signal. Error bars in transcription quantification represent the standard deviation of the mean from independent replicates.

### Preparation of fluorescently labeled nascent transcripts

*In vitro* transcription reactions were performed in two steps to allow the specific incorporation of Cy3 at the 5-end of the RNA sequence. Transcription reactions were performed in the same transcription buffer described above (TB20). Transcription was initiated by adding 100 μM 5’-Cy3-ApU dinucleotide and 25 μM ATP/CTP/GTP nucleotides and incubating the reaction at 37 °C for 10 min, which generates fluorescently-labeled synchronized TECs. The sample was passed through a G50 column (GE-Healthcare) to remove any free nucleotides, and the transcription was resumed upon addition of all four rNTPs at the indicated concentration and heparin (450 μg/mL) to prevent transcription re-initiation. The resulting nascent transcript was hybridized to the 5’ biotinylated capture probe (Anchor Bio oligonucleotide -Table S1) complementary to the 3’ end capture sequence, allowing immobilization of the complex to the microscope slide. The capture probe was mixed to a ratio of 10:1 with the RNA transcript and added for 5 min before flowing the whole complex to the microscope slide. When generating PECs, the DNA templates contained a biotin at the 5’ end of the template DNA strand. Streptavidin was mixed to a ratio of 5:1 with the DNA template for 5 min prior to start the transcription reaction. Glycine (when present) was always added co-transcriptionally at a final concentration of 1 mM, and the same concentration of ligand was added into the corresponding buffer during subsequent dilutions.

### Stepwise transcription followed by Click labeling for smFRET experiments

DNA templates for transcription were produced by PCR using oligonucleotides containing the T7A1 promoter sequence (Supplementary Table 1). The formation of stalled ECs was ensured by using DNA templates containing a Desthio-biotin at the 5’ end of the antisense strand. DNA templates (500 nM) were incubated with 20 µL Streptavidin-coated magnetic beads (Thermofisher Dynabeads M-270) overnight at room temperature in transcription buffer (20 mM Tris HCl pH 8.0, 20 mM MgCl2, 20 mM NaCl, 14 mM 2-mercaptoethanol, and 0.1 mM EDTA). In step 1, transcription reactions were initiated by adding 2 µM RNAP holoenzyme, 100 µM Cy3-ApU dinucleotide, 50 µM rATP/rGTP/rCTP nucleotides at 37°C for 20 min. The sample was washed three times with five volumes of transcription buffer to remove any free nucleotides. In Step 2, RNAP was walked to the next position (EC-12) by adding 50 µM rATP/rGTP/ rUTP-azido at Room temperature for 5 min. The sample was washed three times with five volumes of transcription buffer to remove any free nucleotides. In Step 3, RNAP was walked to the next position (EC-28) by adding 50 µM rATP/rGTP/rCTP at Room temperature for 5 min. The sample was washed three times with five volumes of transcription buffer to remove any free nucleotides. The azide-carrying ECs were incubated with 500 µM DBCO-Cy5 (Jena Bioscience) in the transcription buffer for 1 h at 37°C. ECs were then washed five times to remove unreacted dyes. In step 5, elongation to the roadblock position was achieved by adding 1 mM rNTPs at 37°C for 5 min. To elute the PECs from the beads 10 µM competing biotinylated oligonucleotide was added at 37°C for 20 min. Recovered complexes from the supernatant were directly flowed on a PEG/PEG-biotin treated microscope slide preincubated with 0.2 mg/mL streptavidin.

### Single-molecule experiments

All SiM-KARTS experiments were performed using a prism-based TIRF microscope based on an Olympus IX-71 frame equipped with a 60X∼ water-immersion objective (Olympus UPlanApo, 1.2 NA). SmFRET experiments were performed using the Oxford Nanoimager (ONI) microscope in TIRF (total internal reflection fluorescence) mode. All movies were collected at 100 ms time resolution using an intensified charge-coupled device camera (Hamamatsu C13440-20CU scientific complementary metal-oxide semiconductor camera). PEG-passivated quartz slides with a microfluidic channel containing inlet and outlet ports for buffer exchange were assembled as described in previous works^40,66^. The surface of the microfluidic channel was coated with streptavidin (0.2 mg/mL) for 10 to 15 min prior to flowing the nascent transcripts hybridized to the CP or PEC for smFRET. In the case of PEC analysis, the fluorescent PECs were directly injected into the channel surface using the biotin-streptavidin roadblock for immobilization.

For SiM-KARTS experiments, the nascent RNA transcripts were diluted in SiM-KARTS buffer (16 mM Tris-HCl, pH 8.0, 330 mM KCl, 0.1 mM EDTA, 0.1 mM dithiothreitol, 5 mM MgCl_2_). NusA-Cy5 (2 nM or 5 nM) or the SiM-KARTS probe (2 nM or 25 nM) was injected into the channel in the corresponding imaging buffer along with an enzymatic oxygen scavenging system consisting of 44 mM glucose, 165 U/mL glucose oxidase from *Aspergillus niger*, 2,170 U/mL catalase from *Corynebacterium glutamicum*, 1 mg/mL bovine serum albumin and 5 mM Trolox to extend the lifetime of the fluorophores and to prevent photo-blinking of the dyes. The raw movies were collected for 15 min with direct green (532 nm) and red (638 nm) laser excitation for SiM-KARTS and NusA colocalization assays and 3 min with direct green (532 nm) for smFRET.

### Data acquisition and analysis for SiM-KARTS and colocalization assays

Locations of molecules and fluorophore intensity traces for each molecule were extracted from raw movie files using custom-built MATLAB codes. Traces were manually selected for further analysis using the following criteria: single-step photobleaching of Cy3, ≥ 2 spikes of Cy5 fluorescence of more than 2-fold the background intensity. Traces showing binding events were idealized using a two-states (bound and unbound) model using a segmental *k*-means algorithm in QuB^67^. From the idealized traces, dwell times of NusA or the SiM-KARTS probe in the bound (*τ*_bound_) and the unbound (*τ*_unbound_) states were obtained. Cumulative plots of bound and unbound dwell-time distributions were plotted and fitted in Origin lab with single-exponential or double-exponential functions to obtain the lifetimes in the bound and unbound states. The dissociation rate constants (*k*_off_) were calculated as the inverse of the *τ_bound_*, whereas the association rate constants (*k*_on_) were calculated by dividing the inverse of the *τ*_unbound_ by the concentration of SiM-KARTS probe or NusA used during the data collection.

### Data acquisition and analysis for smFRET assay

Locations of molecules and fluorophore intensity traces for each molecule were extracted from raw movie files using custom-built MATLAB codes. Single-molecule traces were then visualized using custom Matlab code and only those with a minimum combined intensity (Cy3 + Cy5 intensity) of 300 A.U., showing single-step photobleaching of the dyes, a signal-to-noise ratio of >3, and longer than 5 s were selected for further analysis. Selected traces were then background-subtracted to correct for crosstalk and (minimal) bleed-through. We calculated the FRET ratio as *I_A_*/(*I_A_*+*I_D_*), where *I_A_* and *I_D_* are the background-corrected intensities of the acceptor (Cy5) and donor (Cy3), respectively. FRET histograms were made using the first 100 frames of all traces in each condition and fit with a sum of Gaussians using OriginPro 8.5. For kinetic analysis, traces were idealized with a three-state model corresponding to Undocked (low-FRET), and Docked (high-FRET) states using the segmental k-means algorithm in QuB software as previously described^67^. Cumulative dwell-time histograms were plotted from all extracted dwell times and fit with single- or double-exponential functions using OriginPro 8.5 to obtain the lifetimes in the undocked (*τ*_undock_) and docked (*τ*_dock_) states. Rate constants of docking and undocking were then calculated as *k*_dock_ = 1/*τ*_undoc*k*_ and *k*_undock_ = 1/*τ*_dock_. For the double-exponential fits, kinetics were calculated similarly using both the short and long dwell lifetimes to obtain the fast and slow rate constants, respectively. The idealized smFRET traces were used for creating transition occupancy density plots (TODPs), which show the fraction of traces/molecules that exhibit a given type of transition at least once. In TODPs, dynamic traces showing a FRET transition (regardless of the number of transitions in that trace) and static traces (with no transitions over the entire trace) are weighted equally, avoiding over-representation of the traces with fast transitions.

### Oligonucleotides for TECprobe-VL DNA template preparation and experiments

All oligonucleotides used in TECprobe-VL experiments were purchased from Integrated DNA Technologies. A detailed description of all oligonucleotides including sequence, modifications, and purifications is presented in Table S6.

### Proteins for TECprobe-VL DNA template preparation and experiments

Q5 High-Fidelity DNA Polymerase, Vent (exo-) DNA polymerase, *E*. *coli* RNA Polymerase holoenzyme, Mth RNA Ligase (as part of the 5’ DNA Adenylation kit), T4 RNA Ligase 2 truncated KQ, ET SSB, RNase H, and RNase I_f_ were purchased from New England Biolabs. TURBO DNase, SuperaseIN, SuperScript II, and BSA were purchased from ThermoFisher. Streptavidin was purchased from Promega.

### Randomly biotinylated DNA template preparation

Randomly biotinylated DNA templates were prepared as described previously^68^. The original procedure is provided below. Table S7 provides details for the oligonucleotides and processing steps used for every DNA template preparation that was used in a TECprobe-VL experiment in this work. Table S8 provides DNA template sequences.

5’ biotinylated DNA that was used as a PCR template when preparing randomly biotinylated DNA templates below was PCR amplified from plasmid DNA using Q5 High-Fidelity DNA Polymerase (New England Biolabs) and primers PRA1_NoMod.F and HP4_5bio.R (Table S6). PCR was performed as three 100 μl reactions containing 1X Q5 Buffer (New England Biolabs), 1X Q5 High GC Enhancer (New England Biolabs), 200 μM dNTP Solution Mix (New England Biolabs), 250 nM PRA1_NoMod.F, 250 nM HP4_5bio.R, 20 pM template DNA, and 0.02 U/μl Q5 DNA polymerase using the following thermal cycler protocol with a heated lid set to 105 °C: 98 °C for 30 s, [98 °C for 10 s, 65 °C for 20 s, 72 °C for 20 s] x 30 cycles, 72 °C for 5 minutes, hold at 10 °C. The resulting PCR product was purified by UV-free agarose gel extraction using a QIAquick gel extraction kit^69^. A step-by-step protocol for this procedure is available^68^.

Randomly biotinylated DNA templates for TECprobe-VL experiments were PCR amplified from a 5’-biotinylated linear DNA template using Vent (exo-) DNA polymerase (New England Biolabs) and primers PRA1_NoMod.F and HP4_5bio.R (Table S6). 200 μl PCRs contained 1X ThermoPol Buffer (New England Biolabs), 250 nM PRA1_NoMod.F, 250 nM HP4_5bio.R, 20 pM template DNA, 0.02 U/μl Vent (exo-) DNA polymerase, 200 μM dNTP Solution Mix, and a concentration of biotin-11-dNTPs (PerkinElmer, Biotium) that favored the incorporation of ∼2 biotin modifications in the transcribed region of each DNA template^70^. PCR was performed as two 100 μl reactions in thin-walled tubes using the following thermal cycler protocol with a heated lid set to 105 °C: 95 °C for 3 min, [95 °C for 20 s, 58 °C for 30 s, 72 °C for 30 s] x 30 cycles, 72 °C for 5 min, hold at 12 °C. PCR products were purified as described below in the section *SPRI bead purification of DNA*, eluted into 50 μl of 10 mM Tris-HCl (pH 8.0), and quantified using the Qubit dsDNA Broad Range Assay Kit (Invitrogen) with a Qubit 4 Fluorometer (Invitrogen). A step-by-step protocol for this procedure is available^68^.

### SPRI bead purification of DNA

SPRI beads were prepared in-house using the ‘DNA Buffer’ variation of the procedure by Jolivet and Foley^71^. Samples were mixed with an equal volume of SPRI beads, incubated at room temperature for 5 min, and placed on a magnetic stand for 3 min so that the beads collected on the tube wall. The supernatant was aspirated and discarded, and the beads were washed twice by adding a volume of 70% ethanol at least 200 μl greater than the combined volume of the sample and SPRI beads to the tube without disturbing the bead pellet while it remained on the magnetic stand. The samples were incubated at room temperature for 1 min before aspirating and discarding the supernatant. Residual ethanol was evaporated by placing the open microcentrifuge tube in a 37 °C dry bath for ∼15 s with care taken to ensure that the beads did not dry out. Purified DNA templates were eluted by resuspending the beads in a variable amount of 10 mM Tris-HCl (pH 8.0) (depending on the procedure, details are in each relevant section), allowing the samples to sit undisturbed for 3 min, placing the sample on a magnetic stand for 1 min so that the beads collected on the tube wall, and transferring the supernatant, which contained purified DNA, into a screw-cap tube with an O-ring.

### TECprobe-VL

TECprobe-VL experiments were performed as described previously by Szyjka and Strobel with target RNA-specific modifications to the conditions used for single-round *in vitro* transcription^18^. These modifications are presented in the context of the original methods below.

#### Single-round in vitro transcription for TECprobe-VL

All single-round *in vitro* transcription reactions for TECprobe-VL experiments were performed as 60 μl reactions containing 1X Transcription Buffer [20 mM Tris-HCl (pH 8.0), 50 mM KCl, 1 mM dithiothreitol (DTT), and 0.1 mM EDTA], additional Tris-HCl (pH 8.0) to a total concentration of 100 mM, 0.1 mg/ml Molecular Biology-Grade BSA (Invitrogen), 50 μM high-purity NTPs (Cytiva), 10 nM randomly biotinylated template DNA, and 0.024 U/μl *E. coli* RNA polymerase holoenzyme (New England Biolabs). Note that the concentration of Tris (pH 8.0) used for dimethyl sulfate (DMS) probing experiments is higher than our standard concentration (20 mM) to minimize pH changes during the DMS probing reaction. When present, 1 mM glycine was included in the transcription master mix. At the time of preparation, each TECprobe-VL reaction was 48 μl due to the omission of 10X (1 μM) streptavidin (Promega) and 10X Start Solution [100 mM MgCl_2_, 100 μg/ml rifampicin (Gold Biotechnology)] from the reaction.

Single-round *in vitro* transcription reactions were incubated at 37 °C for 10 min to form open promoter complexes. 6 μl of 1 μM streptavidin was then added for a final concentration of 100 nM streptavidin, and reactions were incubated for an additional 10 min at 37 °C. Transcription was initiated by adding 6 μl of 10X Start Solution to the reaction for a final concentration of 10 mM MgCl_2_ and 10 μg/ml rifampicin. The transcription reaction was incubated at 37 °C for 2 min before chemical probing was performed as described below in the section *RNA chemical probing*.

#### RNA chemical probing

Dimethyl sulfate (DMS) probing was performed by splitting the sample into 25 μl aliquots and mixing with 2.8 μl of 6.5% (v/v) DMS (Sigma-Aldrich) in anhydrous ethanol (Sigma-Aldrich) [(+) sample)] or with anhydrous ethanol [(-) sample] and incubating the samples at 37 °C for 5 min. The DMS probing reaction was quenched by adding beta-mercaptoethanol to 2.8 M and incubating the sample at 37 °C for 1 min. 75 μl of TRIzol LS reagent was added to each sample to stop the *in vitro* transcription reaction and the samples were vortexed.

#### RNA purification

Samples, which contained 34.5μl of the cotranscriptional RNA chemical probing reaction in 75 μl of TRIzol LS, were extracted as follows: 20 μl of chloroform was added to each sample, and the samples were mixed by vortexing and inverting the tube and centrifuged at 18,500 x g and 4 °C for 5 min. The aqueous phase was transferred to a new tube and precipitated by adding 1.5 μl of GlycoBlue Coprecipitant (Invitrogen) and 50 μl of ice-cold isopropanol and incubating at room temperature for 15 min. The samples were centrifuged at 18,500 x g and 4 °C for 15 min, the supernatant was aspirated and discarded, 500 μl of ice cold 70% ethanol was added to each sample, and the tubes were gently inverted to wash the samples. The samples were centrifuged at 18,500 x g and 4 °C for 2 min and the supernatant was aspirated and discarded. The samples were centrifuged again briefly to pull down residual liquid, which was aspirated and discarded. The pellet was then resuspended in 25 μl of 1X TURBO DNase buffer (Invitrogen), mixed with 0.75 μl of TURBO DNase (Invitrogen), and incubated at 37 °C for 15 min. 75 μl of TRIzol LS reagent was added to stop the reactions and a second TRIzol extraction was performed as described above, except that the pellet was resuspended in 5 μl of 10% (v/v) DMSO.

#### RNA 3’ adapter ligation

9N_VRA3 adapter oligonucleotide (Table S6) was pre-adenylated with the 5’ DNA Adenylation Kit (New England Biolabs) according to the manufacturer’s protocol at a 5X scale. Briefly, 100 μl of a master mix that contained 1X DNA Adenylation Buffer (New England Biolabs), 100 μM ATP, 5 μM 9N_VRA3 oligo, and 5 μM Mth RNA Ligase (New England Biolabs) was split into two 50 μl aliquots in thin-walled PCR tubes and incubated at 65 °C in a thermal cycler with a heated lid set to 105 °C for 1 hour. Following the reaction, 150 μl of TRIzol LS reagent was added to each 50 μl reaction and the samples were extracted as described above in the section *RNA purification*, except that reaction volumes were scaled to account for the 50 μl reaction volume (40 μl of chloroform was added to the sample-TRIzol mixture and 100 μl of isopropanol was added during the precipitation step). Samples were pooled by resuspending the pellets from each TRIzol extraction in a single 25 μl volume of TE Buffer (10 mM Tris-HCl (pH 7.5), 0.1 mM EDTA). The concentration of the adenylated oligonucleotide was determined using the Qubit ssDNA Assay Kit (Invitrogen) with a Qubit 4 Fluorometer. The molarity of the linker was calculated using 11,142 g/mol as the molecular weight. The adenylation reaction was assumed to be 100% efficient. The linker was diluted to 0.9 μM and aliquoted for future use; aliquots were used within 3 freeze-thaw cycles.

20 μl RNA 3’ adapter ligation reactions were performed by combining purified RNA in 5 μl of 10% DMSO (v/v) from the *RNA purification* section with 15 μl of an RNA ligation master mix such that the final 20 μl reaction contained purified RNA, 2.5% (v/v) DMSO, 1X T4 RNA Ligase Buffer (New England Biolabs), 0.5 U/μl SuperaseIN (Invitrogen), 15% (w/v) PEG 8000, 45 nM 5’-adenylated 9N_VRA3 adapter, and 5 U/μl T4 RNA Ligase 2, truncated, KQ (New England Biolabs). The samples were mixed by pipetting and incubated at 25 °C for 2 hours.

#### SPRI bead purification of RNA

Excess 9N_VRA3 3’ adapter oligonucleotide was depleted using a modified SPRI bead purification that contains isopropanol^72^. 17.5 μl of nuclease-free water and 40 μl of freshly aliquoted anhydrous isopropanol (Sigma-Aldrich) were added to each 20 μl RNA ligation reaction, and the samples were mixed by vortexing. Each sample was then mixed with 22.5 μl of SPRI beads so that the concentration of PEG 8000 was 7.5% (w/v) and the concentration of isopropanol was 40% (v/v) in a sample volume of 100 μl. The samples were incubated at room temperature for 5 min and placed on a magnetic stand for at least 3 min so that the beads collected on the tube wall. The supernatant was aspirated and discarded and the beads were washed twice by adding 200 μl of 80% (v/v) ethanol to the tubes without disturbing the bead pellet while the samples remained on the magnetic stand, incubating the samples at room temperature for 1 min, and aspirating and discarding the supernatant. After discarding the second 80% (v/v) ethanol wash, the sample was briefly spun in a mini centrifuge and placed back onto a magnetic stand for 1 min to collect the beads on the tube wall. The supernatant was aspirated and discarded and the beads were briefly (<15 s) dried in a 37 °C dry bath with the cap of the microcentrifuge tube left open. Purified RNA was eluted by resuspending the beads in 20 μl of 10 mM Tris-HCl (pH 8.0), incubating the sample for 3 min at room temperature, placing the sample on a magnetic stand for 1 min so that the beads collected on the tube wall, and transferring the supernatant into a clean microcentrifuge tube. The eluted RNA was mixed with 11.5 μl of RNase-free water and 40 μl of anhydrous isopropanol. 6 μl of 50% (w/v) PEG 8000 and 22.5 μl of SPRI beads were added to each sample so that the concentration of PEG 8000 was 7.5% (w/v) and the concentration of isopropanol was 40% (v/v) in a sample volume of 100 μl. The RNA was purified as described above a second time, except that the RNA was eluted into 25 μl of 1X Buffer TM (1X Transcription Buffer, 10 mM MgCl_2_).

### cDNA synthesis and cleanup

5 μl of 10 mg/ml Dynabeads MyOne Streptavidin C1 beads (Invitrogen) per sample volume were equilibrated in Buffer TX (1X Transcription Buffer, 0.1% (v/v) Triton X-100)^73^. Briefly, after placing the beads on a magnetic stand and removing the storage buffer, the beads were resuspended in 500 μl of Hydrolysis Buffer (100 mM NaOH, 50 mM NaCl) and incubated at room temperature for 10 minutes with rotation. Hydrolysis Buffer was removed, and the beads were resuspended in 1 ml of High Salt Wash Buffer (50 mM Tris-HCl (pH 7.5), 2 M NaCl, 0.5% (v/v) Triton X-100), transferred to a new tube, and washed by rotating for 5 minutes at room temperature. High Salt Wash Buffer was removed, and the beads were resuspended in 1 ml of Binding Buffer (10 mM Tris-HCl (pH 7.5), 300 mM NaCl, 0.1% (v/v) Triton X-100), transferred to a new tube, and washed by rotating for 5 minutes at room temperature. After removing Binding Buffer, the beads were washed twice with 500 μl of Buffer TX (1X Transcription Buffer supplemented with 0.1% (v/v) Triton X-100) by resuspending the beads, transferring them to a new tube, washing with rotation for 5 minutes at room temperature, and removing the supernatant. After washing the second time with Buffer TX, the beads were resuspended to a concentration of ∼2 μg/μl in Buffer TX (25 μl per sample volume), split into 25 μl aliquots, and stored on ice until use.

The reverse transcription primer, dRP1_5Bio.R (Table S6), contains a 5’ biotin modification and anneals to the 9N_VRA3 adapter. 1 μl of 500 nM dRP1_5Bio.R was added to the purified RNA, and the samples were incubated on a thermal cycler with a heated lid set to 105 °C using the ‘RT anneal’ protocol: pre-heat to 70 °C, 70 °C for 5 min, ramp to 50 °C at 0.1 °C/s, 50 °C for 5 min, ramp to 40 °C at 0.1 °C/s, 40 °C for 5 min, cool to 25 °C. Equilibrated MyOne Streptavidin C1 beads were placed on a magnetic stand to collect the beads on the tube wall, and the supernatant was removed. The bead pellets were resuspended using the samples (which contained dRP1_5Bio.R oligo annealed to RNA), and incubated at room temperature for 15 min on an end-over-end rotator set to ∼15 rpm. The samples were placed on a magnetic stand to collect the beads on the tube wall, the supernatant was aspirated and discarded, and the beads were resuspended in 19.5 μl of reverse transcription master mix, which omits SuperScript II reverse transcriptase at this time. After the 20 μl reverse transcription reaction was completed by adding 0.5 μl of SuperScript II as described below, the concentration of each reagent in the master mix was: 50 mM Tris-HCl (pH 8.0), 75 mM KCl, 0.5 mM dNTP Solution Mix, 10 mM DTT, 2% (v/v) deionized formamide (Millipore), 10 ng/μl ET SSB (New England Biolabs), 3 mM MnCl_2_ (Fisher Scientific), 0.1% (v/v) Triton X-100, and 5 U/μl SuperScript II (Invitrogen). As described in the original SHAPE-MaP procedure, it is crucial to add MnCl_2_ stock to the master mix immediately before performing reverse transcription because the manganese will begin to oxidize and precipitate in this solution^74^. Samples were placed on a pre-heated 42 °C thermal cycler for 2 min before 0.5 μl of SuperScript II was added to complete the master mix. The reverse transcription reaction was incubated at 42 °C for 50 min, and then at 70 °C for 15 min to heat inactivate SuperScript II. Samples were cooled to 12 °C, 1.25 U of RNase H (New England Biolabs) and 12.5 U of RNase I_f_ (New England Biolabs) were added to each sample, and the samples were incubated at 37 °C for 20 min and then at 70 °C for 20 min to heat inactivate the RNases. The samples were briefly spun down in a mini-centrifuge and placed on a magnetic stand. The supernatant was aspirated and discarded, and the bead pellet was washed with 75 μl Storage Buffer (10 mM Tris-HCl (pH 8.0) and 0.05% (v/v) Triton X-100). The beads were resuspended in 25 μl of Storage Buffer, transferred to screw-cap tubes with an O-ring, and stored at −20 °C.

### Test amplification of TECprobe-VL libraries

The number of PCR amplification cycles needed for TECprobe libraries was determined by performing a test amplification adapted from Mahat et al.^75^. Briefly, 14 μl of PCR master mix was added to 6 μl of a 16-fold dilution of the bead-bound cDNA libraries, such that the final concentration of components in the 20 μl PCR was: 1X Q5 Reaction Buffer (New England Biolabs), 1X Q5 High GC Enhancer (New England Biolabs), 200 μM dNTP Solution Mix, 250 nM RPIX Forward Primer (Table S6), 250 nM dRP1_NoMod.R Reverse Primer (Table S6), 10 nM RPIX_SC1_Bridge (Table S6), and 0.02 U/μl Q5 High-Fidelity DNA Polymerase. Amplification was performed for 21 or 25 cycles, at an annealing temperature of 62 °C and an extension time of 20 s. 20 μl of each supernatant was run on native TBE-polyacrylamide gels to assess both the fragment size distribution of the libraries and to determine the appropriate number of cycles for amplification of libraries for high-throughput sequencing.

### Preparation of dsDNA libraries for sequencing

Amplification of cDNA libraries for high throughput sequencing was performed by preparing separate 50 μl PCRs for each (+) and (-) sample that contained 1X Q5 Reaction Buffer, 1X Q5 High GC Enhancer, 200 μM dNTP Solution Mix, 250 nM RPI Indexing Primer (Table S6), 250 nM dRP1_NoMod.R Reverse Primer (Table S6), 10 nM SC1Brdg_MINUS or SC1Brdg_PLUS channel barcode oligo (Table S6), 12 μl of bead-bound cDNA library, and 0.02 U/μl Q5 High-Fidelity DNA Polymerase. Amplification was performed as indicated above, using the number of cycles determined by the test amplification. Supernatants from completed PCRs were each mixed with 100 μl of SPRI beads and purified as described in *SPRI bead purification of DNA*. DNA was eluted into 20 μl of 10 mM Tris-HCl (pH 8.0), mixed with 40 μl of SPRI beads, and purified as described in *SPRI bead purification of DNA* a second time. Twice-purified DNA was eluted into 10 μl of 10 mM Tris-HCl (pH 8.0) and quantified using the Qubit dsDNA HS Assay Kit (Invitrogen) with a Qubit 4 Fluorometer. Molarity was estimated using the length distribution observed during test amplification.

### High-throughput DNA sequencing

Sequencing of chemically probed libraries was performed by Novogene Co. on an Illumina HiSeq X Ten System using 2×150 PE reads with 10% PhiX spike in. TECprobe-VL libraries were sequenced at a depth of ∼60 to ∼80 million PE reads.

### Sequencing read pre-processing using cotrans_preprocessor

All custom software are freely available at https://github.com/e-strobel-lab/TECtools/releases/tag/v1.2.0 or https://github.com/e-strobel-lab/TECprobe_visualization/releases/tag/v1.0.0.

Briefly, 3’ end targets and intermediate transcript targets were generated by running `cotrans_preprocessor` in `MAKE_3pEND_TARGETS` mode. Sequencing reads were then processed by running `cotrans_preprocessor` in `PROCESS_MULTI` mode, which manages adapter trimming using fastp^76^ and demultiplexes sequencing reads by 3’ end identity and channel (modified or untreated). A shell script to run ShapeMapper2^77^ for every intermediate transcript was generated by running `cotrans_preprocessor` in `MAKE_RUN_SCRIPT` mode. Sequencing read alignment and reactivity calculation was then performed by ShapeMapper2. After comparing replicate data sets that were analyzed individually, TECprobe-VL replicate data were merged and normalized using `process_TECprobeVL_profiles`. Reactivity matrix heatmaps were generated using the `generate_cotrans_heatmap` script.

### Modeling of non-native RNA secondary structures

Non-native intermediate glycine riboswitch structures were modeled as described previously^68^: For each transcript of interest, the 3’-most 11 nt were forced to be single-stranded to account for the *E*. *coli* RNAP footprint and possible base pairing configurations for the remaining transcript were generated using the RNAstructure v6.4 stochastic command with default settings^78^. From this set of structures, putative folding intermediates were manually selected based on the agreement of the predicted structure with reactivity data and by assessing which nucleotides had recently emerged from RNAP at the transcript length when the intermediate is first observed and when the intermediate rearranges into a subsequent structure.

### Computational modeling

To model the 3D structures of the GTR, we employed the Vfold-Pipeline^79^, a template-based model for RNA 3D structure prediction, to generate the initial structures for simulation. Subsequently, we conducted coarse-grained molecular dynamics (CGMD) simulations utilizing the IsRNA model, applying 2D structure constraints (predicted based on DMS-seq data) and tertiary interaction constraints (A-minor interactions within α and β, and the kink-turn motif). IsRNA serves as a statistical potential and CGMD-based model for 3D structure prediction, in which each nucleotide is coarse-grained into five beads to enhance efficiency^50–52^. Replica-exchange MD was conducted for 50 ns at ten varying temperatures, with a timestep of 1 fs. The low-energy structures were clustered according to structural similarity (RMSD), and the centroid structure of each cluster was identified and ranked based on cluster size. The top-ranked models were obtained as the predicted structures. Alignment of the three top-ranked models reveals highly dynamic terminator and antiterminator domains (Figures 6D and 6E). For the antiterminator, we modeled the two glycine’s docked into the two binding pockets through a local refinement of the glycine binding pocket based on the solved GTR structure (PDB 3OWI).

Furthermore, we employed IsRNA to run CGMD for the ΔP0 variant and the WT from the same initial structure (the 3D model of antiterminator, with the P0 removed for the ΔP0 variant). The simulations are conducted for 50 ns at a temperature of 300 K, with only 2D structure constraints applied and tertiary interaction constraints removed to examine their impact on structural stability. As shown in Figure S27A, B, the ΔP0 variant is unfolded after 20 ns of simulation and becomes fully extended by the end of the simulation, while WT remains relatively compact throughout the simulation. At the end of the simulation, the RMSD changes for the ΔP0 variant and WT are 47.49 Å and 29.79 Å, respectively. Here the RMSDs are calculated based on the pseudoatoms (beads) in CGMD, excluding the antiterminator domain. The result suggests a lower folding stability of ΔP0 variant than WT. We also modeled (predicted) the 3D structures of the ΔP0 variant, hypothesizing that they function as the antiterminator and terminator, by conducting 50 ns of replica exchange MD. As shown in Fig. S27C, the simulations result in two significantly different structures compared to the WT structures, with RMSD values of 21.62 Å and 16.00 Å to the corresponding WT structures, respectively.

To investigate how each binding pocket contributes to the stability of the GTR, we conducted four CGMD simulations using IsRNA: WT, without-glycine-constraints, without-glycine-constraint-1, and without-glycine-constraint-2. For WT, a folding constraint is applied to the two glycine binding pockets to maintain the ligand-bound conformation, which was obtained from the solved GTR structure (PDB 3OWI). For without-glycine-constraints, the simulation was conducted without folding constraints at the two binding pockets. For without-glycine-constraint-1 and without-glycine-constraint-2, the simulations were conducted without folding constraints at binding pocket in aptamer 1 and aptamer 2, respectively. Starting from the same initial structure, replica exchange molecular dynamics simulations were performed for 50 ns. As a result, the top-ranked model of WT is notably closer to the initial structure than without-glycine-constraints, with RMSD values of 4.5 Å for WT and 12.9 Å for without-glycine-constraints. Moreover, the number of structures in the top cluster is 249 for WT and 115 for without-glycine-constraints, respectively. This observation suggests that the without-glycine-constraints has larger structural changes and lower folding stability (Figure S28A). Furthermore, without-glycine-constraint-1 causes significant disruption in the aptamer 1 region, rather than in the aptamer 2 region, as evidenced by RMSD changes of 5.2 Å in the aptamer 1 region and 2.8 Å in the aptamer 2 region. In contrast, without-glycine-constraint-2 significantly disrupts the aptamer 2 region, resulting in RMSD changes of 3.5 Å and 6.5 Å in the aptamer 1 and aptamer 2 regions, respectively (Figures S28B, C). These results indicate that each glycine binding pocket primarily contributes to the stability of its respective aptamer.

