## Supplemental Information for "Co-transcriptional folding orchestrates sequential multi-effector sensing by a glycine tandem riboswitch"

#### Supplemental information Index:

Supplementary Notes 1

Figures S1-S28

Table S1-S10

**Supplementary Note 1. Co-transcriptional folding of the *B. subtilis gcvT* glycine riboswitch**

The *Bsu*-GTR co-transcriptional RNA folding pathway is populated by several non-native intermediate hairpins that are resolved as native structures fold. Aptamer 1 folding begins with the formation of intermediate hairpin 1 (IH1), which is detected at ~+50 as decreased reactivity at C24 and C25 as they and U26 form base pairs with a stretch of four G bases that span positions 15 to 18 (Figure S22A and S23A). At ~+72, decreased reactivity at C29, C30, C39, C46, and A47 is consistent with formation of both the upper and lower P2.1 stem (Figure S22B and S23B). The reactivity of C46 and A47 remains moderate at this transcript length, which implies that either some nascent transcripts do not form the lower P2.1 stem or that the lower P2.1 stem is not yet stably formed. Consistent with the latter interpretation, the reactivity of C46 and A47 increases from +77 to +84 which indicates that the U26:A47 and G27:C46 base pairs have been broken and the lower P2.1 stem disrupted (Figure S22C and S23C). This increased reactivity at C46 and A47 occurs in coordination with decreased reactivity at C52, C53, A55, and A56 (Figure S23C). While this reduction in reactivity reaches a minimum at +84, it begins at +75 when ~61 nucleotides have emerged from RNAP. At this transcript length, C52 and C53 can pair with G58 and G57 in intermediate hairpin 2 (IH2), respectively (Figure S22C). Consistent with this interpretation, the reactivity of C50, which is predicted to pair with G60 in IH2, also decreases slightly (Figure S23C). However, C61, which is predicted to pair with G49, remains reactive to DMS. This suggest that even if the G49:C61 base pair does form, it is not stable.

The reactivity of C46 and A47 decreases again at ~+95 indicating that the lower P2.1 stem has folded a second time (Figure S22D and S23D). As above, the reactivity of C46 and A47 remains moderate at this transcript length, suggesting that the lower P2 stem is not yet stably folded. In coordination with lower P2 stem folding, increased reactivity at C50, A55, and A56 suggests that IH2 has been disrupted (Figure S23C). The disruption of IH2 is likely driven by the simultaneous formation of a third intermediate hairpin (IH3) that overlaps with most IH2 nucleotides but does not overlap with P2 (Figure S22D). The primary evidence for the formation of IH3 is decreased reactivity at C65 and C68, which are predicted to pair with G57 and G54, respectively (Figure S23D). At ~95, decreased reactivity at C65 is unlikely to be caused by formation of the C65:G72 P3b.1 base pair because the reactivity of A64, which pairs with U73 in P3b.1, remains approximately constant (Figure S23D). Despite exhibiting a substantial reactivity decrease at ~+95, C65 and C68 remain moderately reactive to DMS (Figure S23D). In contrast, the reactivity of C52 and C53, which are predicted to pair with G70 and G69 in IH3, remains low (Figure S22D and S23C). This suggest that IH3 may not fold in all cases and that C52 and C53 form other base pairs if IH3 does not fold.

The reactivity of C78, C80, and C82 decreases from ~+100 to ~+106 (Figure S23E). While these nucleotides form base pairs with G60, G58, and G57 in the native P3a.1 stem, the reactivity decrease that begins at ~+100 is unlikely to be caused by P3a.1 folding because the reactivity of A59, which pairs with U79 in P3a.1, remains approximately constant (Figure S23E). A coordinated decrease in the reactivity of A71 across these same transcript lengths suggests the formation of a fourth intermediate hairpin comprised of G70:C80, A71:U79, and G72:C78 base pairs (Figure S22E and S23E). Consistent with this interpretation, the reactivity of A55 and A56 in the upstream segment of binding site 1 remains approximately constant from ~+100 to ~+106, indicating that binding site 1 has not yet folded (Figure S23F).

P1 folding is observed as decreased reactivity at A10, C12, A13, and A14, which participate in P1.1 base pairs, and occurs in two steps (Figure S22F and S23F). The first step occurs at +111 and is coordinated with decreased reactivity at C46 and A47 in P2.1, C86 in P3.1, and A59, C78, C80, and C82 in P3a.1 (Figure 23E, F). The second step occurs from +115 to +118, and is coordinated with a further decrease in the reactivity of C46 and A47 in P2.1 and C86 in P3.1 and increased reactivity at A55, A56, and A83 in binding site 1 (Figure 24F). When glycine is present, glycine-dependent reactivity changes in these and other nucleotides are first detected from +115 to +118, indicating that the second step of P1 folding is coordinated with complete folding of the first aptamer (Figure S8).

P0 folding is observed from +118 to +130 as decreased reactivity at C2, C105, and C107 as the G1:C107, C2:G106, and G3:C105 base pairs form (Figure S23G). The reactivity of these nucleotides remains approximately constant for most of the remaining the folding pathway, but increases slightly when intermediate hairpin 7 (IH7) within the second aptamer domain is folded. Notably, the position of the C126 pause could facilitate Aptamer 1 and P0 folding and glycine binding (Figure S24A).

Like Aptamer 1, Aptamer 2 folding begins with the formation of an intermediate hairpin. Intermediate hairpin 5 (IH5) is detected as decreased reactivity at C110 and C112, which pair with G119 and G117, respectively (Figure S25A and S26A). At ~+157, P2.2 folding is detected as decreased reactivity at C128, C134, and nucleotides 136-140, all of which participate in P2.2 base pairs (Figure S25B and S26B). A131 in the P2.2 loop remains reactive (Figure S26B). At ~+172, a sixth intermediate hairpin (IH6) may fold, however the evidence for this structure is limited because the IH6 stem contains only one DMS-reactive nucleotide (Figure S25C). At +178, a coordinated decrease in the reactivity of nucleotides 141 to 145 corresponds to the formation of IH7, which extends P2.2 (Figure S25D and S26C). A116 is predicted to form a non-canonical pair with A142 but remains highly reactive, indicating that its Watson-Crick face is exposed (Figure S25D). P3b.2 folding is then detected at +194 as decreased reactivity at C156, C161, and nucleotides 167 to 170, which participate in P3b.2 base pairs (Figure S25E and S26D). A164 in the P3b.2 loop remains reactive (Figure S26D).

The observation of glycine-dependent reactivity differences at +211 indicates that Aptamer 2 can fully fold as soon as it has fully emerged from RNAP in some nascent transcripts (Figure S8). However, reactivity changes that are associated with the formation of inter-aptamer contacts continue until ~+220 (Figure S8). This suggests that the synthesis of additional nascent RNA is necessary to facilitate contacts between aptamers 1 and 2, presumably due to steric conflicts with RNAP. The signatures of P1.2 folding are relatively weak because upstream P1.2 nucleotides are engaged in IH7 base pairs until P1.2 folds and downstream P1.2 nucleotides likely begin to form P1.2 base pairs as soon as they emerge from RNAP. Nonetheless, decreased reactivity at C110, C112, and C192 from ~+211 to ~+216 is consistent with P1.2 folding (Figure S25F and

S26A). P3.2 and P3a.2 folding is detected as decreased reactivity at  C143, C144, A146, A147, C176, C180, and A181 from ~+214 to ~+220 (Figure S26C, E).

Notably, the +225 pause is located in a window from ~+220 to ~+229 within which the glycine aptamers can bind glycine and inter-aptamer contacts can form (Figure S8). This suggests that the +225 pause contributes to glycine riboswitch function by allowing time for these folding events to occur before the transcription terminator is transcribed. The position of the +225 pause may permit several base pairs of the terminator hairpin to fold within the RNA exit channel of RNAP, which at this location would likely stabilize the pause (Figure S24C). While the reactivity of C203 and A204 decreases slightly as RNAP approaches +225, these nucleotides remain highly reactive until terminator folding occurs from ~+228 to ~+236 (Figure S24D). This indicates that if the apical base pairs of the terminator hairpin function has a pause hairpin at +225, these base pairs do not stably form until additional terminator hairpin base pairs are able to form.

### Supplemental Figures

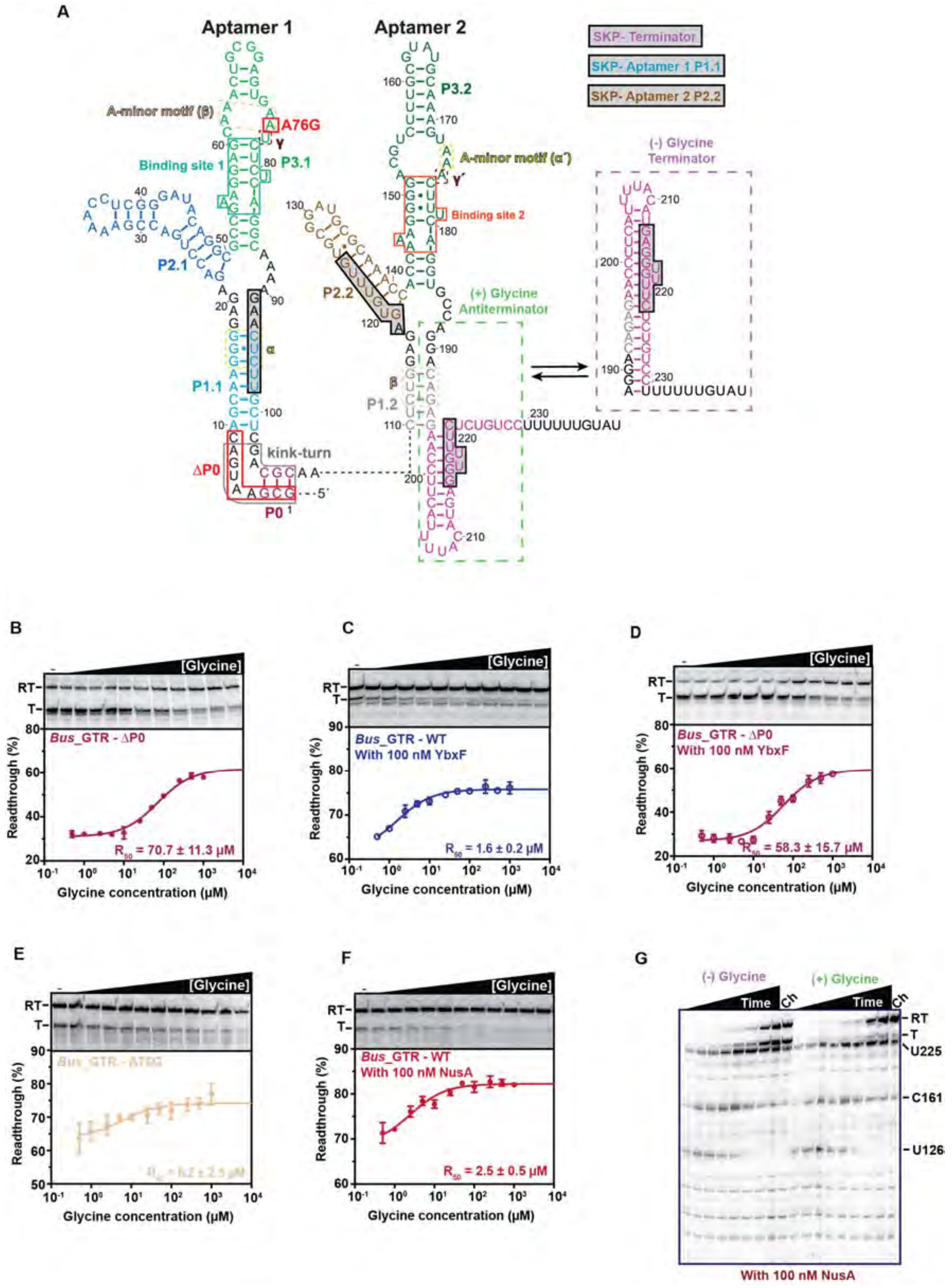

**Supplemental Figure 1. Secondary structure of the *Bsu*-GTR and transcription gels.**

(**A**) The full-length riboswitch sequence color coded with binding site 1 boxed in green-blue, binding site 2 boxed in orange and inter-aptamer contacts shown as dashed boxes (α, lime yellow; β, peach; and γ, dark red). The regions targeted by SiM-KARTS are shaded in light gray boxes. Perturbations are indicated by red boxes. (**B-F**) Glycine dose-response curves for the ΔP0 variant (B), WT riboswitch with 100 nM YbxF (C), ΔP0 variant with 100 nM YbxF (D), A76G variant (E) and WT riboswitch with 100 nM NusA (F) measured by single-round *in vitro* transcription. Representative gels are shown above each plot. Error bars represent the SD from independent replicates (n=3). (**G**) Time-resolved single-round *in vitro* transcription of the *B. subtilis* glycine riboswitch in the presence of 100 nM NusA and in the presence or absence of glycine. Paused (U126, C161, U225), terminated (T) and readthrough (RT) transcripts are indicated.

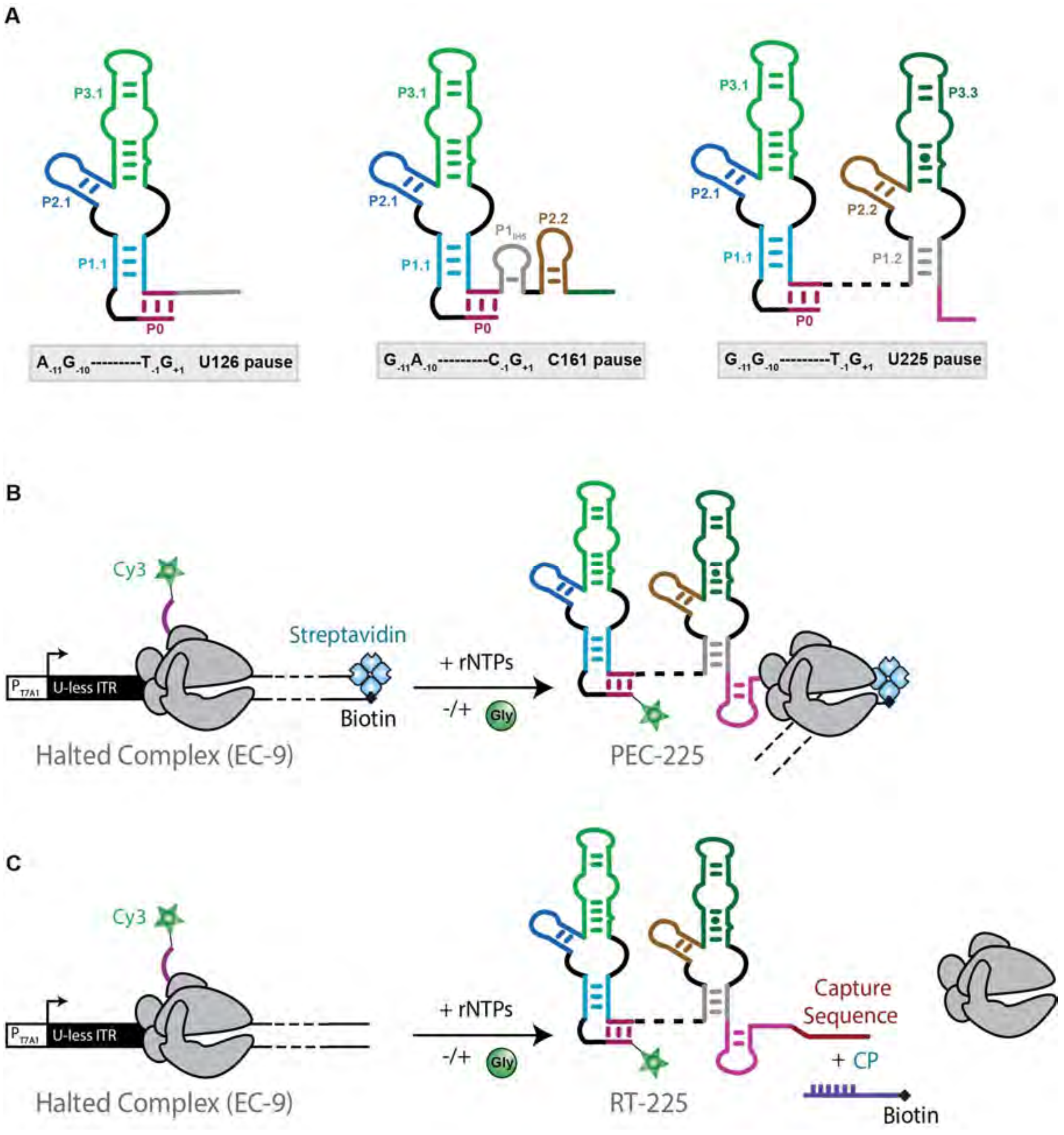

**Supplemental Figure 2. Pause secondary structure and how to make PEC's and release transcripts.** (**A**) Cartoon depictions of secondary structures at pauses U126, C161 and U226 with the identity of nucleotides at positions known to be determinants of the consensus pause shown below. (**B-C**) Fluorescently-labeled nascent RNA transcripts are transcribed *in vitro* using *E. coli* RNAP. The halted complex (EC-9) is prepared through the addition of a dinucleotide labeled with Cy3 (ApU-Cy3) and UTP deprivation (ATP/CTP/GTP) to halt the RNAP at the end of the U-less ITR. (B) Transcription will continue until RNAP reaches the biotin-streptavidin roadblock forming the Paused Elongation Complex (PEC). (C) Once the transcription is completed, a 5′-biotinylated capture probe is hybridized to the released transcript (RT-225) for subsequent immobilization on the microscope slide.

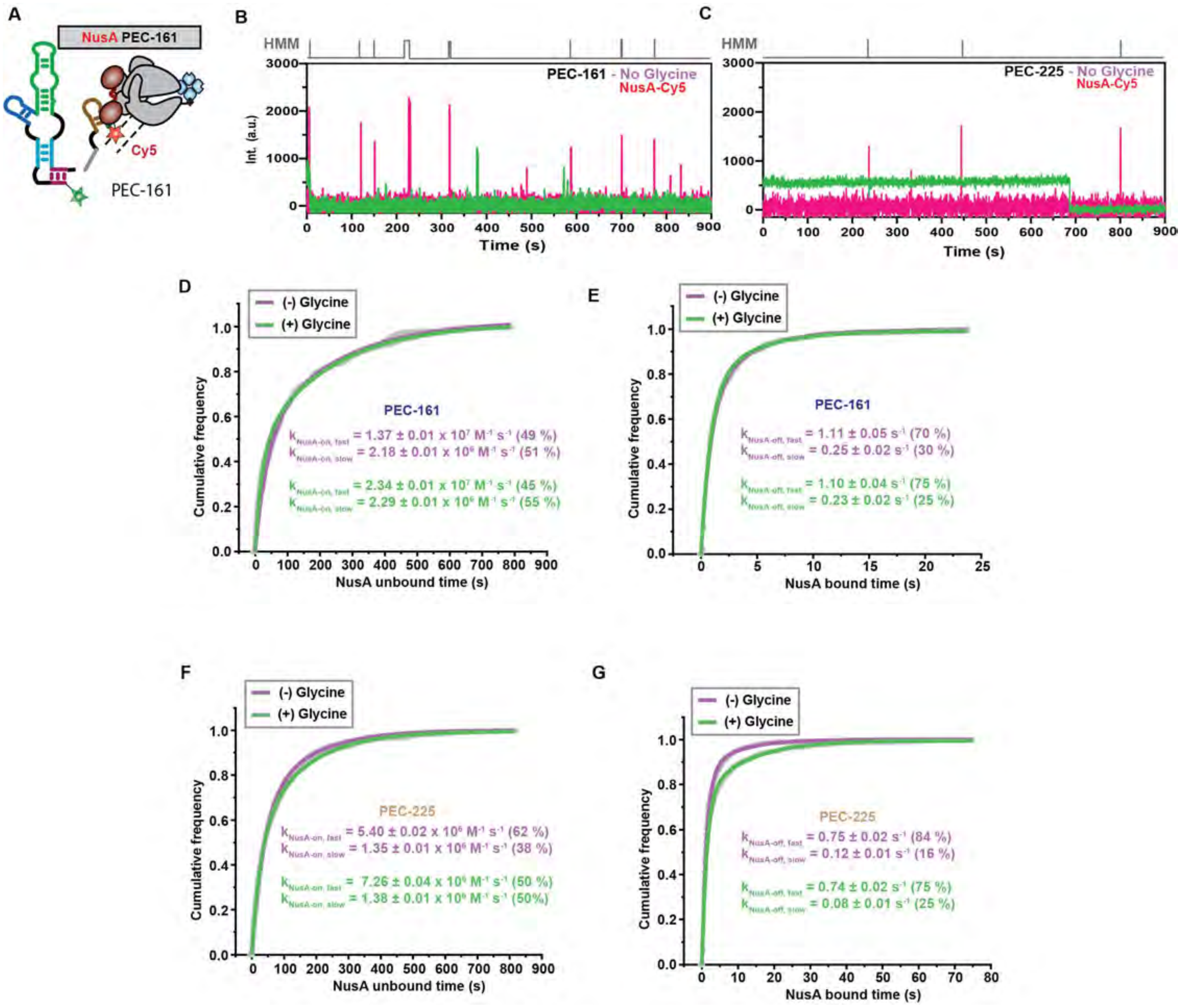

**Supplemental Figure 3. NusA binding kinetics curves.**

(**A**) Diagram of PEC-161 in complex with NusA-Cy5. (**B**, **C**) Representative single-molecule trajectories showing NusA-Cy5 binding (pink) to PEC-161 without (B) and with (C) 1 mM glycine. (**D, E**) Plots displaying the cumulative unbound (D) and bound (E) dwell times of NusA-Cy5 binding in the absence (purple) and presence (green) of 1 mM glycine in the context of PEC-161. The association ($k_{on}$) and dissociation ($k_{off}$) rate constants of NusA-Cy5 binding are indicated on the plot. The reported errors are the error of the fit. Total number of molecules analyzed for each condition is: (−) Glycine = 116; (+) Glycine = 220. (**F**, **G**) Plots displaying the cumulative unbound (F) and bound (G) dwell times of NusA-Cy5 binding in the absence (purple) and presence (green) of 1 mM glycine in the context of PEC-225. The association ($k_{on}$) and dissociation ($k_{off}$) rate constants of NusA-Cy5 binding are indicated on the plot. The reported errors are the error of the fit. The total number of molecules analyzed for each condition is: (−) Glycine = 237; (+) Glycine = 248.

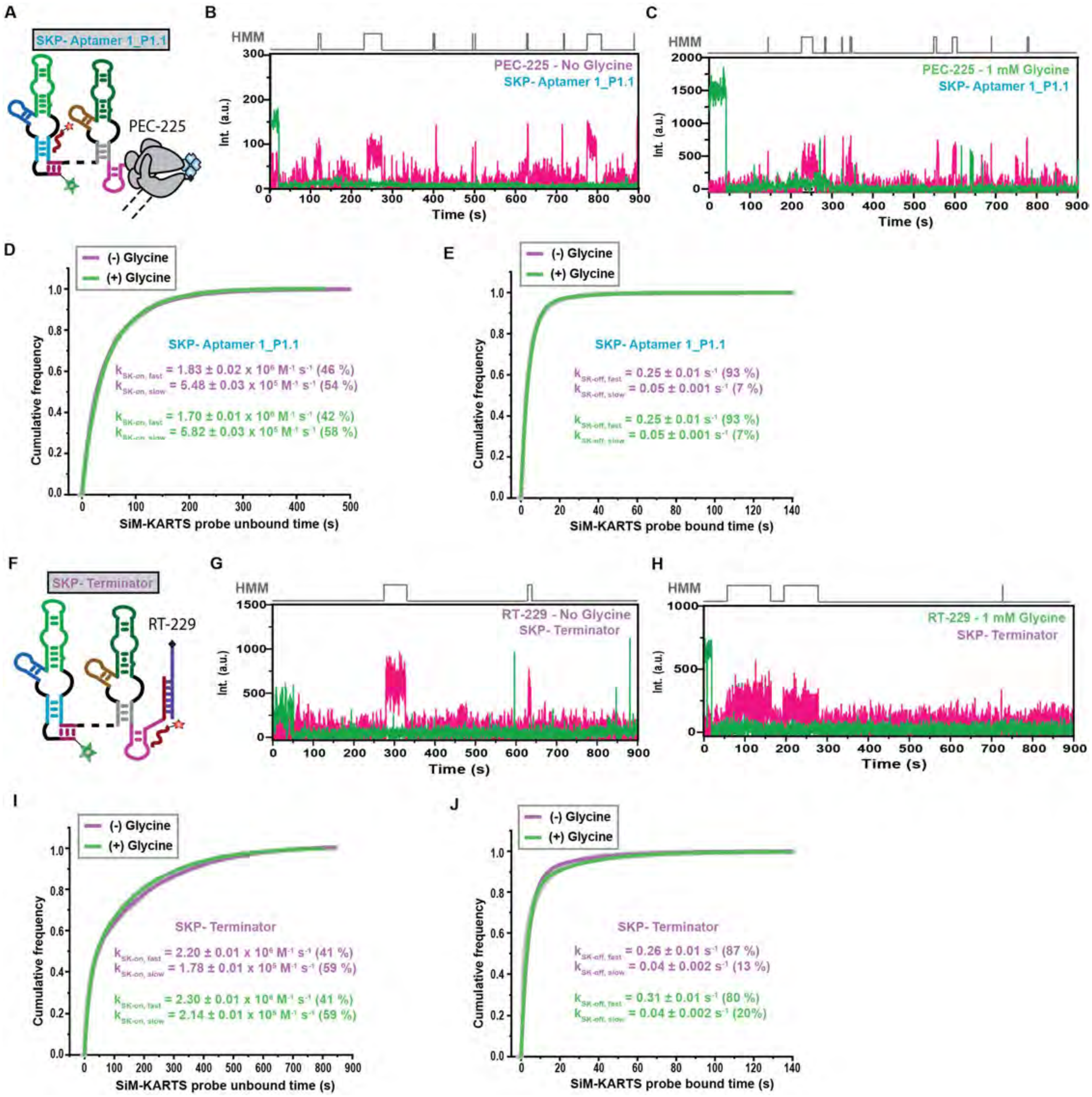

**Supplemental Figure 4. SiM-KARTS probing P1.1 and terminator hairpin.**

(**A**) Diagram of SiM-KARTS probe targeting aptamer 1_P1.1 in PEC-225 (**B**, **C**) Representative single-molecule trajectories showing the SiM-KARTS probe binding (pink) to PEC-225 without (B) and with (C) 1 mM glycine. (**D**, **E**) Plots displaying the cumulative unbound (D) and bound (E) dwell times of SiM-KARTS probe binding in the absence (purple) and presence (green) of 1 mM glycine in the context of PEC-161. The association ($k_{on}$) and dissociation ($k_{off}$) rate constants of the SiM-KARTS probe binding are indicated on the plot. The reported errors are the error of the fit. The total number of molecules analyzed for each condition is:

(−) Glycine = 91; (+) Glycine = 99. (**F**) Diagram of SiM-KARTS probe targeting the terminator hairpin in RT-229 (**G**, **H**) Representative single-molecule trajectories showing SiM-KARTS probe binding (pink) to PEC-225 without (G) and with (H) 1 mM glycine. Plots displaying the cumulative unbound (I) and bound (J) dwell times of SiM-KARTS probe binding in the absence (purple) and presence (green) of 1 mM glycine in the context of PEC-161. The association ($k_{on}$) and dissociation ($k_{off}$) rate constants of SiM-KARTS probe binding are indicated on the plot. The reported errors are the error of the fit. The total number of molecules analyzed for each condition is: (−) Glycine = 367; (+) Glycine = 413.

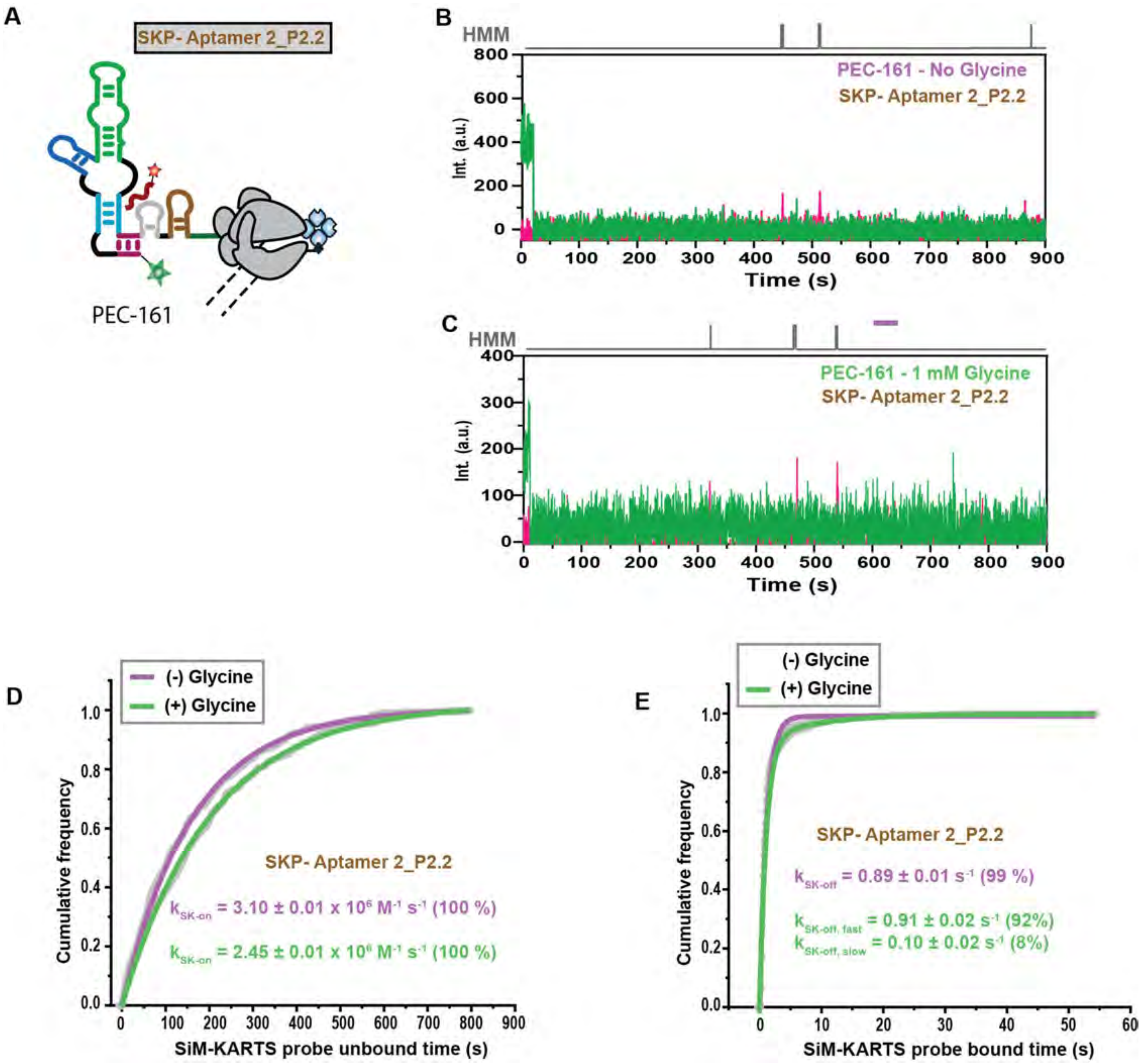

**Supplemental Figure 5. SiM-KARTS probing aptamer 2_P2.2.**

(**A**) Diagram of SiM-KARTS probe targeting aptamer 2_P2.2 in PEC-161 (**B**, **C**) Representative single-molecule trajectories showing the SiM-KARTS probe binding (pink) to PEC-161 without (B) and with (C) 1 mM glycine. (**D**, **E**) Plots displaying the cumulative unbound (D) and bound (E) dwell times of the SiM-KARTS probe binding in the absence (purple) and presence (green) of 1 mM glycine in the context of PEC-161. The association ($k_{on}$) and dissociation ($k_{off}$) rate constants of SiM-KARTS probe binding are indicated on the plot. The reported errors are the error of the fit. The total number of molecules analyzed for each condition is: (−) Glycine = 192; (+) Glycine = 197.

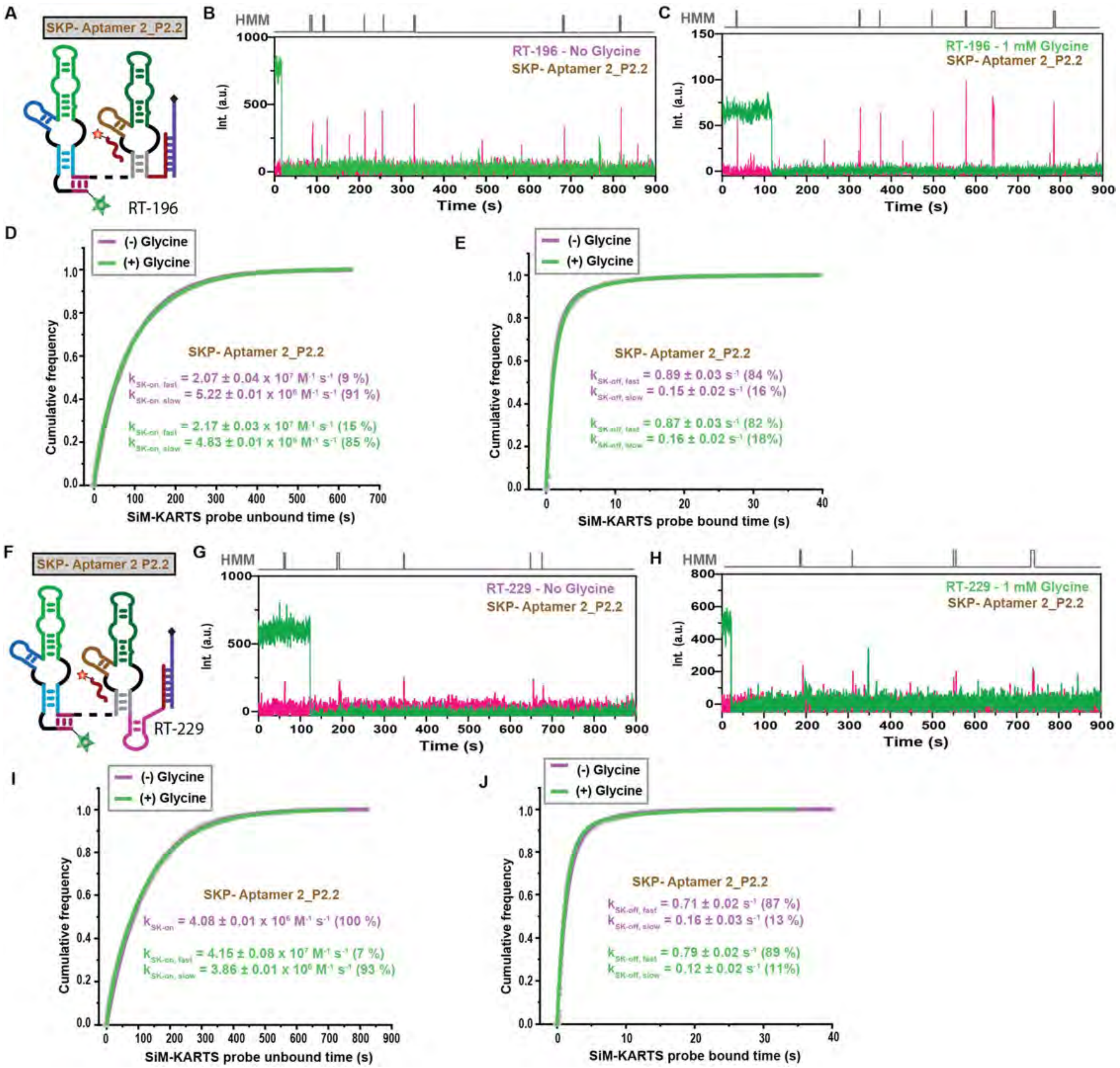

**Supplemental Figure 6. SiM-KARTS probing aptamer 2_P2.2 in release transcripts.**
(**A**) Diagram of SiM-KARTS probe targeting aptamer 2_P2.2 in RT-196. (**B**-**C**) Representative single-molecule trajectories showing the SiM-KARTS probe binding (pink) to PEC-196 without (B) and with (C) 1 mM glycine. (**D**, **E**) Plots displaying the cumulative unbound (D) and bound (E) dwell times of the SiM-KARTS probe binding in the absence (purple) and presence (green) of 1 mM glycine in the context of PEC-161. The association ($k_{on}$) and dissociation ($k_{off}$) rate constants of SiM-KARTS probe binding are indicated on the plot. The reported errors are the error of the fit. The total number of molecules analyzed for each condition is: (−) Glycine = 210; (+) Glycine = 358. (**F**) Diagram of SiM-KARTS probe targeting aptamer 2_P2.2 in RT-229. (**G**, **H**) Representative single-molecule trajectories showing the SiM-KARTS probe binding (pink) to RT-229 without

(G) and with (H) 1 mM glycine. (**I**, **J**) Plots displaying the cumulative unbound (I) and bound (J) dwell times of SiM-KARTS probe binding in the absence (purple) and presence (green) of 1 mM glycine in the context of PEC-161. The association ($k_{on}$) and dissociation ($k_{off}$) rate constants of SiM-KARTS probe binding are indicated on the plot. The reported errors are the error of the fit. The total number of molecules analyzed for each condition is: (−) Glycine = 281; (+) Glycine = 270. All error bars are the SD of the mean of independent replicates.

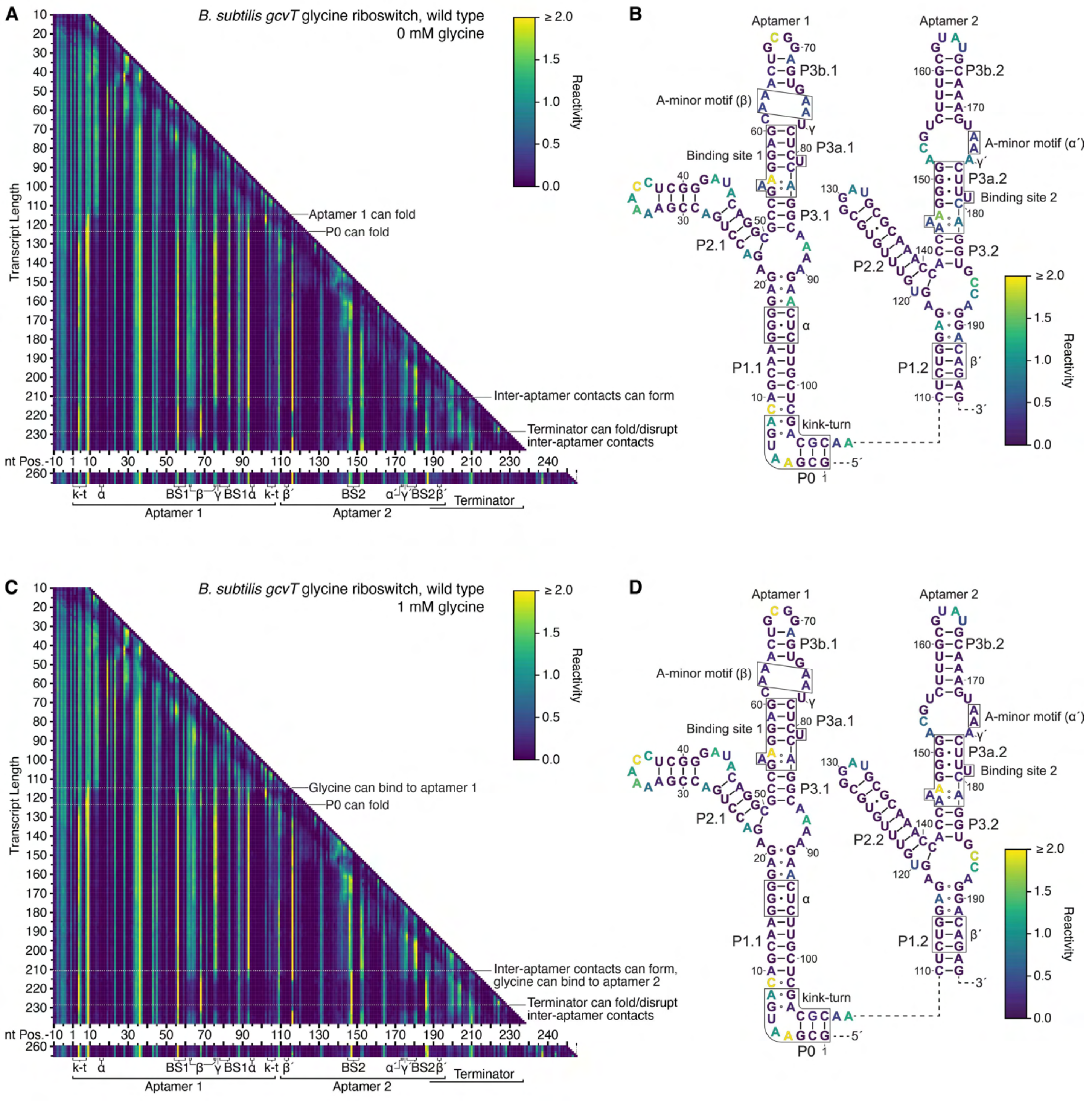

**Supplementary Figure 7. Cotranscriptional DMS structure probing of the wild type *B. subtilis gcvT* riboswitch.**

(**A**, **C**) TECprobe-VL DMS reactivity matrices for the wild type *B. subtilis gcvT* riboswitch in the absence (A) and presence (C) of 1 mM glycine. Transcripts 239-259, which were not enriched, are excluded. (**B**, **D**) *B. subtilis gcvT* glycine aptamer secondary structures colored by the TECprobe-VL DMS reactivity of the 223 nt transcript in the absence (B) and presence (D) of 1 mM glycine. Data are from two independent replicates that were processed separately and merged.

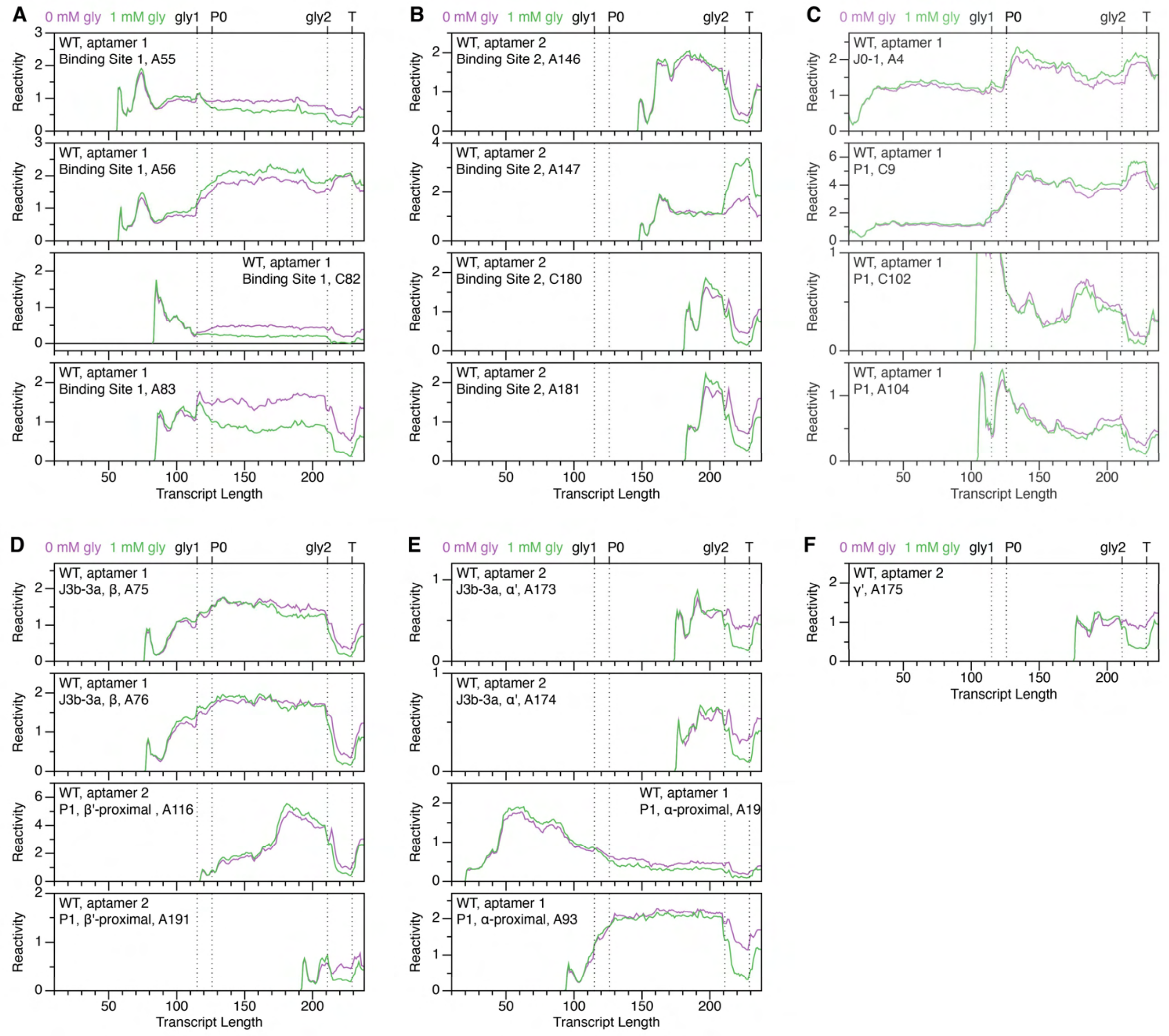

**Supplementary Figure 8. DMS reactivity trajectories for select nucleotides of the wild type *B. subtilis gcvT* riboswitch.** (**A**-**F**) TECprobe-VL DMS reactivity trajectories for nucleotides in or proximal to binding site 1 (A), binding site 2 (B), the kink-turn motif (C), β and β’ (D), α and α’ (E), and γ ‘ (F). Data are from the experiment shown in Supplementary Figure 7.

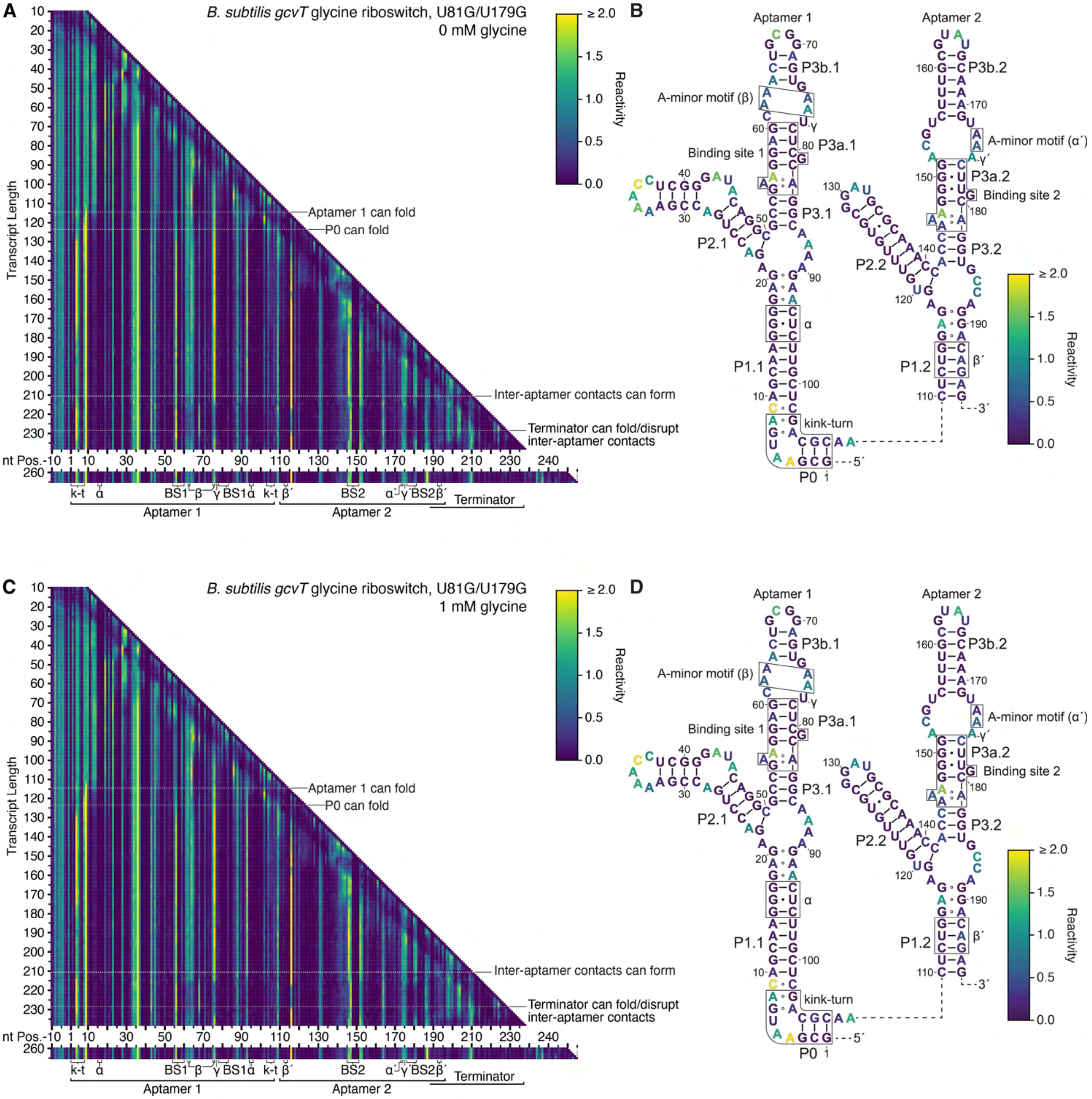

**Supplementary Figure 9. Cotranscriptional DMS structure probing of the *B. subtilis gcvT* riboswitch U81G, U179G variant.** (**A**, **C**) TECprobe-VL DMS reactivity matrices for the *B. subtilis gcvT* riboswitch U81G, U179G variant in the absence (A) and presence (C) of 1 mM glycine. Transcripts 239-259, which were not enriched, are excluded. (**B**, **D**) *B. subtilis gcvT* glycine aptamer U81G, U179G variant secondary structures colored by the TECprobe-VL DMS reactivity of the 223 nt transcript in the absence (B) and presence (D) of 1 mM glycine. Data are from two independent replicates that were processed separately and merged.

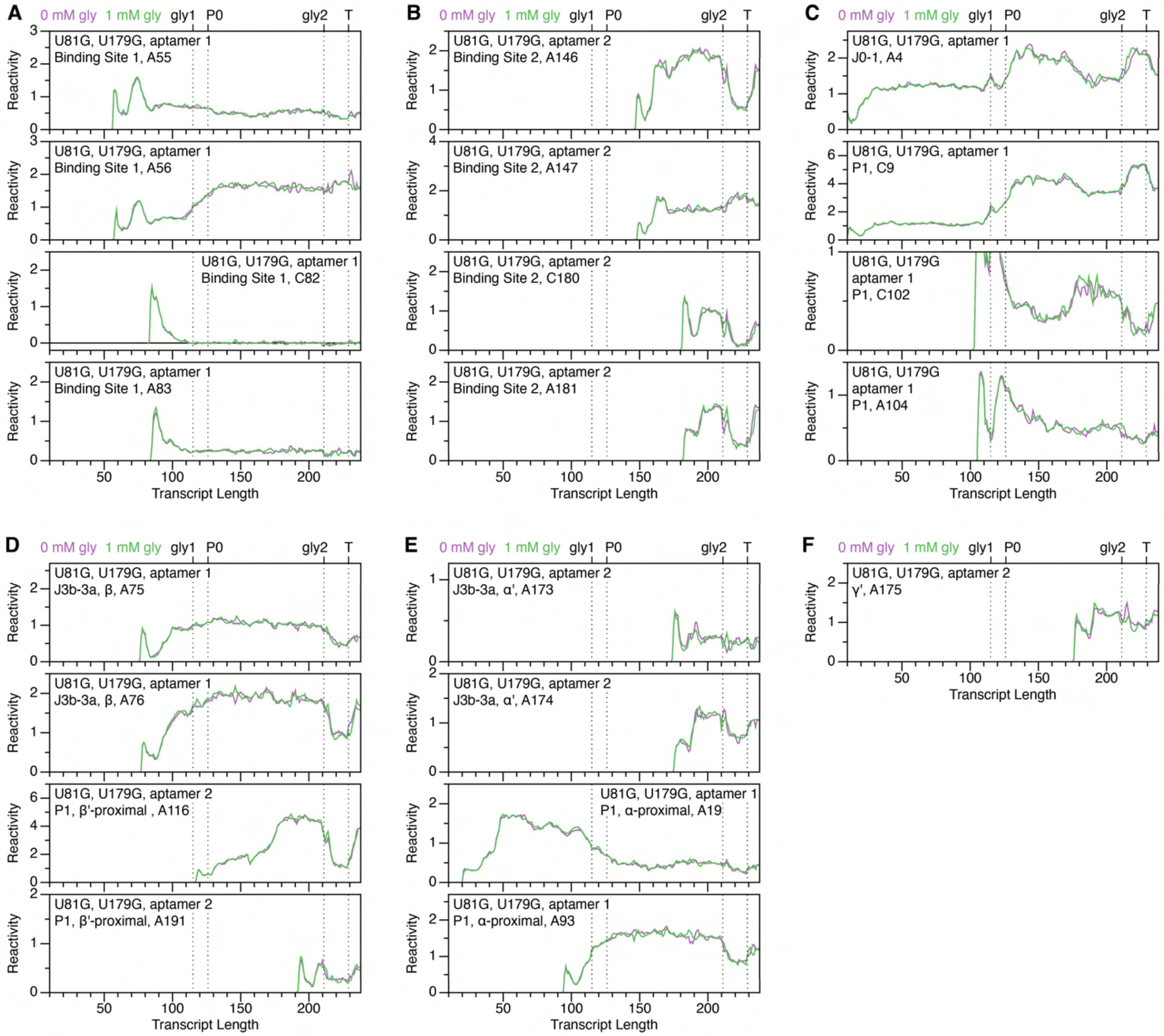

**Supplementary Figure 10. DMS reactivity trajectories for select nucleotides of the *B. subtilis gcvT* riboswitch U81G, U179G variant.** (**A**-**F**) TECprobe-VL DMS reactivity trajectories for nucleotides in or proximal to binding site 1 (A), binding site 2 (B), the kink-turn motif (C), β and β' (D), α and α' (E), and γ' (F). Data are from the experiment shown in Supplementary Figure 9.

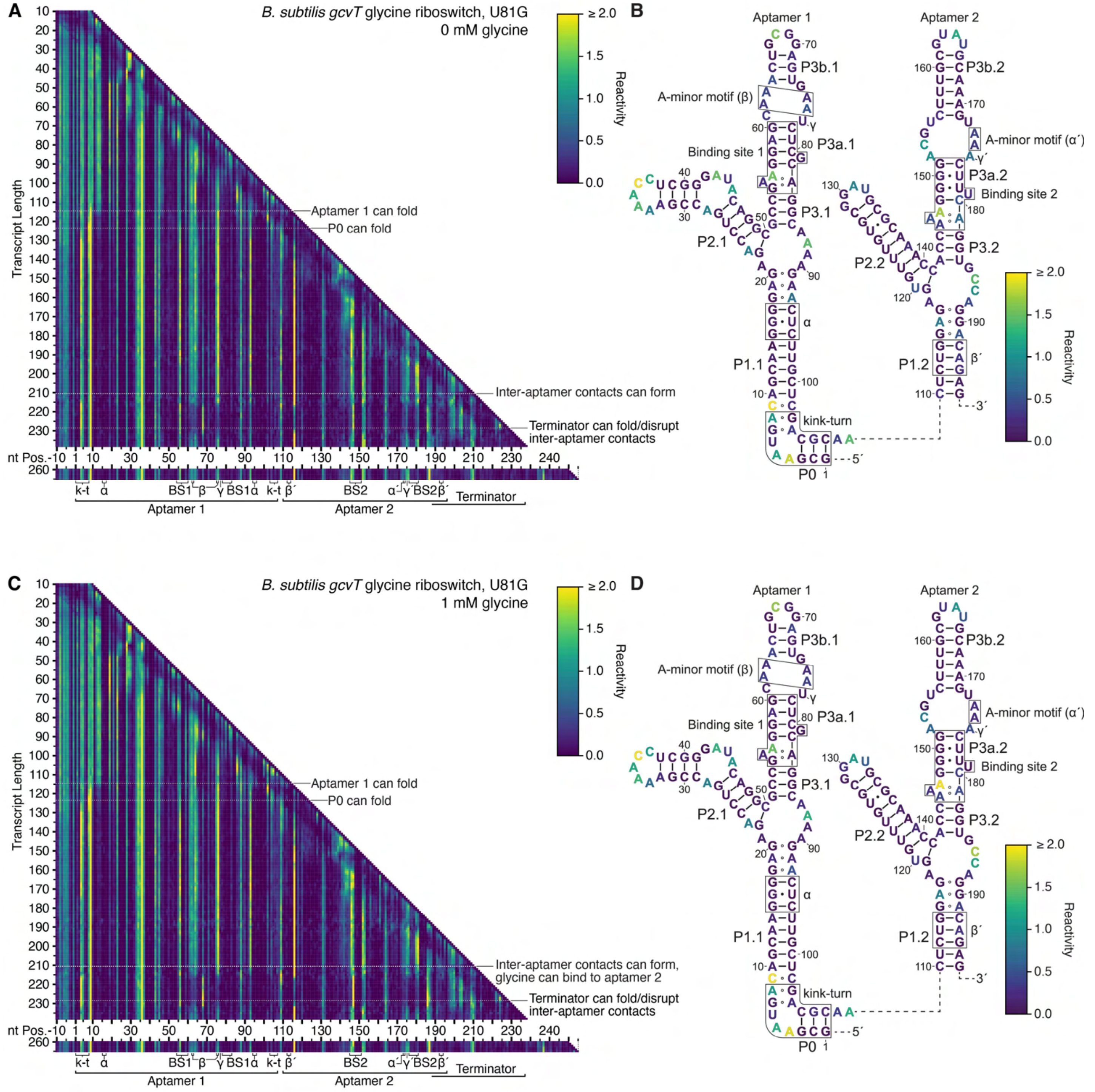

**Supplementary Figure 11. Cotranscriptional DMS structure probing of the *B. subtilis gcvT* riboswitch U81G variant.** (**A**, **C**) TECprobe-VL DMS reactivity matrices for the *B. subtilis gcvT* riboswitch U81G variant in the absence (A) and presence (C) of 1 mM glycine. Transcripts 239-259, which were not enriched, are excluded. (**B**, **D**) *B. subtilis gcvT* glycine aptamer U81G variant secondary structures colored by the TECprobe-VL DMS reactivity of the 223 nt transcript in the absence (B) and presence (D) of 1 mM glycine. Data are from two independent replicates that were processed separately and merged.

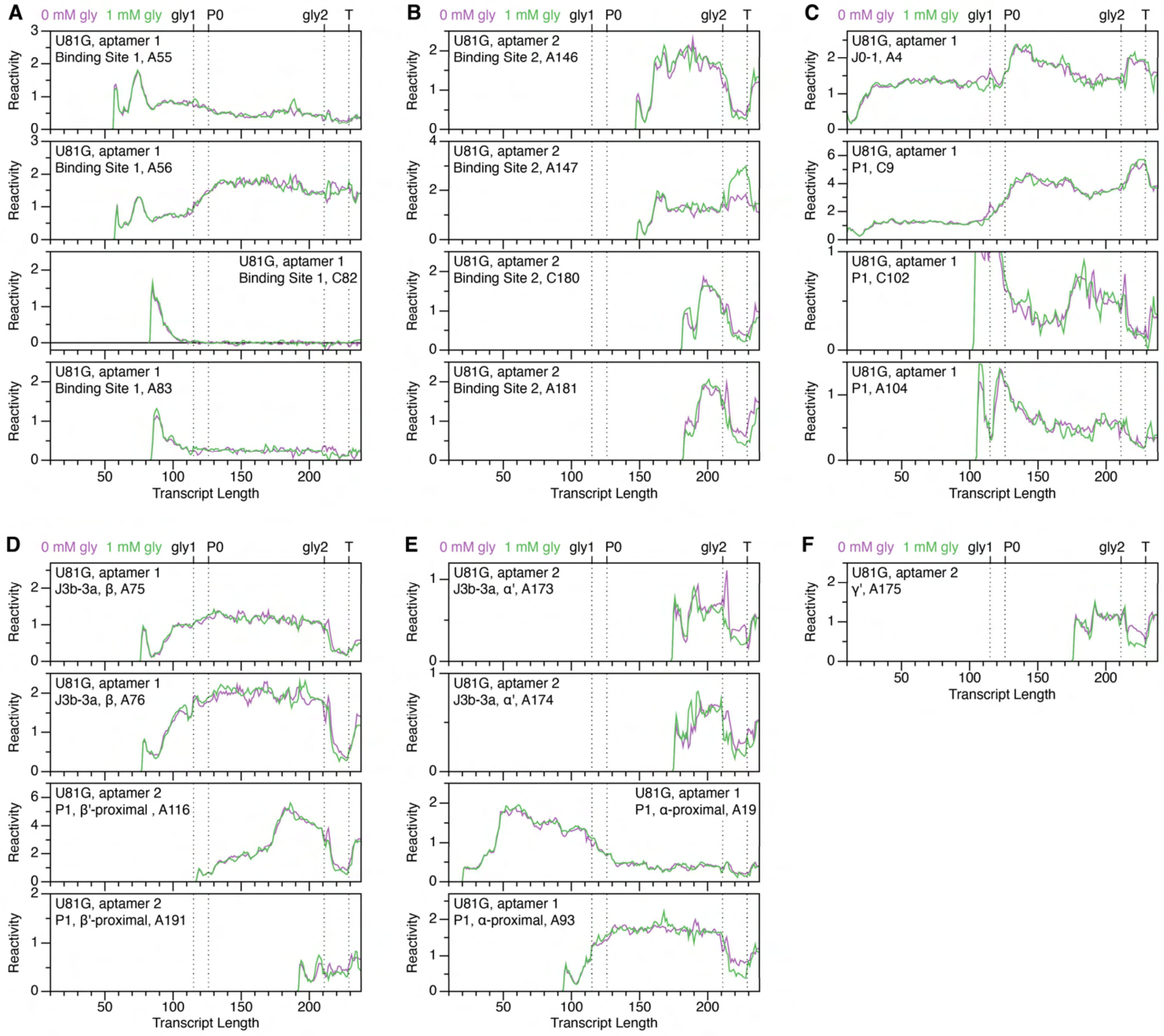

**Supplementary Figure 12. DMS reactivity trajectories for select nucleotides of the *B. subtilis gcvT* riboswitch U81G variant.** (**A-F**) TECprobe-VL DMS reactivity trajectories for nucleotides in or proximal to binding site 1 (A), binding site 2 (B), the kink-turn motif (C),β and β' (D), α and α' (E), and γ' (F). Data are from the experiment shown in Supplementary Figure 11.

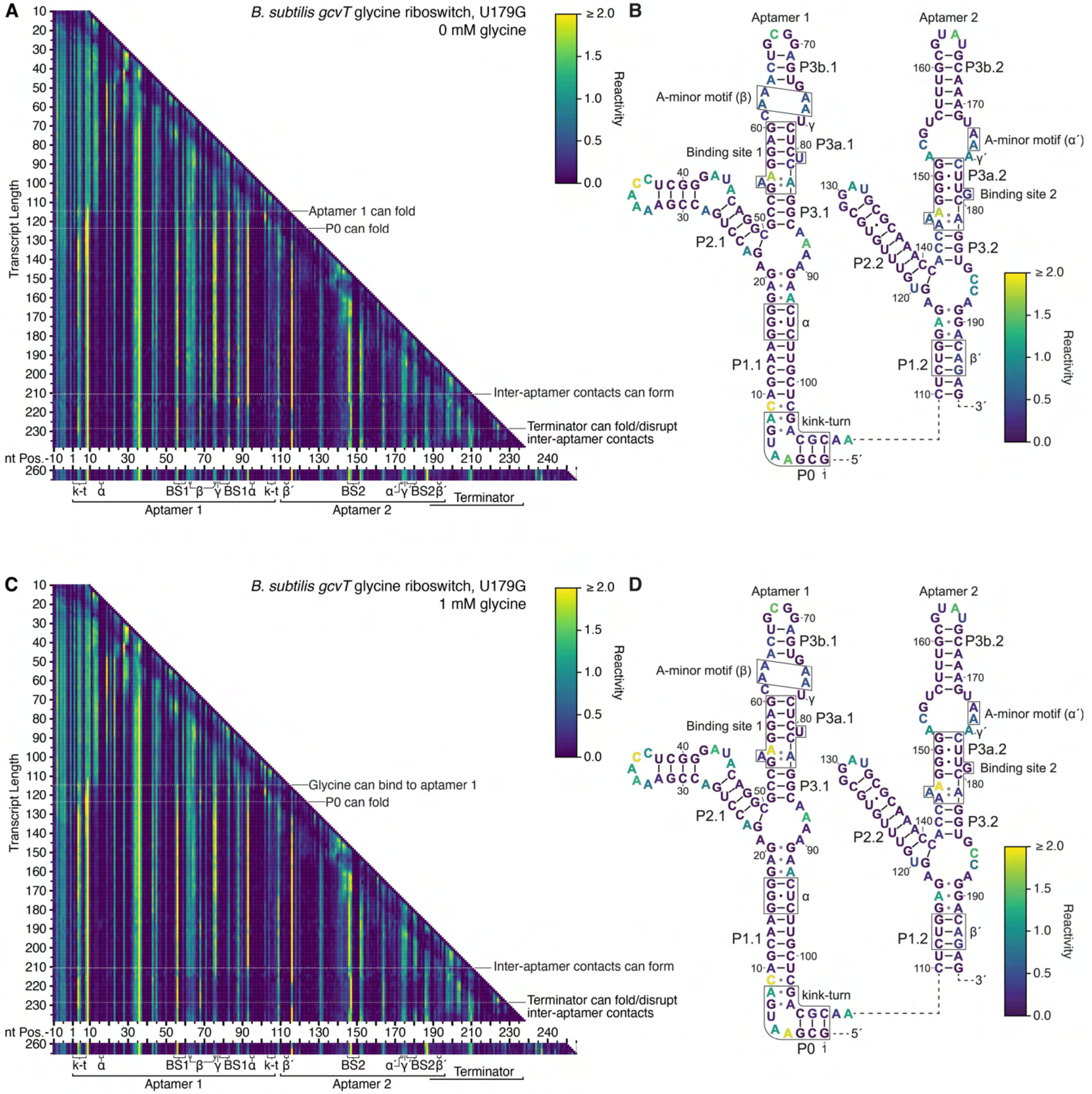

**Supplementary Figure 13. Cotranscriptional DMS structure probing of the *B. subtilis gcvT* riboswitch U179G variant.** (**A**, **C**) TECprobe-VL DMS reactivity matrices for the *B. subtilis gcvT* riboswitch U179G variant in the absence (A) and presence (C) of 1 mM glycine. Transcripts 239-259, which were not enriched, are excluded. (**B**, **D**) *B. subtilis gcvT* glycine aptamer U179G variant secondary structures colored by the TECprobe-VL DMS reactivity of the 223 nt transcript in the absence (B) and presence (D) of 1 mM glycine. Data are from two independent replicates that were processed separately and merged.

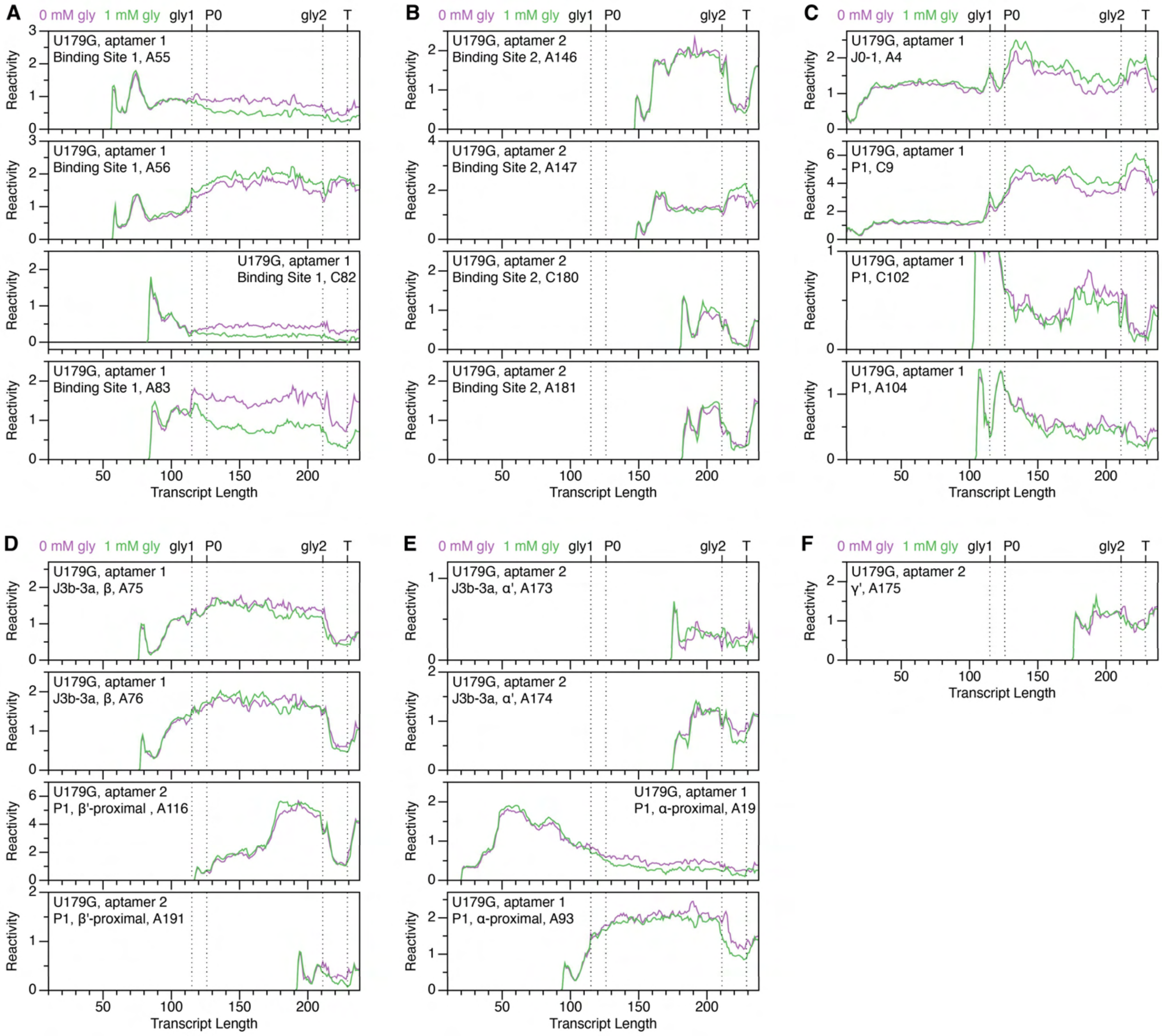

**Supplementary Figure 14. DMS reactivity trajectories for select nucleotides of the *B. subtilis gcvT* riboswitch U179G variant.** (**A-F**) TECprobe-VL DMS reactivity trajectories for nucleotides in or proximal to binding site 1 (A), binding site 2 (B), the kink-turn motif (C), β and β' (D), α and α' (E), and γ' (F). Data are from the experiment shown in Supplementary Figure 13.

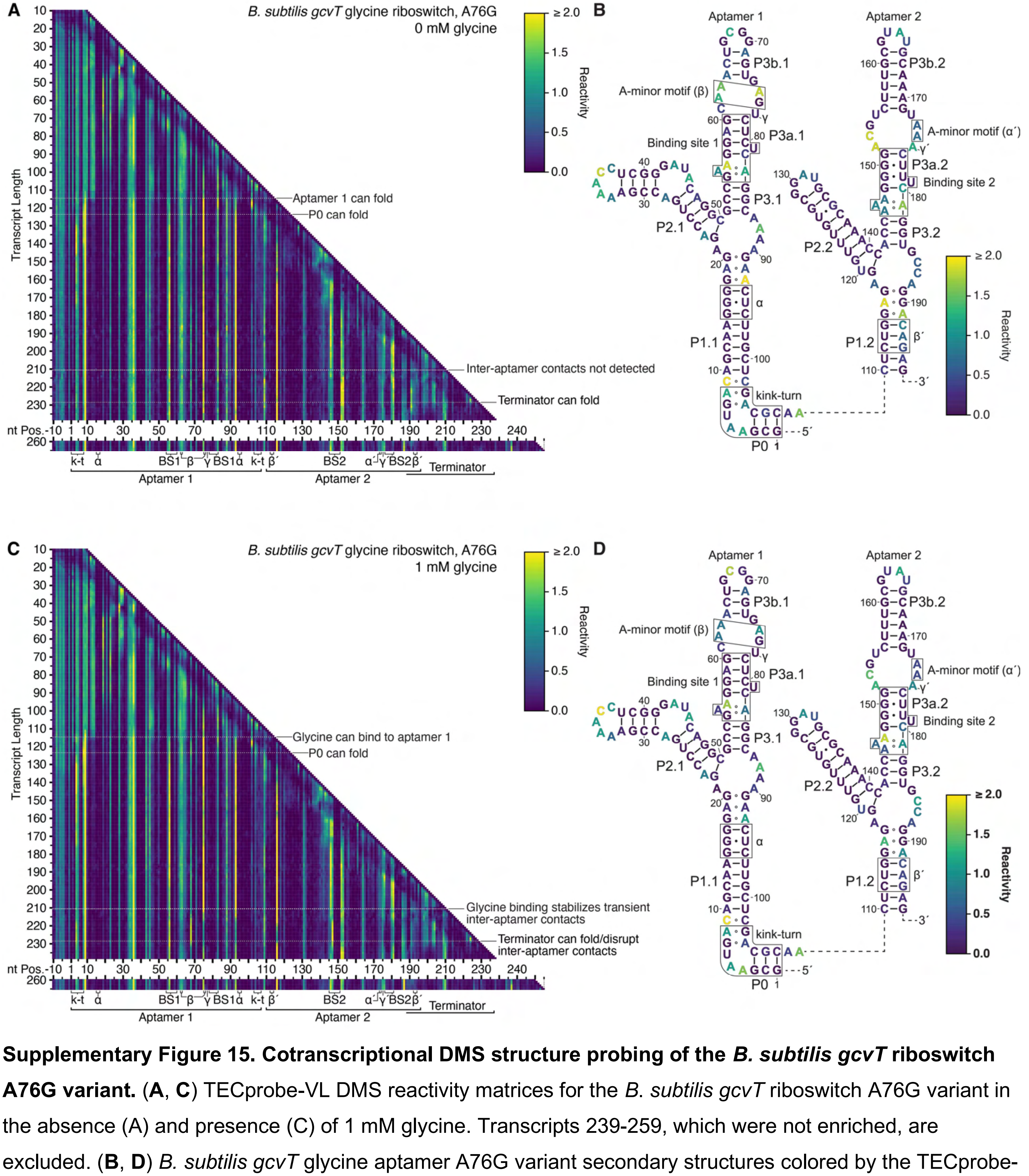

**Supplementary Figure 15. Cotranscriptional DMS structure probing of the *B. subtilis gcvT* riboswitch A76G variant.** (**A**, **C**) TECprobe-VL DMS reactivity matrices for the *B. subtilis gcvT* riboswitch A76G variant in the absence (A) and presence (C) of 1 mM glycine. Transcripts 239-259, which were not enriched, are excluded. (**B**, **D**) *B. subtilis gcvT* glycine aptamer A76G variant secondary structures colored by the TECprobe-VL DMS reactivity of the 223 nt transcript in the absence (B) and presence (D) of 1 mM glycine. Data are from two independent replicates that were processed separately and merged.

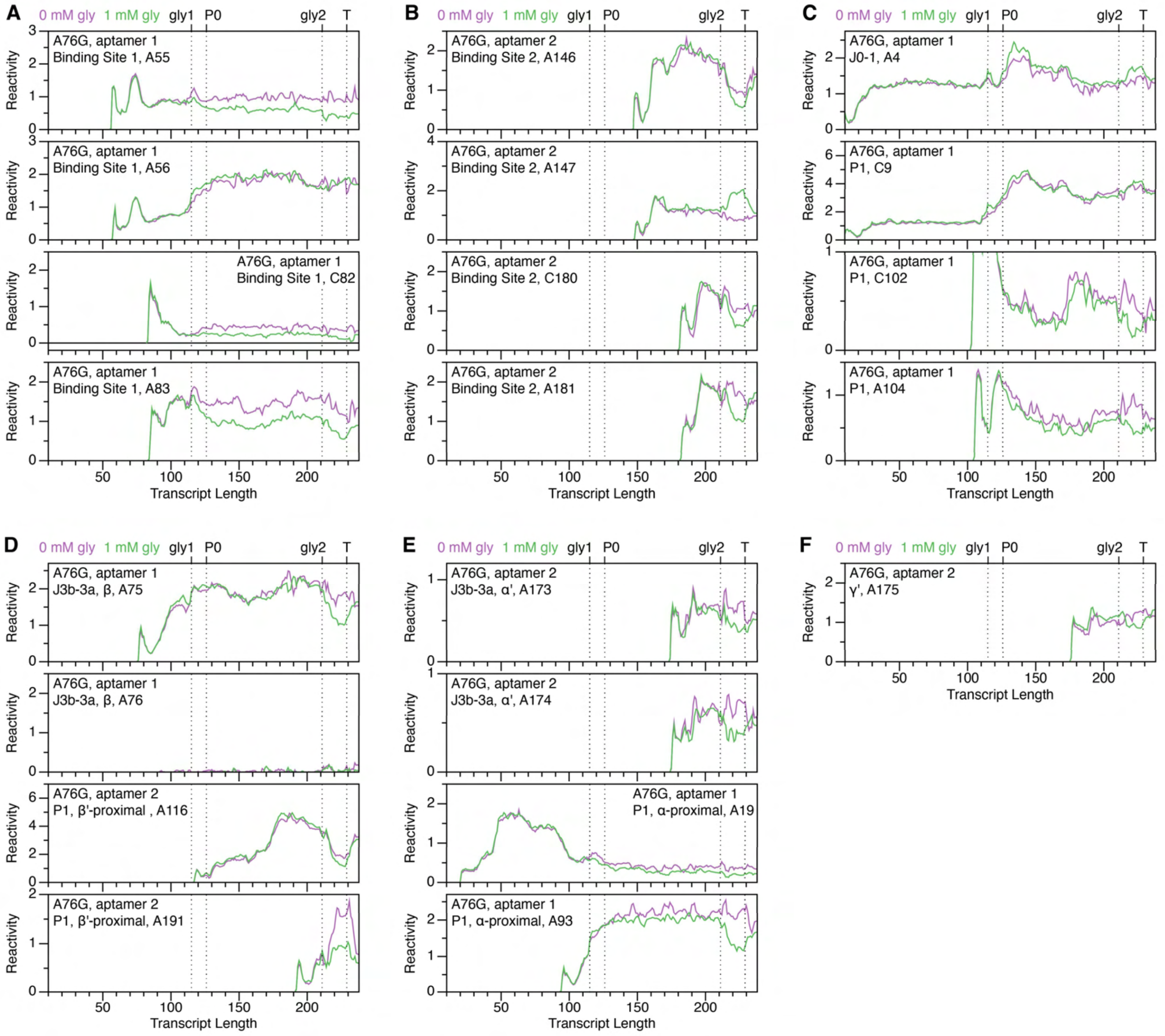

**Supplementary Figure 16. DMS reactivity trajectories for select nucleotides of the *B. subtilis gcvT* riboswitch A76G variant.** (**A**-**F**) TECprobe-VL DMS reactivity trajectories for nucleotides in or proximal to binding site 1 (A), binding site 2 (B), the kink-turn motif (C), β and β' (D), α and α' (E), and γ' (F). Data are from the experiment shown in Supplementary Figure 15.

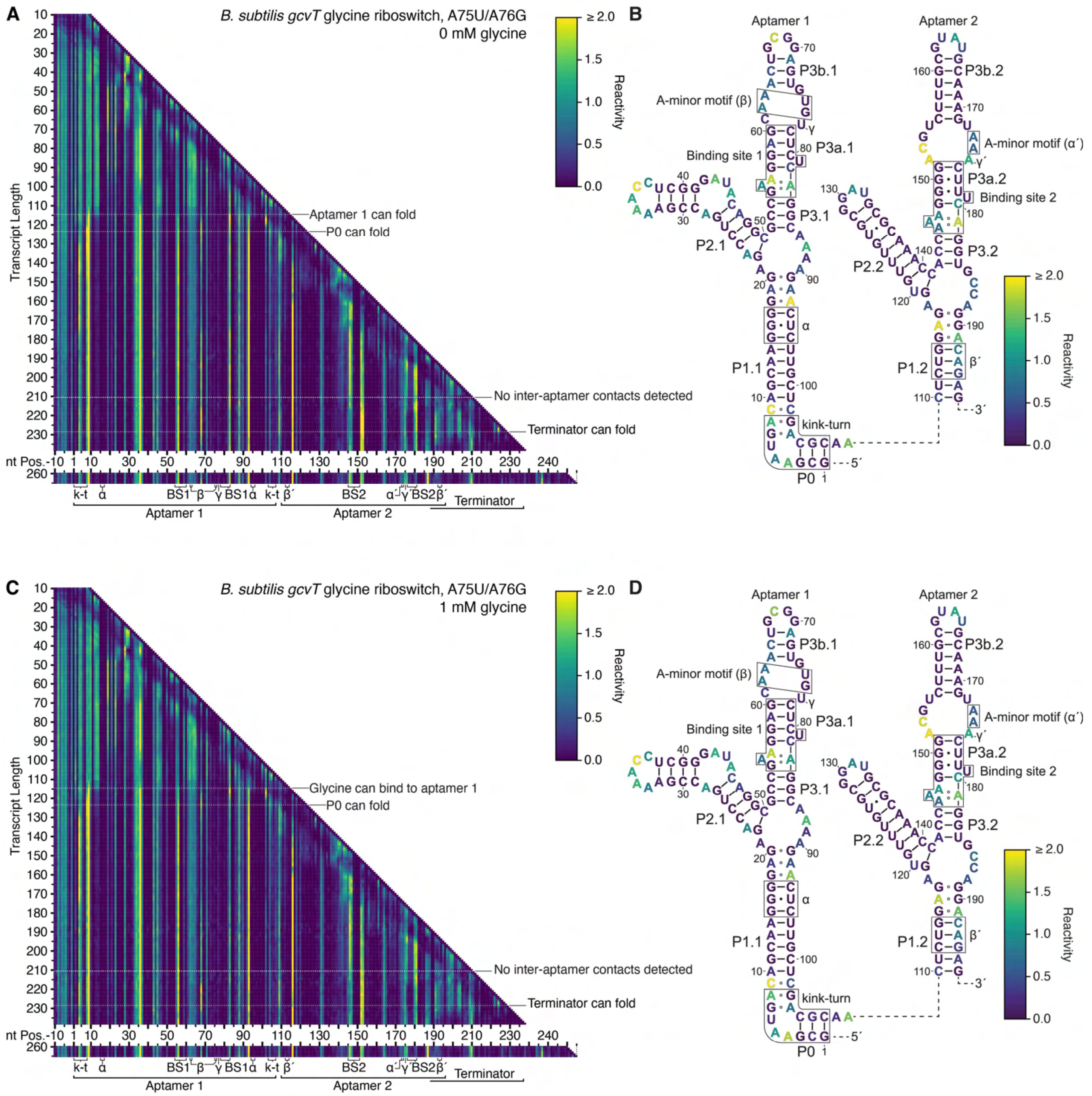

**Supplementary Figure 17. Cotranscriptional DMS structure probing of the *B. subtilis gcvT* riboswitch A75U, A76G variant.** (**A**, **C**) TECprobe-VL DMS reactivity matrices for the *B. subtilis gcvT* riboswitch A75U, A76G variant in the absence (A) and presence (C) of 1 mM glycine. Transcripts 239-259, which were not enriched, are excluded. (**B**, **D**) *B. subtilis gcvT* glycine aptamer A75U, A76G variant secondary structures colored by the TECprobe-VL DMS reactivity of the 223 nt transcript in the absence (B) and presence (D) of 1 mM glycine. Data are from two independent replicates that were processed separately and merged.

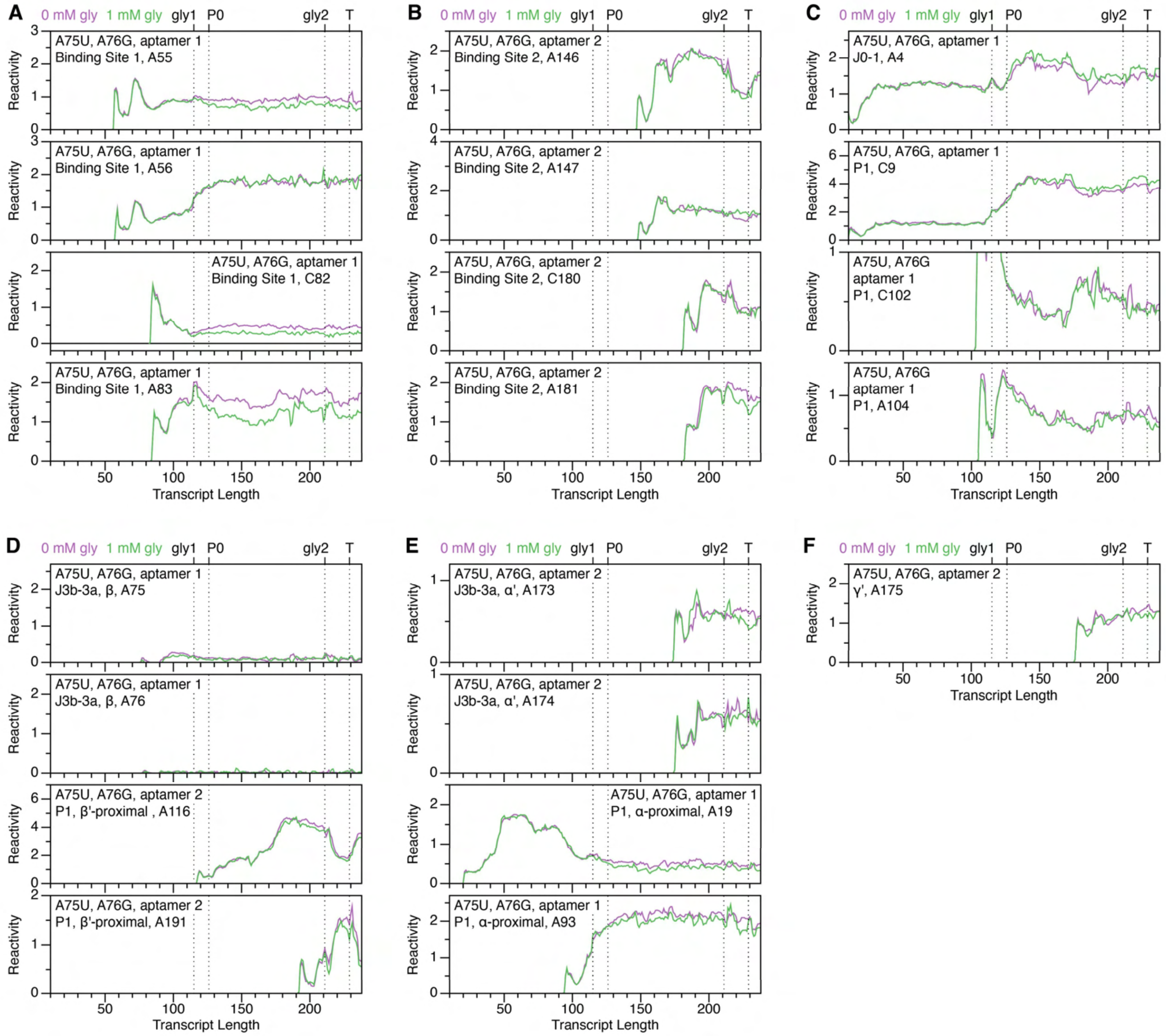

**Supplementary Figure 18. DMS reactivity trajectories for select nucleotides of the *B. subtilis gcvT* riboswitch A75U, A76G variant.** (**A**-**F**) TECprobe-VL DMS reactivity trajectories for nucleotides in or proximal to binding site 1 (A), binding site 2 (B), the kink-turn motif (C), β and β' (D), α and α' (E), and γ' (F). Data are from the experiment shown in Supplementary Figure 17.

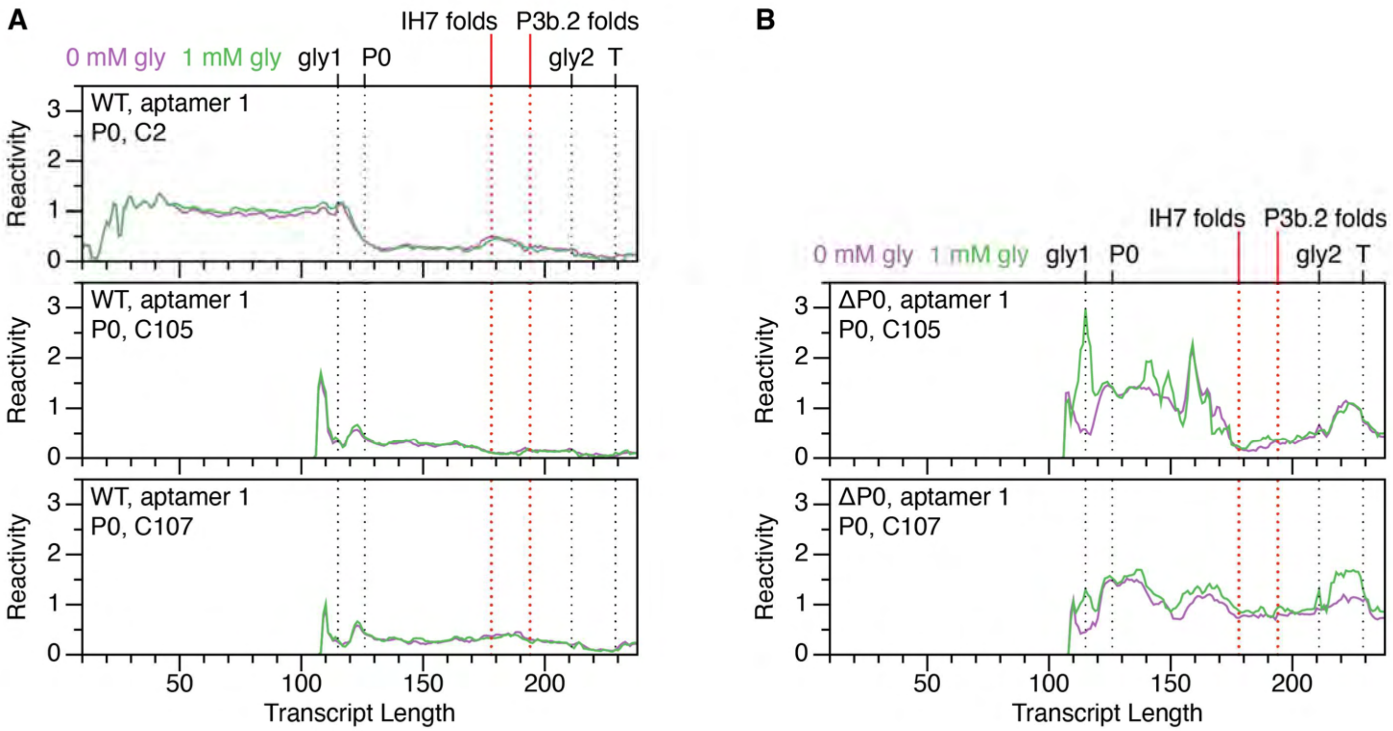

**Supplementary Figure 19. Cotranscriptional folding of P0.** (**A**, **B**) TECprobe-VL DMS reactivity trajectories for P0 nucleotides in the wild-type *B. subtilis gcvT* riboswitch (A) and ΔP0 variant (B). Reactivity data are from Supplementary Figures 7 and 20.

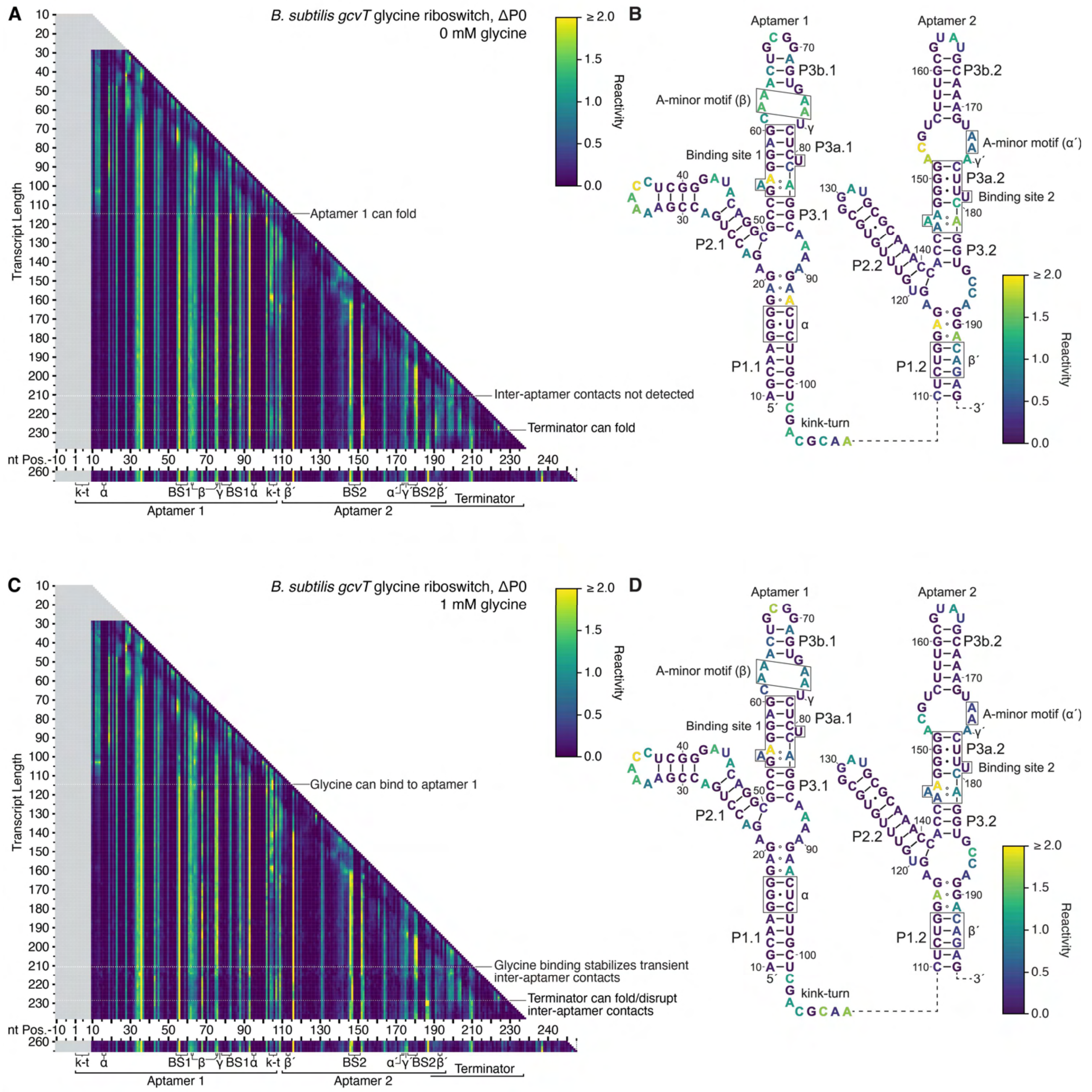

**Supplementary Figure 20. Cotranscriptional DMS structure probing of the *B. subtilis gcvT* riboswitch ΔP0 variant.** (**A**, **C**) TECprobe-VL DMS reactivity matrices for the *B. subtilis gcvT* riboswitch ΔP0 variant in the absence (A) and presence (C) of 1 mM glycine. Transcripts 239-259, which were not enriched, are excluded. (**B**, **D**) *B. subtilis gcvT* glycine aptamer ΔP0 variant secondary structures colored by the TECprobe-VL DMS reactivity of the 223 nt transcript in the absence (B) and presence (D) of 1 mM glycine. Data are from two independent replicates that were processed separately and merged.

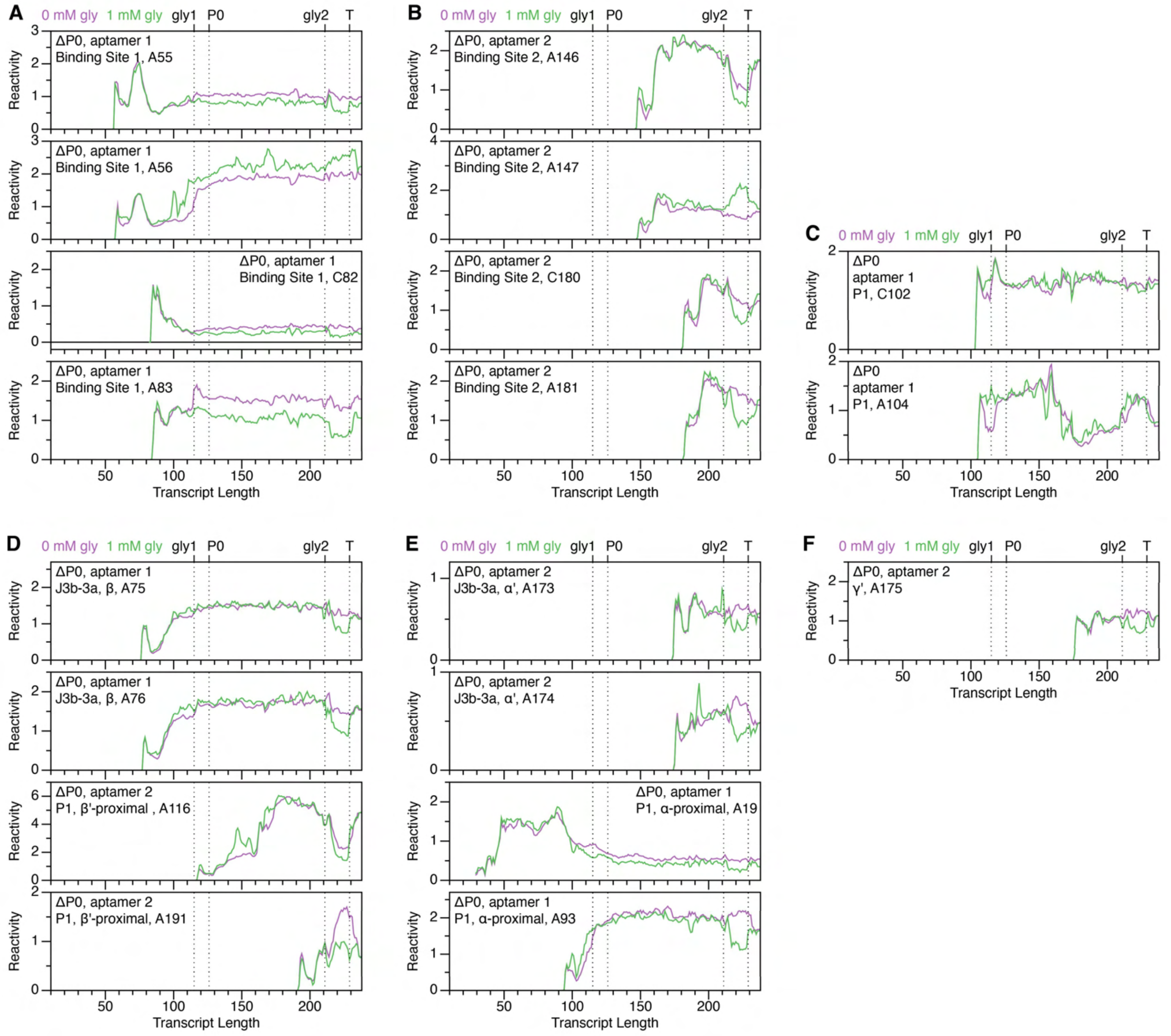

**Supplementary Figure 21. DMS reactivity trajectories for select nucleotides of the *B. subtilis gcvT* riboswitch ΔP0 variant.** (**A-F**) TECprobe-VL DMS reactivity trajectories for nucleotides in or proximal to binding site 1 (A), binding site 2 (B), the kink-turn motif (C), β and β' (D), α and α' (E), and γ' (F). Data are from the experiment shown in Supplementary Figure 20.

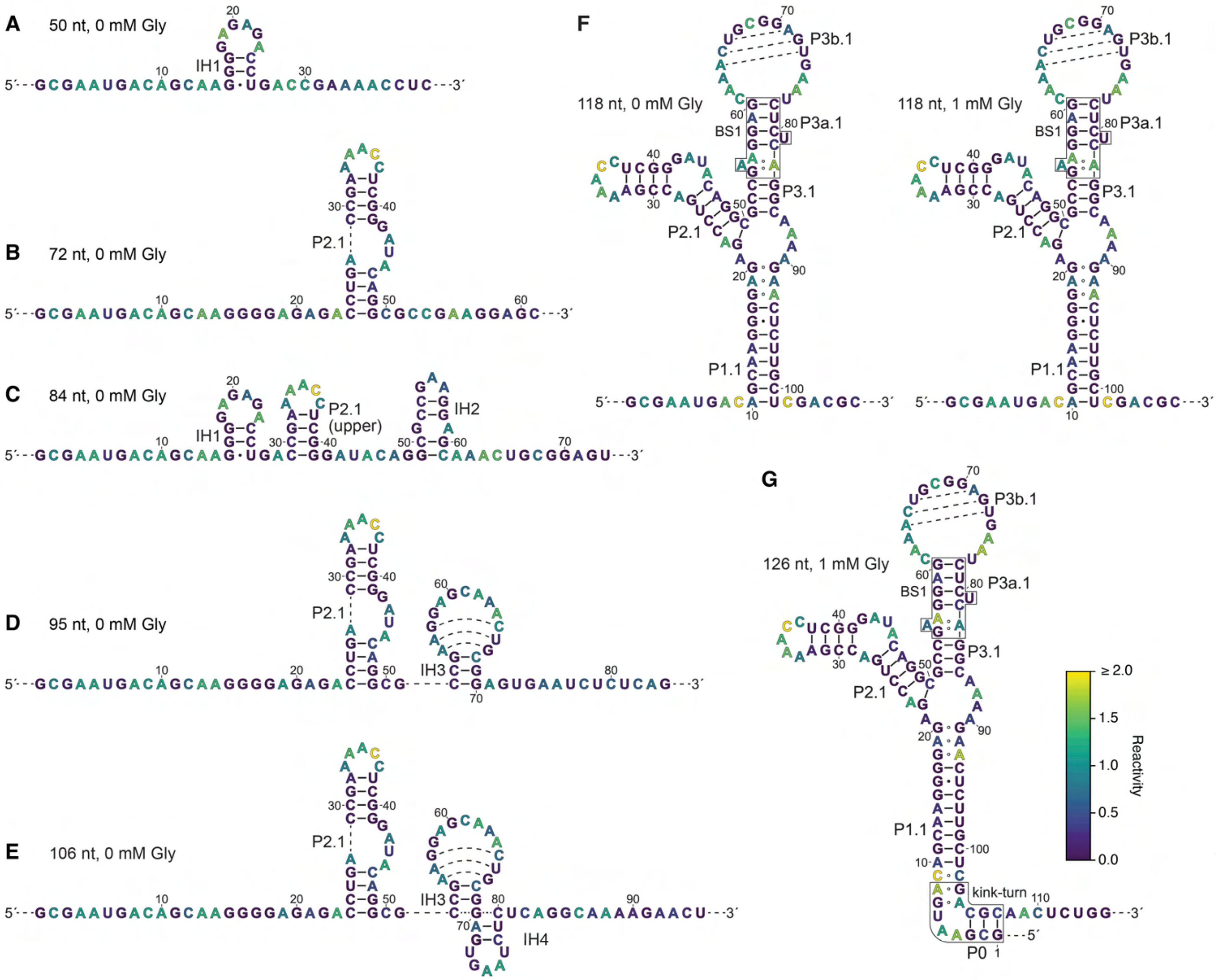

**Supplementary Figure 22. Reactivity-guided structure predictions of aptamer 1 folding intermediates.** (**A**-**G**) *B. subtilis gcvT* glycine aptamer 1 folding intermediate secondary structures colored by TECprobe-VL DMS reactivity. Reactivity data are from Supplementary Figure 7. Secondary structures omit the 3’-most 11 nucleotides of the transcript.

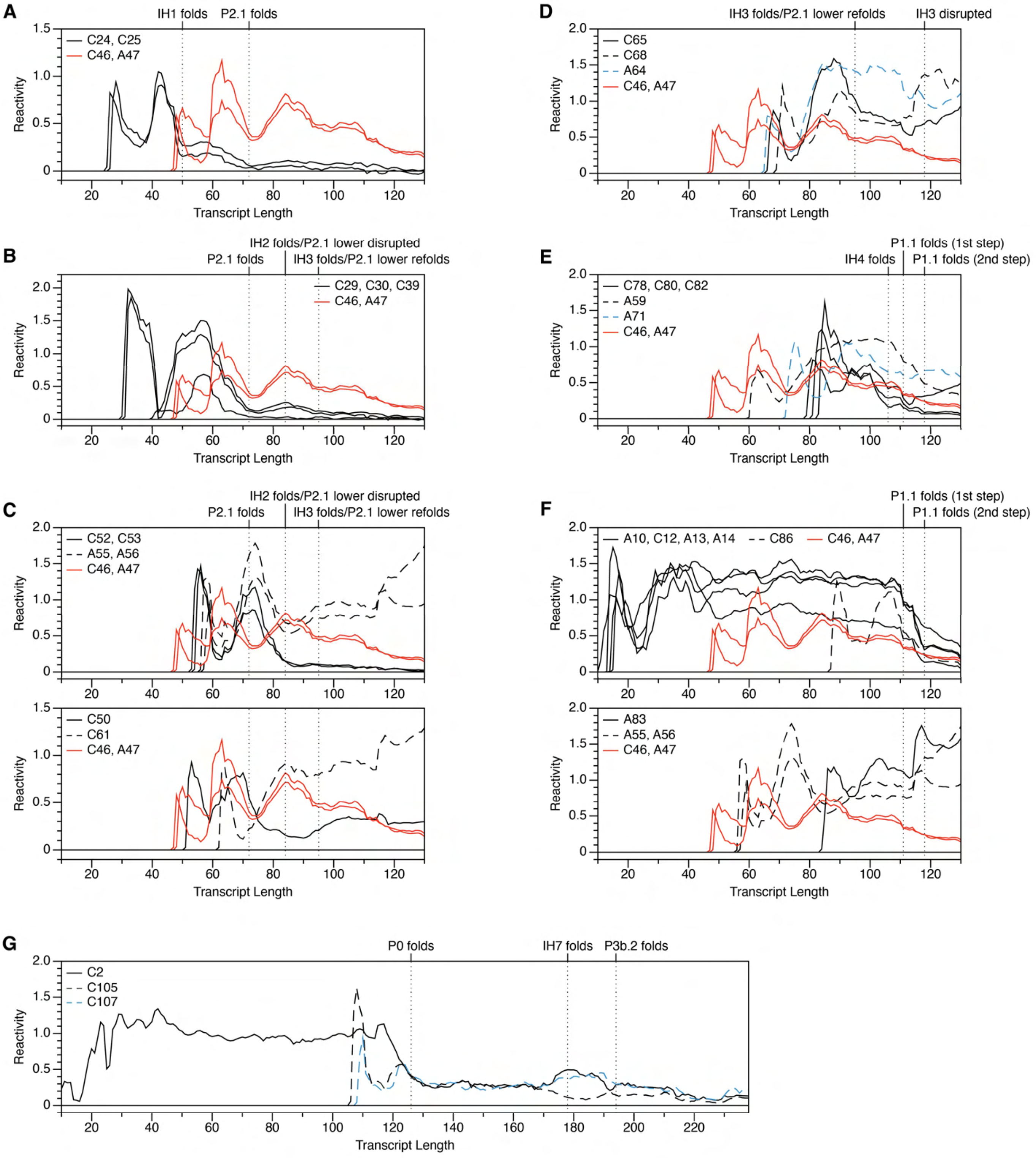

**Supplementary Figure 23. Reactivity trajectories supporting aptamer 1 folding transitions.** (**A-G**) TECprobe-VL DMS reactivity trajectories that support the folding transitions depicted in Supplementary Figure 22. Reactivity data are from Supplementary Figure 7.

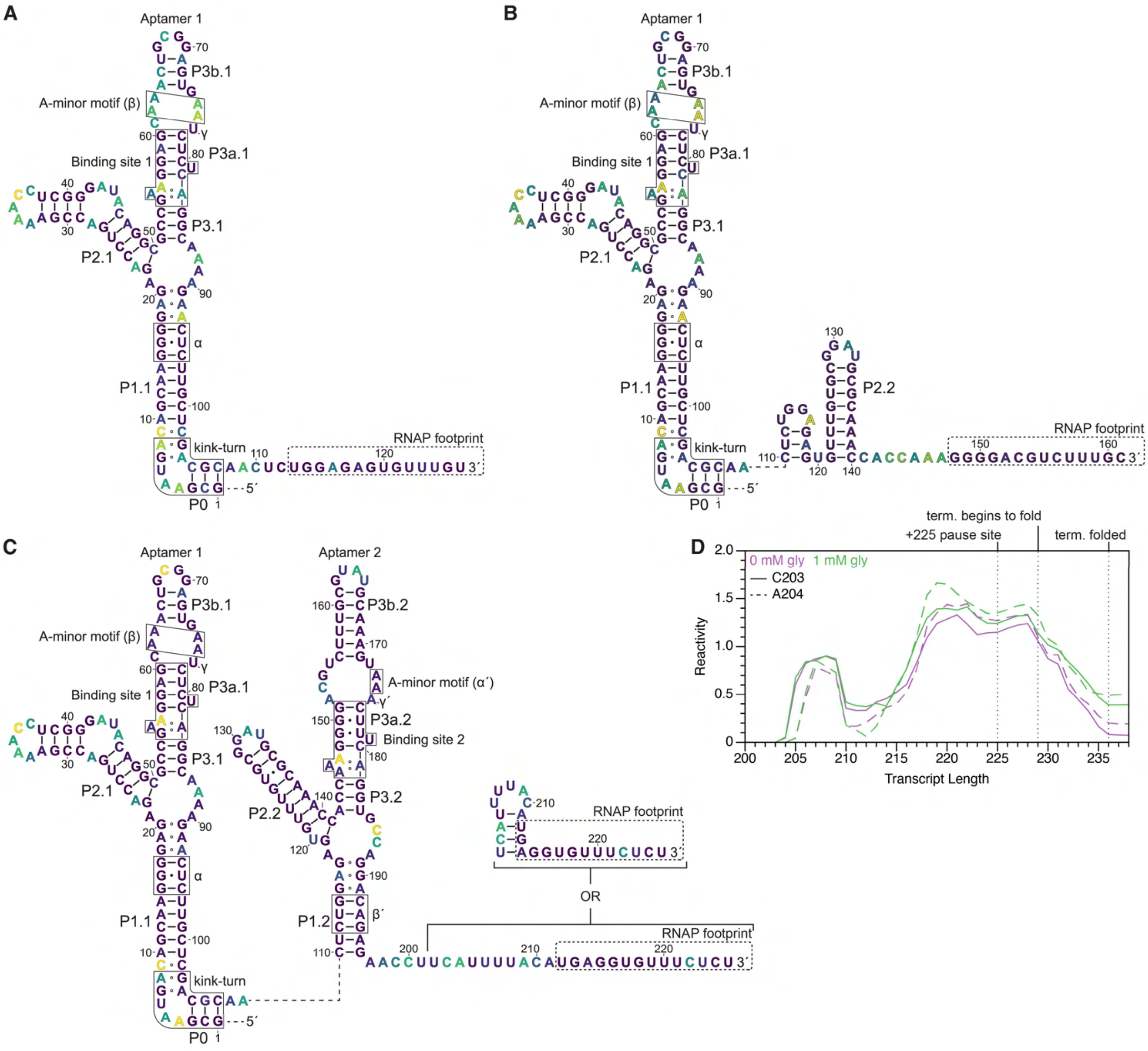

**Supplementary Figure 24. Reactivity-guided structure predictions of paused folding intermediates.** (**A**-**C**) *B. subtilis gcvT* glycine aptamer secondary structures for the +126 (A), +161 (B), and +225 (C) pauses. (**D**) TECprobe-VL DMS reactivity trajectories for C203 and A204. Reactivity data are from Supplementary Figure 7.

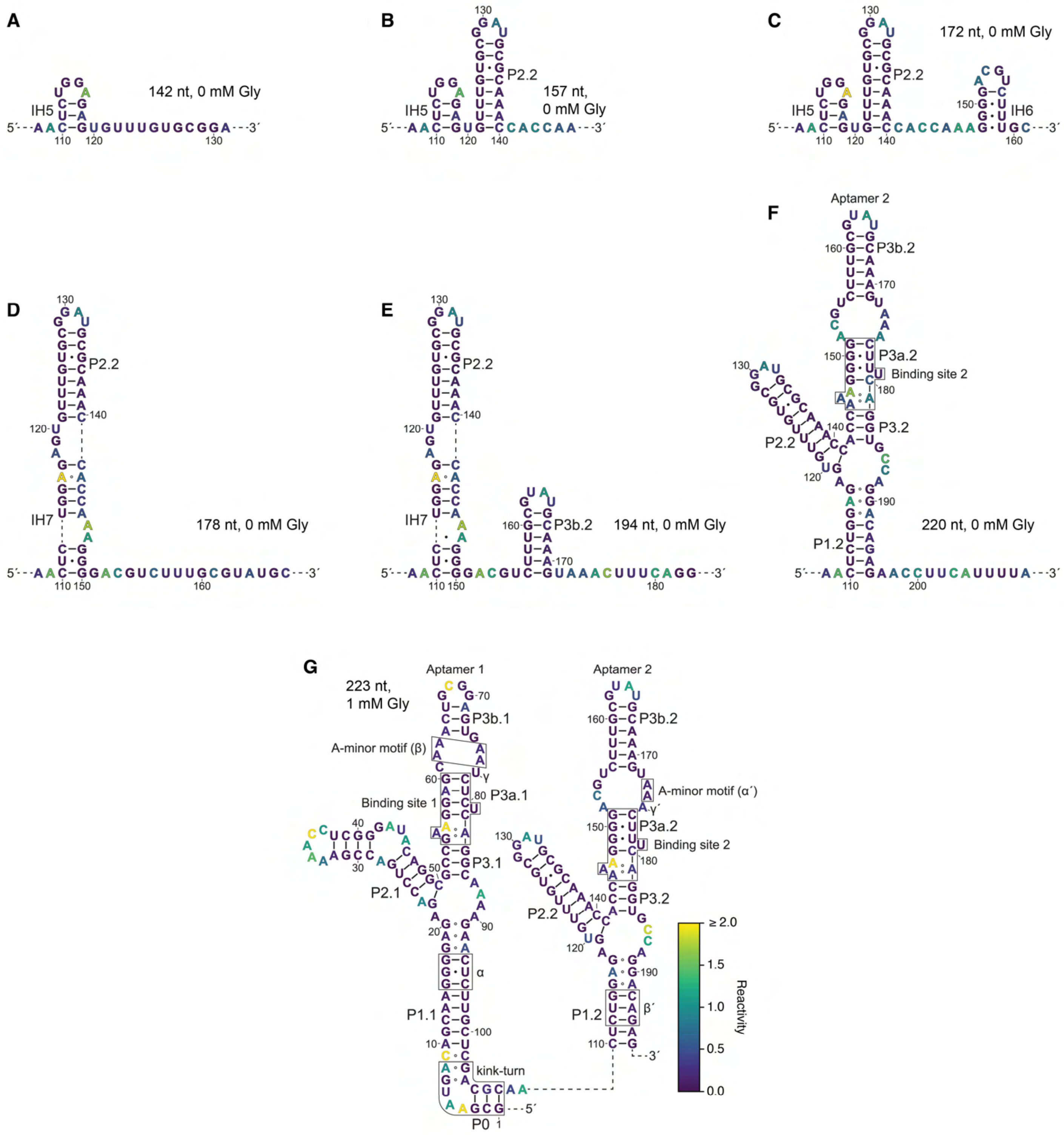

**Supplementary Figure 25. Reactivity-guided structure predictions of aptamer 2 folding intermediates.** (**A**-**G**) *B. subtilis gcvT* glycine aptamer 2 folding intermediate secondary structures colored by TECprobe-VL DMS reactivity. Reactivity data are from Supplementary Figure 7. Secondary structures omit the 3’-most 11 nucleotides of the transcript.

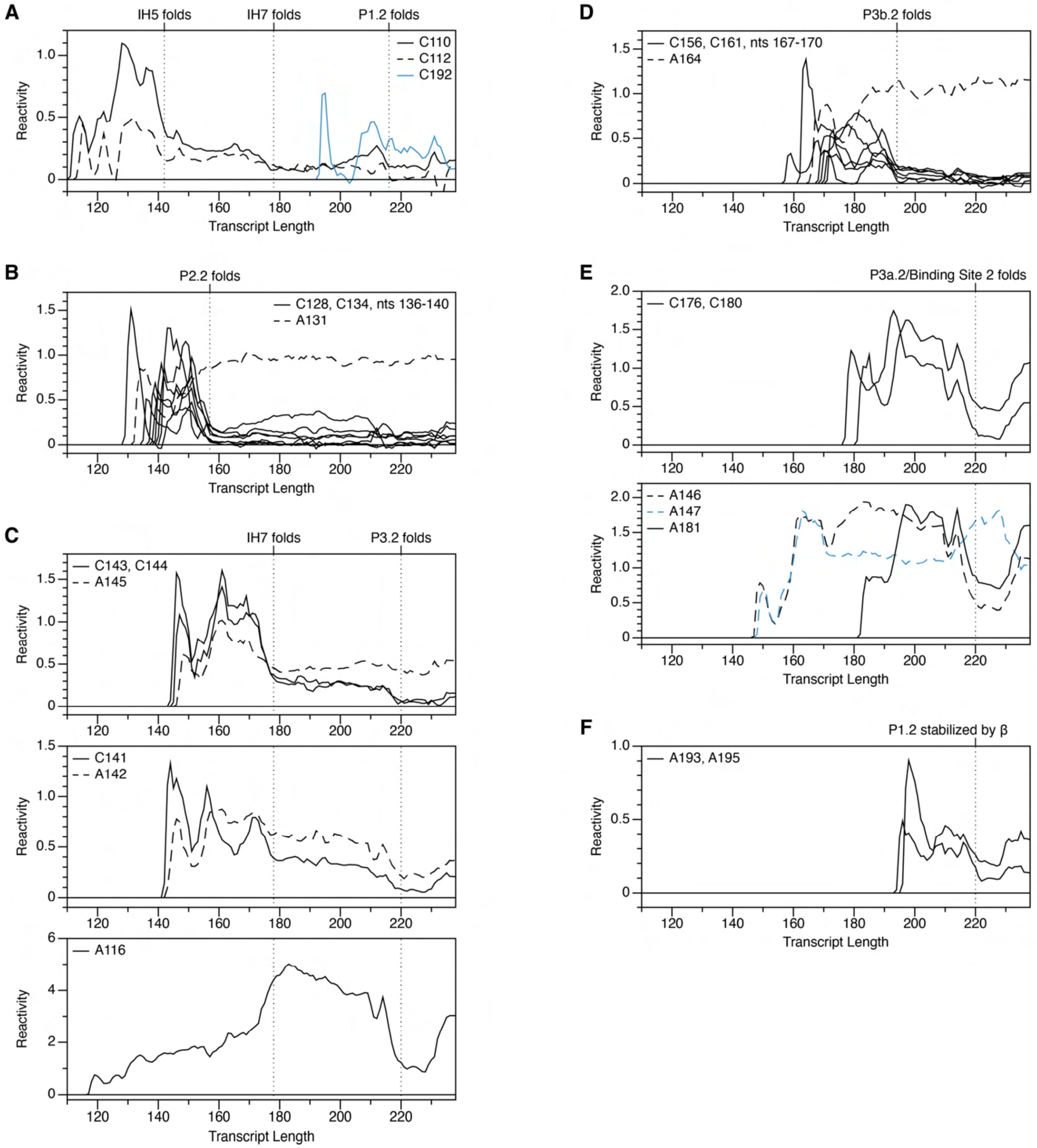

**Supplementary Figure 26. Reactivity trajectories supporting aptamer 2 folding transitions.**

(**A-F**) TECprobe-VL DMS reactivity trajectories that support the folding transitions depicted in Supplementary Figure 25. Reactivity data are from Supplementary Figure 7.

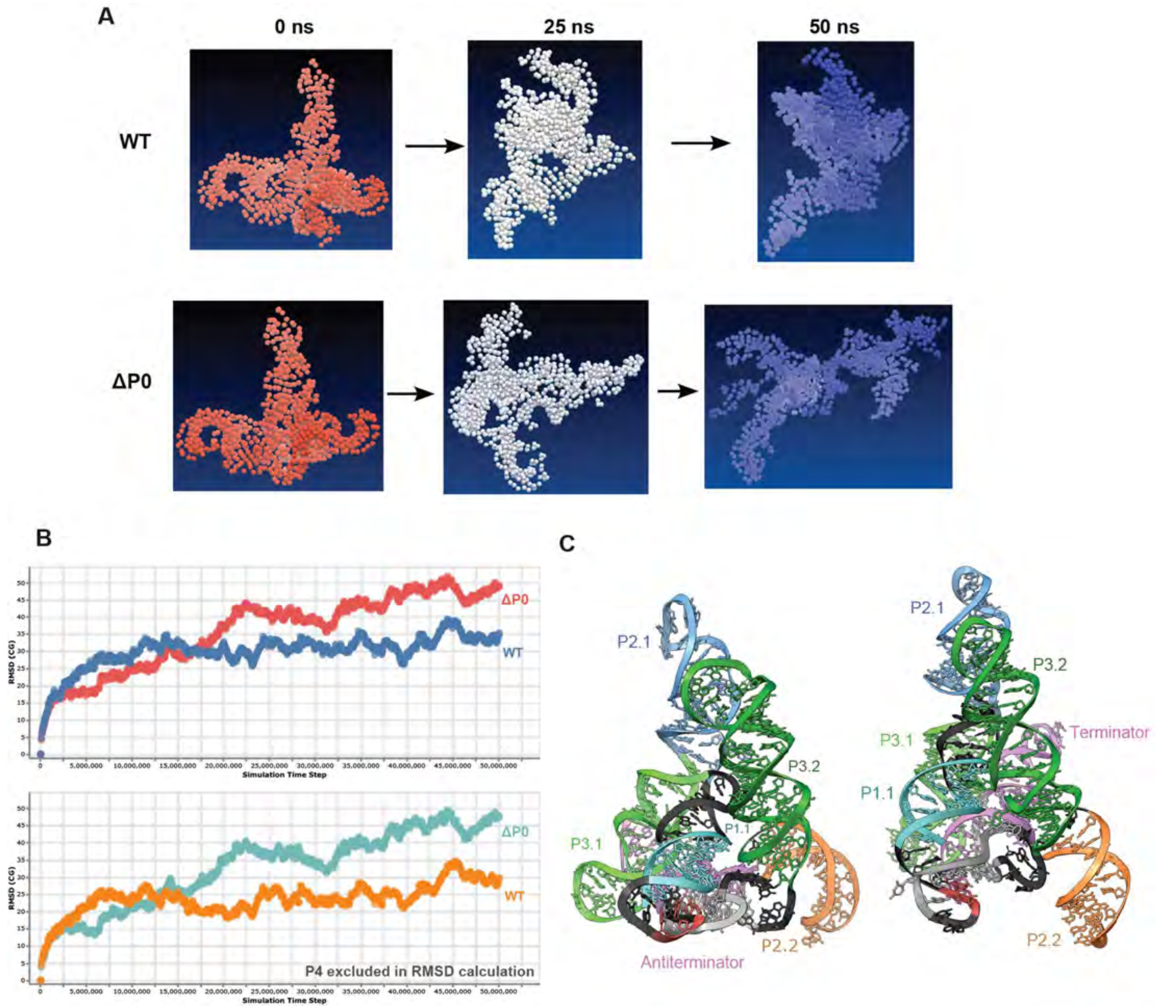

**Supplementary Figure 27. Computational simulations probing folding stability of the ΔP0 mutant.** Simulation reveals that the elimination of P0 reduces folding stability. (**A**) The coarse-grained structures obtained from the simulation trajectory for the WT and the ΔP0 variant. The ΔP0 variant becomes disrupted after 25 ns and fully extends by the end of the 50 ns simulation. In contrast, the WT structure remains relatively compact throughout the simulation. (**B**) The RMSD of the coarse-grained structures during the simulation against the initial structure. In the top panel, the RMSD values are computed using the complete structures, while in the bottom panel, the RMSD values are calculated using the structure excluding P4. The ΔP0 variant experienced a larger structural change throughout the simulation, even excluding the flexible P4 domain. (**C**) Two structures modeled for the ΔP0 variant as an antiterminator (left) and terminator (right), respectively. The structures are highlighted to show the nucleotide regions in Figure 6a and 6b. The antiterminator and terminator structures for the ΔP0 variant exhibit significant structural differences when compared to the WT structures (Figures 6a and 6b), where the domain topologies are remarkably different.

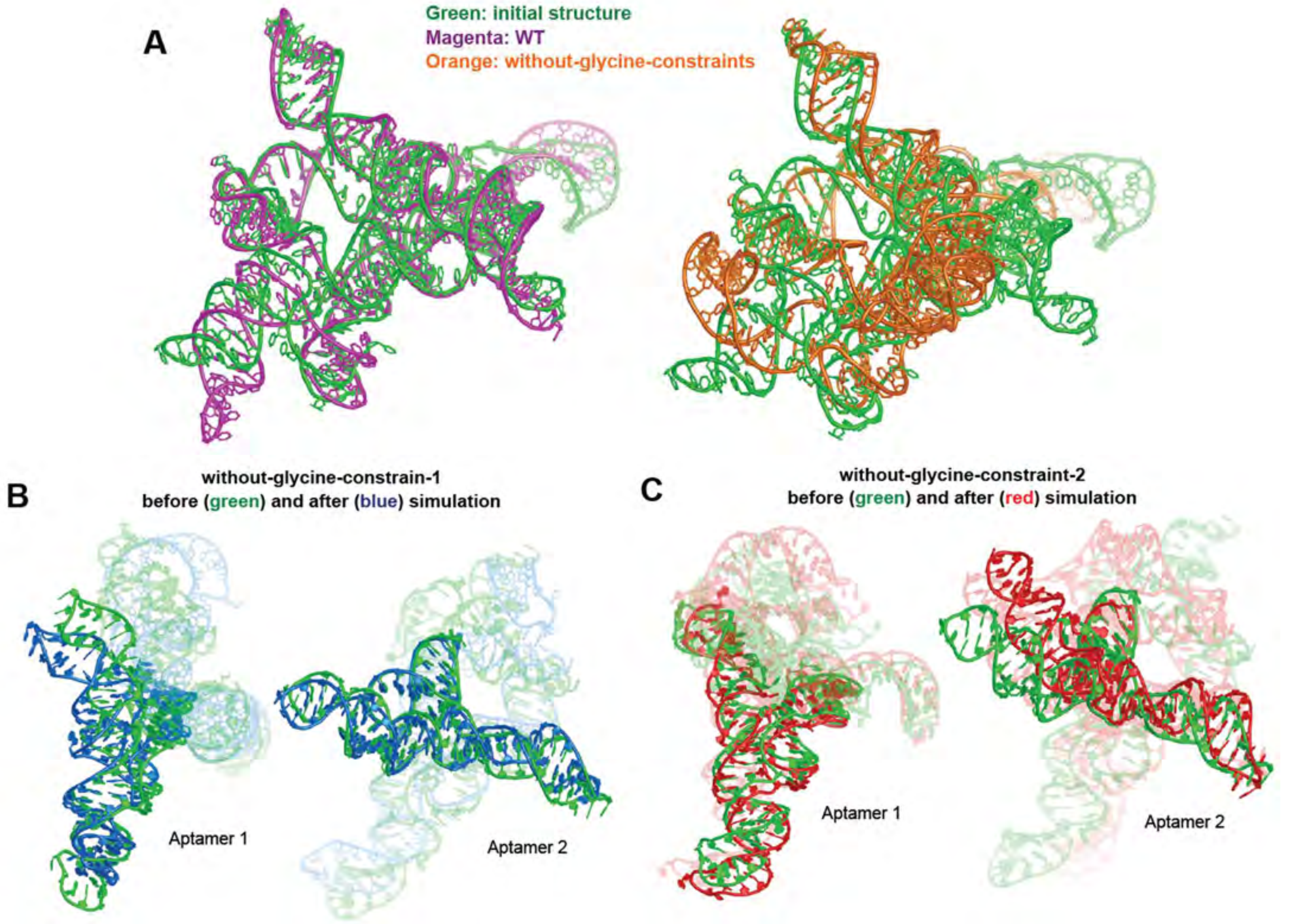

**Supplementary Figure 28**. **Glycine binding stabilizes individual aptamers.**

Simulation reveals that the disruption of glycine binding pockets results in decreased stability. (**A**) Simulation results for the WT and the "without-glycine-constraints" model. Starting from the same initial structure (green), we obtained the top-ranked models for the WT (magenta) and without-glycine-constraints model (orange). The two models are aligned with the initial structure, demonstrating that compared to the WT, the without-glycine-constraints model involves significant structural changes along the simulation. (**B**) Simulation results for the "without-glycine-constraint-1" model. The top-ranked model obtained from the simulation is aligned with the initial structure (green) using aptamer 1 (left) and aptamer 2 (right), indicating that the without-glycine-constraint-1 model causes disruption in the aptamer 1 region, rather than in the aptamer 2 region. (**C**) Simulation results for the without-glycine-constraint-2 model. The top-ranked model obtained from the simulation is aligned with the initial structure (green) using aptamer 1 (left) and aptamer 2 (right), indicating that the without-glycine-constraint-2 model causes disruption in the aptamer 2 region, rather than in the aptamer 1 region.

**Table S1: Oligonucleotides used in single-molecule experiments.** The table below shows oligonucleotides used to perform single-molecule experiments in this study. The modification codes are derived from the Integrated DNA Technologies ordering form, in particular:

/5Biosg/: 5' Biotin

/5deSBioTEG/: 5' Desthiobiotin-TEG

/5Cy5/: 5’ Cy5™

| Oligonucleotide | Sequence (5’-3’) | Purification |
|---|---|---|
| Bsu_GTR-Prom-Fwd (1) | TCCAGATCCCGAAAATTTATCAAAAAGAGTATTGACTTAAAGTCTAACCTATAGGATACTTACAGCCATGAGCGAATGACAGCAAGGGGA | None |
| Bsu_GTR-Prom–Amp-REV(1) | GAGCAAGAGTTCTTTTGCCTGAGAGATTCACTCCGCAGTTTGCTCCTTCGGCGCCTGTATCCCGAGGTTTTCGGTCAGGTCTCTCCCCTTGCTGTCATTCGCT | None |
| Bsu_GTR-Amp-Fwd (2) | TCTCAGGCAAAAGAACTCTTGCTCGACGCAACTCTGGAGAGTGTTTGTGCGGATGCGCAAACCACCAAAGGGGACGTCTTTGCGTATGCA | None |
| Bsu_GTR-Amp-ExpP-Rev(2) | CCCTTTATCAACGGCGCAGCTAAAAAACATACAAAAAAGGACAGAGAAACACCTCATGTAAAATGAAGGTTCTCTGTCCTGGCACCTGAAAGTTTACTTTGCATACGCAAAGACGTCCC | None |
| Bsu_GTR-T7A1-PCR | TCCAGATCCCGAAAATTTATCAAAAAGAGTATTG | None |
| Bsu_GTR-AN-Rev | AGACCACGTTGAAAGATTGGGTTACGGACAGAGAAACACCTCATGTAAAATGAAG | None |
| Anchor_bio | /5Biosg/AGACCACGTTGAAAGATTGGGTTAC | HPLC |
| Bsu_GTR-PEC225-Desthio | /5deSBioTEG/AAAAAGGACAGAGAAACACCTCATGTAAAATG | HPLC |
| Bsu-GTR-ΔP0 (2) | CTCCCCTTGCTATGGCTGTAAGTATC | None |
| Bsu-GTR-ΔP0 (3) | GATACTTACAGCCATAGCAAGGGGAG | None |
| Bsu_GTR-A76G (2) | TTGCCTGAGAGACTCACTCCGCAGT | None |
| Bsu_GTR-A76G (3) | ACTGCGGAGTGAGTCTCTCAGGCAA | None |
| Bsu_GTR-Mut-BS2-U183G-Rev | GTCCTGGCACCTGCAAGTTTACTTTG | None |
| Bsu_GTR-Mut-BS1 U85G-Fwd | GGAGTGAATCGCTCAGGCAAAAGAA | None |
| Bsu_GTR-Mut-BS1 U85G-Rev | TTCTTTTGCCTGAGCGATTCACTCC | None |
| Bsu_GTR-Mut-BS2-U183G-Fwd | CAAAGTAAACTTGCAGGTGCCAGGAC | None |
| Bsu_GTR-RT-196 Rev | AGACCACGTTGAAAGATTGGGTTACCTCTGTCCTGGCACCTGAAAGT | HPLC |
| Bsu_GTR-PEC225 Rev | /5Biosg/AAAAAGGACAGAGAAACACCTCATGTAAAAT | HPLC |
| Bsu_GTR-PEC161 Rev | /5Biosg/TTTGCATACGCAAAGACGTCCC | HPLC |
| Bsu_GTR-PEC 126 Rev | /5Biosg/CGCATCCGCACAAACACTCTC | HPLC |
| SKM-BsuTerm | /5Cy5/GAA ACA CC | HPLC |
| SKP-Apt1-P1 | /5Cy5/AAG AGT TC | HPLC |
| SKP-Apt2-P2b | /5Cy5/CAA ACA CT | HPLC |

**Table S2: Termination parameters determined in this study.** This table summarizes all the *in vitro* transcription assays, with standard deviations of the fits.

| Template | Amplitude (%)[a] | $R_{50}$ (μM)[b] |
|---|---|---|
| WT | 43.4 ± 1.2 | 5.2 ± 0.7 |
| WT + NusA | 13.5 ± 0.7 | 2.5 ± 0.5 |
| WT + YbxF | 14.4 ± 1.3 | 1.6 ± 0.2 |
| A76G | 10.2 ± 1.1 | 6.2 ± 2.3 |
| ∆P0 | 30.7 ± 1.2 | 70.7 ± 11.3 |
| ∆P0 + YbxF | 26.4 ± 3.0 | 58.3 ± 15.7 |
| U81G | 14.5 ± 0.3 | 28.6 ± 3.8 |
| U179G | 8.4 ± 1.0 | 7.7 ± 4 |

[a]The reported error is the standard deviation (SD) of the mean from three independent replicates.

[b]The reported error is the standard deviation of the fits.

**Table S3: Pause half-lives measured in this study.** This table summarizes the pause half-lives of all *Bsu*-GTR pause sites in the presence and absence of NusA, with standard deviations of the fits.

| **Pause species** | **Glycine** | **τ (s) [pause half-life]*** |
|---|---|---|
| U126 | - | 28 ± 7 |
| | + | 31 ± 14 |
| C161 | - | 65 ± 18 |
| | + | 62 ± 12 |
| U225 | - | 160 ± 44 |
| | + | 121 ± 5 |
| U126 + NusA | - | 39 ± 6 |
| | + | 50 ± 1 |
| C161 + NusA | - | 125 ± 14 |
| | + | 213 ± 23 |
| U225 + NusA | - | 101 ± 43 |
| | + | 1052 ± 194 |

*The reported error is the standard deviation (SD) of the mean from three independent replicates.

**Table S4: Kinetic parameters extracted from the NusA co-localization assay.** Table summarizing the NusA binding kinetics for PEC-161 and PEC-225, with standard deviations of the fits.

| Construct | Glycine | $k_{on}$ ($10^6$ $M^{-1}$ $s^{-1}$) | $k_{off}$ ($s^{-1}$) |
|---|---|---|---|
| PEC-225 | - | Fast[a]: 5.40 ± 0.02 (62%)<br>Slow[a]: 1.35 ± 0.008 (38%)<br>Overall[b]: 4.55 ± 1.33 | Fast[a]: 0.76 ± 0.02 (84%)<br>Slow[a]: 0.12 ± 0.01 (16%)<br>Overall[b]: 0.74 ± 0.16 |
| PEC-225 | + | Fast[a]: 7.26 ± 0.04 (50%)<br>Slow[a]: 1.38 ± 0.01 (50%)<br>Overall[b]: 4.16 ± 0.98 | Fast[a]: 0.74 ± 0.02 (75%)<br>Slow[a]: 0.08 ± 0.01 (25%)<br>Overall[b]: 0.47 ± 0.11 |
| PEC-161 | - | Fast[a]: 13.7 ± 0.1 (49%)<br>Slow[a]: 2.18 ± 0.01 (51%)<br>Overall[b]: 7.51 ± 0.47 | Fast[a]: 1.11 ± 0.05 (70%)<br>Slow[a]: 0.25 ± 0.02 (30%)<br>Overall[b]: 0.87 ± 0.06 |
| PEC-161 | + | Fast[a]: 23.4 ± 0.14 (45%)<br>Slow[a]: 2.29 ± 0.01 (55%)<br>Overall[b]: 11.2 ± 2.31 | Fast[a]: 1.10 ± 0.04 (75%)<br>Slow[a]: 0.23 ± 0.02 (25%)<br>Overall[b]: 0.81 ± 0.12 |

[a]Values were calculated from single- or double-exponential fits of the pooled data from all the experiments under a given condition. The percentages indicate the contributions of each phase to the overall rate constant. The reported error is the standard deviation (SD) of the fit.

[b]Values represent the average ± the standard deviation (SD) of the mean from three independent replicates.

**Table S5: Kinetic parameters extracted from SiM-KARTS analysis.** Table summarizing all SiM-KARTS probe binding kinetics for different RNAs and PECs, with standard deviations of the fits.

| Construct | Glycine | $k_{on}$ ($10^6$ $M^{-1}$ $s^{-1}$) | | $k_{off}$ ($s^{-1}$) |
|---|---|---|---|---|
| PEC-225 | - | Fast[a]: 23.7 ± 0.31 (20%)<br>Slow[a]: 6.42 ± 0.01 (80%)<br>Overall[b]: 9.95 ± 1.62 | | Fast[a]: 0.88 ± 0.02 (91%)<br>Slow[a]: 0.12 ± 0.02 (9%)<br>Overall[b]: 0.82 ± 0.10 |
| | + | Fast[a]: 39.7 ± 0.10 (16%)<br>Slow[a]: 2.41 ± 0.01 (84%)<br>Overall[b]: 4.84 ± 1.41 | | Fast[a]: 0.76 ± 0.02 (82%)<br>Slow[a]: 0.16 ± 0.01 (18%)<br>Overall[b]: 0.74 ± 0.24 |
| PEC-161 | - | Fast[a]: 3.10 ± 0.01 (100%)<br>Slow[a]: NA<br>Overall[b]: 2.89 ± 0.75 | | Fast[a]: 0.89 ± 0.01 (100%)<br>Slow[a]: NA<br>Overall[b]: 0.95 ± 0.18 |
| | + | Fast[a]: 2.45 ± 0.01 (100%)<br>Slow[a]: NA<br>Overall[b]: 2.37 ± 0.51 | | Fast[a]: 0.92 ± 0.02 (92%)<br>Slow[a]: 0.10 ± 0.02 (8%)<br>Overall[b]: 0.85 ± 0.07 |
| RT-229 | - | Fast[a]: 4.08± 0.01 (100%)<br>Slow[a]: NA<br>Overall[b]: 4.06 ± 0.51 | | Fast[a]: 0.71 ± 0.02 (87%)<br>Slow[a]: 0.16 ± 0.03 (13%)<br>Overall[b]: 0.67 ± 0.08 |
| | + | Fast[a]: 41.5 ± 0.83 (7%)<br>Slow[a]: 3.87 ± 0.01 (93%)<br>Overall[b]: 4.53 ± 1.01 | | Fast[a]: 0.79 ± 0.02 (89%)<br>Slow[a]: 0.12 ± 0.02 (11%)<br>Overall[b]: 0.71 ± 0.07 |
| RT-196 | - | Fast[a]: 20.7 ± 0.38 (9%)<br>Slow[a]: 5.22 ± 0.007 (91%)<br>Overall[b]: 6.52 ± 0.59 | | Fast[a]: 0.89 ± 0.03 (84%)<br>Slow[a]: 0.15 ± 0.02 (16%)<br>Overall[b]: 0.77 ± 0.05 |
| | + | Fast[a]: 21.7 ± 0.29 (15%)<br>Slow[a]: 4.83 ± 0.009 (85%)<br>Overall[b]: 6.97 ± 1.37 | | Fast[a]: 0.87 ± 0.03 (84%)<br>Slow[a]: 0.15 ± 0.02 (16%)<br>Overall[b]: 0.70 ± 0.04 |
| PEC-225 | - | Fast[a]: 1.83 ± 0.02 (46%)<br>Slow[a]: 0.55 ± 0.003 (54%)<br>Overall[c]: 1.14 | | Fast[a]: 0.25 ± 0.01 (93%)<br>Slow[a]: 0.05 ± 0.001 (7%)<br>Overall[c]: 0.24 |
| | + | Fast[a]: 1.70 ± 0.01 (42%)<br>Slow[a]: 0.58 ± 0.003 (58%)<br>Overall[c]: 1.05 | | Fast[a]: 0.25 ± 0.01 (93%)<br>Slow[a]: 0.05 ± 0.005 (7%)<br>Overall[c]: 0.23 |
| RT-229 | - | Fast[a]: 2.20 ± 0.01 (41%)<br>Slow[a]: 0.18 ± 0.01 (59%)<br>Overall[b]: 0.94 ± 0.11 | | Fast[a]: 0.26 ± 0.01 (87%)<br>Slow[a]: 0.04 ± 0.002 (13%)<br>Overall[b]: 0.29 ± 0.14 |
| | + | Fast[a]: 2.30 ± 0.01 (41%)<br>Slow[a]: 0.22 ± 0.01 (59%)<br>Overall[b]: 0.99 ± 0.11 | | Fast[a]: 0.31 ± 0.01 (80%)<br>Slow[a]: 0.04 ± 0.002 (20%)<br>Overall[b]: 0.27 ± 0.06 |

[a]Values were calculated from single- or double-exponential fits of the pooled data from all experiments under a given condition. The percentages indicate the contribution of each phase to the overall rate constant. The reported error is the standard deviation of the fits. In the case of a single-exponential fit, the second rate constant is not applicable, NA.

[b]Values represent the average ± the standard deviation (SD) of the mean from three independent replicates.

[c]Values represent the average of the mean from two independent experiments.

**Table S6: Oligonucleotides for TECprobe-VL experiments.** The table below shows oligonucleotides used to perform TECprobe-VL experiments in this study. The modification codes are derived from the Integrated DNA Technologies ordering form, in particular:

/5bioSG/: "standard" 5' biotin

/5Phos/: 5' phosphate

/3AmMO/: 3' amino modifier

| **ID** | **Name** | **Sequence** | **Purification** |
|---|---|---|---|
| TECD006 | PRA1_NoMod.F | TTATCAAAAAGAGTATTGACTCTTTTACCTCTGGCGGTGATAATGGTTGCAT | HPLC |
| TECD017 | dRP1_NoMod.R | AATGATACGGCGACCACCGAGATCTACACGTTCAGAGTTCTACAGTCCGACGATC | HPLC |
| STD002 | HP4_5bio.R | /5Biosg/AATGTCTTCCAGCACACATCGCCTGACGAATCA | none |
| TECP001 | 9N_VRA3 | /5Phos/rNrNrNrNrNrNNNNGATCGTCGGACTGTAGAACTCTGAAC/3AmMO/ | HPLC |
| TECP002 | SC1Brdg_MINUS | CCTTGGCACCCGAGAATTCCAYYYRRATGGCCTTCGGGCCAA | HPLC |
| TECP003 | SC1Brdg_PLUS | CCTTGGCACCCGAGAATTCCARRRYYATGGCCTTCGGGCCAA | HPLC |
| TECP006 | dRP1_5bio.R | /5Biosg/AATGATACGGCGACCACCGAGATCTACACGTTCAGAGTTCTACAGTCCGACGATC | HPLC |
| RPIX_SC1 | RPIX_SC1_Bridge | CCTTGGCACCCGAGAATTCCAATGGCCTTCGGGCCAA | none |
| RPIX | RPIX | CAAGCAGAAGACGGCATACGAGATCGTGATGTGACTGGAGTTCCTTGGCACCCGAGAATTCCA | none |
| RPI | RPI | CAAGCAGAAGACGGCATACGAGAT[Index]GTGACTGGAGTTCCTTGGCACCCGAGAATTCCA | PAGE |

**Table S7. TECprobe-VL DNA templates prepared for this study.** The table below describes the DNA templates prepared for TECprobe-VL experiments, including the primers and templates used, DNA modifications, how the DNA template was purified, and the figures in which each DNA template was used.

| ID | Fwd Primer | Rev Primer | Template | Modifications | Clean up | Used in Fig(s) |
|---|---|---|---|---|---|---|
| 1 | TECD006 | STD002 | pCES019 (WT) | 5’ biotin | Gel extracted | N/A |
| 2 | TECD006 | STD002 | pST001 (U81G, U179G) | 5’ biotin | Gel extracted | N/A |
| 3 | TECD006 | STD002 | pST002 (U81G) | 5’ biotin | Gel extracted | N/A |
| 4 | TECD006 | STD002 | pST003 (U179G) | 5’ biotin | Gel extracted | N/A |
| 5 | TECD006 | STD002 | pST004 (A76G) | 5’ biotin | Gel extracted | N/A |
| 6 | TECD006 | STD002 | pST005 (A75U, A76G) | 5’ biotin | Gel extracted | N/A |
| 7 | TECD006 | STD002 | pST006 (ΔP0) | 5’ biotin | Gel extracted | N/A |
| 8 | TECD006 | STD002 | Template 1 (WT) | Internal biotin-11 nucleotides, 5’ biotin | SPRI beads | 3, 4, S7, S8, S19, S22, S23, S24, S25, S26 |
| 9 | TECD006 | STD002 | Template 2 (U81G, U179G) | Internal biotin-11 nucleotides, 5’ biotin | SPRI beads | S9, S10 |
| 10 | TECD006 | STD002 | Template 3 (U81G) | Internal biotin-11 nucleotides, 5’ biotin | SPRI beads | S11, S12 |
| 11 | TECD006 | STD002 | Template 4 (U179G) | Internal biotin-11 nucleotides, 5’ biotin | SPRI beads | S13, S14 |
| 12 | TECD006 | STD002 | Template 5 (A76G) | Internal biotin-11 nucleotides, 5’ biotin | SPRI beads | 4, S15, S16 |
| 13 | TECD006 | STD002 | Template 6 (A75U, A76G) | Internal biotin-11 nucleotides, 5’ biotin | SPRI beads | S17, S18 |
| 14 | TECD006 | STD002 | Template 7 (ΔP0) | Internal biotin-11 nucleotides, 5’ biotin | SPRI beads | 4, S19, S20, S21 |

**Supplementary Table 8. TECprobe-VL DNA template sequences.** The table below contains the DNA sequences used to perform TECprobe-VL experiments in this study.

| ID | Name | Sequence |
|---|---|---|
| 1 | PRA1_SC1_BsuGly_WT_HP4 | ttatcaaaaagagtattgactcttttacctctggcggtgataatggttgcatggccttcgggccaagaaaatatgagcgaatgacagcaaggg gagagacctgaccgaaaacctcgggatacaggcgccgaaggagcaaactgcggagtgaatctctcaggcaaaagaactcttgctcgac gcaactctggagagtgtttgtgcggatgcgcaaaccaccaaaggggacgtctttgcgtatgcaaagtaaactttcaggtgccaggacagag aaccttcattttacatgaggtgtttctctgtcctttttttgtatcctgattcgtcaggcgatgtgtgctggaagacatt |
| 2 | PRA1_SC1_BsuGly_U81G_U179G_HP4 | ttatcaaaaagagtattgactcttttacctctggcggtgataatggttgcatggccttcgggccaagaaaatatgagcgaatgacagcaaggg gagagacctgaccgaaaacctcgggatacaggcgccgaaggagcaaactgcggagtgaatctcgcaggcaaaagaactcttgctcgac gcaactctggagagtgtttgtgcggatgcgcaaaccaccaaaggggacgtctttgcgtatgcaaagtaaacttgcaggtgccaggacagag aaccttcattttacatgaggtgtttctctgtcctttttttgtatcctgattcgtcaggcgatgtgtgctggaagacatt |
| 3 | PRA1_SC1_BsuGly_U81G_HP4 | ttatcaaaaagagtattgactcttttacctctggcggtgataatggttgcatggccttcgggccaagaaaatatgagcgaatgacagcaaggg gagagacctgaccgaaaacctcgggatacaggcgccgaaggagcaaactgcggagtgaatctcgcaggcaaaagaactcttgctcgac gcaactctggagagtgtttgtgcggatgcgcaaaccaccaaaggggacgtctttgcgtatgcaaagtaaactttcaggtgccaggacagag aaccttcattttacatgaggtgtttctctgtcctttttttgtatcctgattcgtcaggcgatgtgtgctggaagacatt |
| 4 | PRA1_SC1_BsuGly_U179G_HP4 | ttatcaaaaagagtattgactcttttacctctggcggtgataatggttgcatggccttcgggccaagaaaatatgagcgaatgacagcaaggg gagagacctgaccgaaaacctcgggatacaggcgccgaaggagcaaactgcggagtgaatctctcaggcaaaagaactcttgctcgac gcaactctggagagtgtttgtgcggatgcgcaaaccaccaaaggggacgtctttgcgtatgcaaagtaaacttgcaggtgccaggacagag aaccttcattttacatgaggtgtttctctgtcctttttttgtatcctgattcgtcaggcgatgtgtgctggaagacatt |
| 5 | PRA1_SC1_BsuGly_A76G_HP4 | ttatcaaaaagagtattgactcttttacctctggcggtgataatggttgcatggccttcgggccaagaaaatatgagcgaatgacagcaaggg gagagacctgaccgaaaacctcgggatacaggcgccgaaggagcaaactgcggagtgagtctctcaggcaaaagaactcttgctcgac gcaactctggagagtgtttgtgcggatgcgcaaaccaccaaaggggacgtctttgcgtatgcaaagtaaactttcaggtgccaggacagag aaccttcattttacatgaggtgtttctctgtcctttttttgtatcctgattcgtcaggcgatgtgtgctggaagacatt |
| 6 | PRA1_SC1_BsuGly_A75U_A76G_HP4 | ttatcaaaaagagtattgactcttttacctctggcggtgataatggttgcatggccttcgggccaagaaaatatgagcgaatgacagcaaggg gagagacctgaccgaaaacctcgggatacaggcgccgaaggagcaaactgcggagtgtgtctctcaggcaaaagaactcttgctcgacg caactctggagagtgtttgtgcggatgcgcaaaccaccaaaggggacgtctttgcgtatgcaaagtaaactttcaggtgccaggacagaga accttcattttacatgaggtgtttctctgtcctttttttgtatcctgattcgtcaggcgatgtgtgctggaagacatt |
| 7 | PRA1_SC1_BsuGly_deltaP0_HP4 | ttatcaaaaagagtattgactcttttacctctggcggtgataatggttgcatggccttcgggccaaagcaaggggagagacctgaccgaaaac ctcgggatacaggcgccgaaggagcaaactgcggagtgaatctctcaggcaaaagaactcttgctcgacgcaactctggagagtgtttgtg cggatgcgcaaaccaccaaaggggacgtctttgcgtatgcaaagtaaactttcaggtgccaggacagagaaccttcattttacatgaggtgtt tctctgtcctttttttgtatcctgattcgtcaggcgatgtgtgctggaagacatt |

**Table S9 Sequence Read Archive (SRA) deposition table.** All primary sequencing data generated in this work are freely available from the Sequence Read Archive (http://www.ncbi.nlm.nih.gov/sra), accessible via the BioProject accession number PRJNA938111 or using the individual accession numbers below.

| Accession | Sample Name | Sample Description |
|---|---|---|
| SAMN33434193 | Gly_DMS_0mM_rep1 | wt glycine riboswitch multilength cotranscriptional DMS probing with 0mM glycine, replicate 1 |
| SAMN33434194 | Gly_DMS_0mM_rep2 | wt glycine riboswitch multilength cotranscriptional DMS probing with 0mM glycine, replicate 2 |
| SAMN33434195 | Gly_DMS_1mM_rep1 | wt glycine riboswitch multilength cotranscriptional DMS probing with 1mM glycine, replicate 1 |
| SAMN33434196 | Gly_DMS_1mM_rep2 | wt glycine riboswitch multilength cotranscriptional DMS probing with 1mM glycine, replicate 2 |
| SAMN48083625 | GlyU81G_U179G_DMS_0mM_Rep1 | glycine riboswitch U81G_U179G variant multilength cotranscriptional DMS probing with 0mM glycine, replicate 1 |
| SAMN48083626 | GlyU81G_U179G_DMS_0mM_Rep2 | glycine riboswitch U81G_U179G variant multilength cotranscriptional DMS probing with 0mM glycine, replicate 2 |
| SAMN48083627 | GlyU81G_U179G_DMS_1mM_Rep1 | glycine riboswitch U81G_U179G variant multilength cotranscriptional DMS probing with 1mM glycine, replicate 1 |
| SAMN48083628 | GlyU81G_U179G_DMS_1mM_Rep2 | glycine riboswitch U81G_U179G variant multilength cotranscriptional DMS probing with 1mM glycine, replicate 2 |
| SAMN33482618 | GlyU81G_DMS_0mM_Rep1 | glycine riboswitch U81G variant multilength cotranscriptional DMS probing with 0mM glycine, replicate 1 |
| SAMN33482619 | GlyU81G_DMS_0mM_Rep2 | glycine riboswitch U81G variant multilength cotranscriptional DMS probing with 0mM glycine, replicate 2 |
| SAMN33482620 | GlyU81G_DMS_1mM_Rep1 | glycine riboswitch U81G variant multilength cotranscriptional DMS probing with 1mM glycine, replicate 1 |
| SAMN33482621 | GlyU81G_DMS_1mM_Rep2 | glycine riboswitch U81G variant multilength cotranscriptional DMS probing with 1mM glycine, replicate 2 |
| SAMN33482622 | GlyU179G_DMS_0mM_Rep1 | glycine riboswitch U179G variant multilength cotranscriptional DMS probing with 0mM glycine, replicate 1 |
| SAMN33482623 | GlyU179G_DMS_0mM_Rep2 | glycine riboswitch U179G variant multilength cotranscriptional DMS probing with 0mM glycine, replicate 2 |
| SAMN33482624 | GlyU179G_DMS_1mM_Rep1 | glycine riboswitch U179G variant multilength cotranscriptional DMS probing with 1mM glycine, replicate 1 |
| SAMN33482625 | GlyU179G_DMS_1mM_Rep2 | glycine riboswitch U179G variant multilength cotranscriptional DMS probing with 1mM glycine, replicate 2 |
| SAMN34441076 | GlyA76G_DMS_0mM_Rep1 | glycine riboswitch A76G variant multilength cotranscriptional DMS probing with 0mM glycine, replicate 1 |
| SAMN34441077 | GlyA76G_DMS_0mM_Rep2 | glycine riboswitch A76G variant multilength cotranscriptional DMS probing with 0mM glycine, replicate 2 |
| SAMN34441078 | GlyA76G_DMS_1mM_Rep2 | glycine riboswitch A76G variant multilength cotranscriptional DMS probing with 1mM glycine, replicate 2 |
| SAMN34441079 | GlyA76G_DMS_1mM_Rep4 | glycine riboswitch A76G variant multilength cotranscriptional DMS probing with 1mM glycine, replicate 4 |
| SAMN48083629 | GlyA75U_A76G_DMS_0mM_Rep1 | glycine riboswitch A75U_A76G variant multilength cotranscriptional DMS probing with 0mM glycine, replicate 1 |
| SAMN48083630 | GlyA75U_A76G_DMS_0mM_Rep2 | glycine riboswitch A75U_A76G variant multilength cotranscriptional DMS probing with 0mM glycine, replicate 2 |
| SAMN48083631 | GlyA75U_A76G_DMS_1mM_Rep1 | glycine riboswitch A75U_A76G variant multilength cotranscriptional DMS probing with 1mM glycine, replicate 1 |
| SAMN48083632 | GlyA75U_A76G_DMS_1mM_Rep2 | glycine riboswitch A75U_A76G variant multilength cotranscriptional DMS probing with 1mM glycine, replicate 2 |
| SAMN48083633 | GlyDeltaP0_DMS_0mM_Rep1 | glycine riboswitch deltaP0 variant multilength cotranscriptional DMS probing with 0mM glycine, replicate 1 |
| SAMN48083634 | GlyDeltaP0_DMS_0mM_Rep2 | glycine riboswitch deltaP0 variant multilength cotranscriptional DMS probing with 0mM glycine, replicate 2 |
| SAMN48083635 | GlyDeltaP0_DMS_1mM_Rep1 | glycine riboswitch deltaP0 variant multilength cotranscriptional DMS probing with 1mM glycine, replicate 1 |
| SAMN48083636 | GlyDeltaP0_DMS_1mM_Rep2 | glycine riboswitch deltaP0 variant multilength cotranscriptional DMS probing with 1mM glycine, replicate 2 |

**Table S10. RMDB data deposition table.** Reactivity data generated in this work are freely available from the RNA Mapping Database (RMDB, http://rmdb.stanford.edu), accessible using the RMDB ID numbers indicated.

| RMDB Accession | RNA | Experiment | Folding | Probe | Ligand | NTP Conc. | Merged | Smoothing |
|---|---|---|---|---|---|---|---|---|
| BSUGLY_DMS_0001 | wt glycine riboswitch | TECprobe-VL | Cotranscriptional | DMS | none | 50uM | Yes | Yes |
| BSUGLY_DMS_0002 | wt glycine riboswitch | TECprobe-VL | Cotranscriptional | DMS | 1mM glycine | 50uM | Yes | Yes |
| BSUGLY_DMS_0003 | glycine riboswitch, U81G, U179G | TECprobe-VL | Cotranscriptional | DMS | none | 50uM | Yes | Yes |
| BSUGLY_DMS_0004 | glycine riboswitch, U81G, U179G | TECprobe-VL | Cotranscriptional | DMS | 1mM glycine | 50uM | Yes | Yes |
| BSUGLY_DMS_0005 | glycine riboswitch, U81G | TECprobe-VL | Cotranscriptional | DMS | none | 50uM | Yes | Yes |
| BSUGLY_DMS_0006 | glycine riboswitch, U81G | TECprobe-VL | Cotranscriptional | DMS | 1mM glycine | 50uM | Yes | Yes |
| BSUGLY_DMS_0007 | glycine riboswitch, U179G | TECprobe-VL | Cotranscriptional | DMS | none | 50uM | Yes | Yes |
| BSUGLY_DMS_0008 | glycine riboswitch, U179G | TECprobe-VL | Cotranscriptional | DMS | 1mM glycine | 50uM | Yes | Yes |
| BSUGLY_DMS_0009 | glycine riboswitch, A76G | TECprobe-VL | Cotranscriptional | DMS | none | 50uM | Yes | Yes |
| BSUGLY_DMS_0010 | glycine riboswitch, A76G | TECprobe-VL | Cotranscriptional | DMS | 1mM glycine | 50uM | Yes | Yes |
| BSUGLY_DMS_0011 | glycine riboswitch, A75U, A76G | TECprobe-VL | Cotranscriptional | DMS | none | 50uM | Yes | Yes |
| BSUGLY_DMS_0012 | glycine riboswitch, A75U, A76G | TECprobe-VL | Cotranscriptional | DMS | 1mM glycine | 50uM | Yes | Yes |
| BSUGLY_DMS_0013 | glycine riboswitch, delta P0 | TECprobe-VL | Cotranscriptional | DMS | none | 50uM | Yes | Yes |
| BSUGLY_DMS_0014 | glycine riboswitch, delta P0 | TECprobe-VL | Cotranscriptional | DMS | 1mM glycine | 50uM | Yes | Yes |
